# Structures of influenza A and B replication complexes explain avian to human host adaption and reveal a role of ANP32 as an electrostatic chaperone for the apo-polymerase

**DOI:** 10.1101/2024.04.20.590211

**Authors:** Benoit Arragain, Tim Krischuns, Martin Pelosse, Petra Drncova, Martin Blackledge, Nadia Naffakh, Stephen Cusack

**Affiliations:** European Molecular Biology Laboratory, 71 Avenue des Martyrs, CS 90181, 38042 Grenoble Cedex 9, France; RNA Biology of Influenza Virus, Institut Pasteur, Université Paris Cité, CNRS UMR 3569, Paris, France; Institut de Biologie Structurale, Université Grenoble-Alpes-CEA-CNRS UMR5075, 38000 Grenoble, France

**Keywords:** Influenza virus, negative-strand RNA virus, virus-host interactions, virus-host adaptation, acidic nuclear protein 32 (ANP32), chaperone, RNA-dependent RNA polymerase, RNA synthesis, replication, electron cryo-microscopy, atomic structure, mass photometry

## Abstract

Replication of influenza viral RNA depends on at least two viral polymerases, a parental replicase and an encapsidase, and cellular factor ANP32. ANP32 comprises an LRR domain and a long C-terminal low complexity acidic region (LCAR). Here we show that ANP32 is recruited to the replication complex (replicase-ANP32-encapsidase) by first acting as an electrostatic chaperone to stabilise the encapsidase moiety within apo-polymerase symmetric dimers that are distinct for influenza A and B polymerases. The encapsidase, with ANP32, then forms an asymmetric complex with the replicase. Cryo-EM structures of the influenza A and B replication complexes give new insight into the mutations known to adapt avian strain polymerases to use the distinct ANP32 in mammalian cells. The cryo-EM map of the FluPolB complex shows extra density attributable to the ANP32 LCAR wrapping around and stabilising the apo-encapsidase conformation. These results suggest a functional requirement for three polymerases for replication.

## Introduction

Influenza polymerase (FluPol) uses the viral genomic RNA (vRNA) as template to perform synthesis of either capped and polyadenylated viral mRNA (transcription) or unmodified progeny genome copies (replication) in the nucleus of the infected cell (Wandzik et al., 2021; Zhu et al., 2023). The functional context for both processes is the viral ribonucleoprotein complex (RNP), a flexible supercoiled rod-shaped particle, in which the vRNA is packaged by multiple copies of the viral nucleoprotein (NP) with one FluPol bound to the conserved 3’ and 5’ ends of the vRNA. Both processes require FluPol to recruit essential host factors. For transcription, FluPol binds to cellular RNA polymerase II (Pol II) to gain access to nascent, capped transcripts from which capped transcription primers are excised, a process known as cap-snatching (Krischuns et al., 2021; Lukarska et al., 2017). In particular, FluPol binding to the serine 5 phosphorylated (pS5) C-terminal domain (CTD) of Pol II is conserved amongst influenza A, B and C polymerases although the binding sites on FluPol are divergent (Krischuns et al., 2022; Lukarska et al., 2017; Serna Martin et al., 2018). It has been recently proposed that the CTD may serve as a platform for both transcription and replication (Krischuns et al., 2024), but additionally for replication, the highly conserved acidic nuclear protein 32 (ANP32) is an obligatory host factor (Long et al., 2016; Sugiyama et al., 2015). ANP32 comprises a folded, N-terminal leucine-rich repeat (LRR) domain followed by a Glu-, Asp-rich intrinsically disordered region known as the low complexity acidic region (LCAR). ANP32 proteins have multiple cellular functions, notably as histone chaperones (Yu et al., 2022). Of the three functional ANP32 isoforms in human cells, hANP32A and hANP32B support human adapted influenza A and B virus replication (Long et al., 2019; Park et al., 2020; Zhang et al., 2019; Zhang et al., 2020), but not hANP32E (Sheppard et al., 2023). ANP32 is required for both vRNA to cRNA and cRNA to vRNA replication (Nilsson-Payant et al., 2022; Swann et al., 2023). It is thought that ANP32 plays at least two mechanistic roles. First, it stabilises the formation of an asymmetric FluPol dimer comprising a replicase, which is part of an RNP and synthesises the genome copy, and an encapsidase, a newly synthesised apo-FluPol, which binds the 5’ end of the nascent replicate to nucleate formation of a progeny RNP. Second, ANP32 is proposed to recruit successive NPs via a direct interaction between the LCAR and NP thus facilitating co-replicational packaging of product RNA into a progeny RNP (Camacho-Zarco et al., 2023; Wang et al., 2022). Extensive biochemical and mutagenesis studies have shown that hANP32A binds to FluPol (Camacho-Zarco et al., 2020; Mistry et al., 2019) and NP (Camacho-Zarco et al., 2023; Wang et al., 2022). Moreover, the cryogenic electron microscopy (cryo-EM) structure of the putative influenza C replication complex, comprising the replicase-encapsidase dimer bound to ANP32 has been determined (Carrique et al., 2020). These data underlie the proposed model, but there are a number of aspects of the replication mechanism that remain unclear. Firstly, the structure of the replication complex (i.e. replicase-encapsidase dimer bound to ANP32) has not been determined for influenza A or B viruses, for which most of the biochemical and molecular virological data have been obtained. In the case of influenza A, an avian specific 33 residue insertion in avANP32A compared to hANP32A is critical to explain why avian adapted influenza A strain polymerases cannot replicate in human cells (Long et al., 2016). Indeed, avian to human inter-species transmission necessitates adaptive mutations in the avian polymerase (typically PB2/E627K, D701N or Q591R, see below) to be able to productively use the mammalian ANP32 for replication (Peacock et al., 2023). A complete molecular understanding behind these intriguing observations is still lacking. Given that the binding sites of Pol II CTD on influenza A, B and C polymerases are significantly different, it is likely that there has been co-evolution in the mode of ANP32 binding since the divergence of influenza A, B and C. Therefore, it is important to characterise structurally the influenza A and B replication complexes.

As a step towards further understanding of the role of ANP32 in replication, we first analyse binding of hANP32A to FluPolA and FluPolB and show that, at least *in vitro*, it acts as an electrostatic chaperone (Huang et al., 2021) to solubilise the apo-polymerase at physiological salt concentrations, with distinct roles for the LRR and LCAR domains. In the case of FluPolB, cryo-EM shows that hANP32A stabilises, at low salt, a previously undescribed apo-dimer with a 2-fold symmetrical interface, quite different from that of FluPolA (Chang et al., 2015; Fan et al., 2019; Kouba et al., 2023), with one monomer being preferentially in the encapsidase configuration. Cryo-EM structures also show that hANP32A is an integral part of the asymmetric replication complex for both FluPolA strain A/Zhejiang/DTID-ZJU01/2013(H7N9) (A/H7N9) and FluPolB strain B/Memphis/2003. These structures (summarised in Table 1) reveal that there are significant differences in the contacts between the replicase, encapsidase and hANP32A and in domain orientations, compared to the FluPolC complex. In the FluPolB replication complex, additional density clearly indicates the trajectory of the LCAR wrapping around the encapsidase. We also provide minigenome data that combined with extensive existing data in the literature validate these structures as functionally relevant. Importantly, they also provide a rationale for many of the observed mutations that favour adaption of avian FluPol to mammalian cells and that have eluded explanation for many years. Finally, we propose a generalised trimer model of replication, whereby an ANP32-stabilised incoming apo-dimer (distinct for FluPolA and FluPolB) interacts with a replication competent vRNP or cRNP to form the functional replication complex.

**Table 1.** Summary of FluPolA and FluPolB polymerase structures.

| Structure No. | Short name | Resolution | PDB/EMDB |
| --- | --- | --- | --- |
| **FluPolA-4M monomers** |  |  |  |
| 1 | Core 1 with ENDO(R) | 2.77 Å | PDB 8RMP, EMD-19366 |
| 2 | Core 2 with ENDO(R) | 2.54 Å | PDB 8RMQ, EMD-19367 |
| **FluPolA-4M replication complex with hANP32A** |  |  |  |
| 3 | Focus Replicase | 3.21 Å | PDB 8RMS, EMD-19369 |
| 4 | Focus Encapsidase + 627(R) + hANP32A | 3.13 Å | PDB 8RN0, EMD-19382- |
| 5 | Complete Replication Complex | 3.25 Å | PDB 8RMR, EMD-19368 |
| **FluPolB monomers** |  |  |  |
| 6 | Apo-FluPolB encapsidase | 2.89 Å | PDB 8RN2, EMD-19384 |
| 7 | FluPolB with 5' cRNA | 3.64 Å | PDB 8RN1, EMD-19383 |
| **FluPolB symmetrical dimers** |  |  |  |
| 9 | Focus Encapsidase moiety | 2.75 Å | PDB 8RN3, EMD-19385- |
| 10 | Focus ENDO(T) moiety | 2.87 Å | PDB 8RN4, EMD-19386 |
| 11 | Focus ENDO(R) moiety | 2.88 Å | PDB 8RN5, EMD-19387 |
| 12 | Focus ENDO(E) moiety | 2.82 Å | PDB 8RN6, EMD-19388 |
| 13 | Focus core moiety | 3.09 Å | PDB 8RN7, EMD-19389 |
| 14 | Complete dimer | 2.92 Å | PDB 8RN8, EMD-19390 |
| **FluPolB trimer with hANP32A** |  |  |  |
| 15 | Focus Replicase | 3.31 Å | PDB 8RN9, EMD-19391- |
| 16 | Focus Encapsidase+ hANP32A | 3.13 Å | PDB 8RNB, EMD-19393 |
| 17 | Focus Replication Complex | 3.52 Å | PDB 8RNC, EMD-19394 |
| 18 | Complete Trimer | 3.57 Å | PDB 8RNA, EMD-19392 |

### *In vitro*, ANP32 is as an electrostatic chaperone for apo-influenza polymerase

Recently it has been shown that proteins DAXX and ANP32 act as ‘electrostatic’ chaperones that exhibit disaggregase activity dependent on extensive polyD/E (Asp/Glu) stretches within their sequences (Huang et al., 2021). Such chaperones bind to basic peptides on the target protein and have a maximal effect between 25-150 mM salt, declining in activity between 150-300 mM, indicative of electrostatic interactions. To investigate complex formation between recombinant FluPol and ANP32 we analysed by gel filtration and mass photometry(Foley et al., 2021) mixtures of apo-FluPol A/H7N9 or B/Memphis with hANP32A as a function of NaCl concentration and with different truncations of hANP32A (Figure 1, Figure S1).

**Figure 1.**
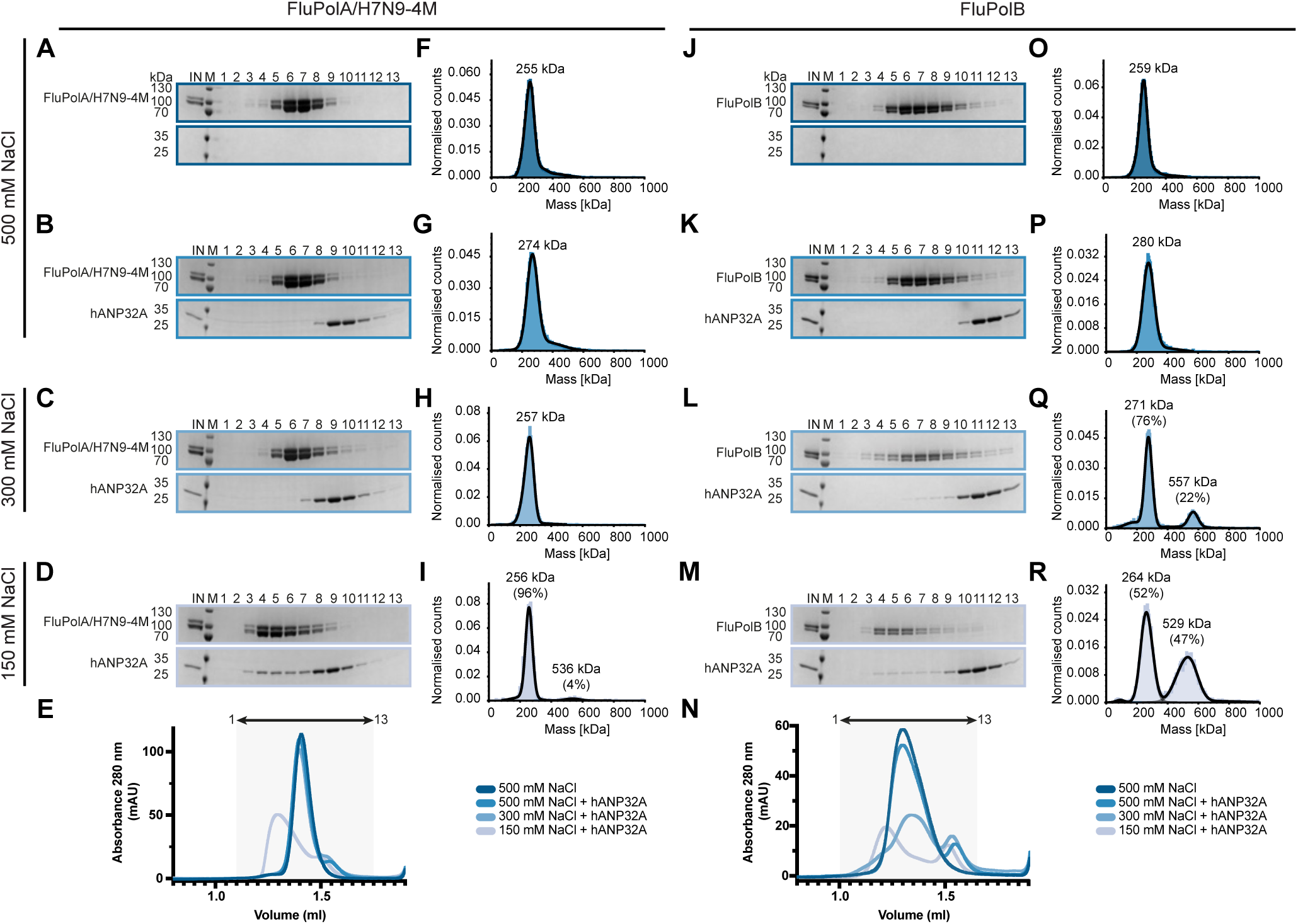
Biochemical analysis of the interaction of FluPolA/H7N9-4M and FluPolB with hANP32A. **(A)** SDS-PAGE analysis of FluPolA/H7N9-4M at 500 mM NaCl. The molecular ladder (M) in kDa and FluPolA/H7N9-4M heterotrimer are indicated on the left of the gel. “IN” corresponds to the input. **(B-D)** SDS-PAGE analysis of FluPolA/H7N9-4M-hANP32A at 500 **(B)**, 300 **(C)**, and 150 mM NaCl **(D)**. The molecular ladder (M) in kDa, FluPolA/H7N9-4M heterotrimer, and hANP32A are indicated on the left of the gel. “IN” corresponds to the input. **(E)** Superposition of size exclusion chromatography profiles of FluPolA/H7N9-4M alone at 500 mM NaCl (dark blue), and with hANP32A at 500 mM (blue), 300 mM (light blue) and 150 mM (blue-grey) NaCl. The relative absorbance at 280 nm (mAU) is on the y-axis. The elution volume (ml) is on the x-axis, graduated every 50 µl. SDS-PAGE fractions 1 to 13 corresponds to the elution volume 1.1 ml - 1.75 ml, represented as an arrow on top. **(F)** Mass photometry analysis of FluPolA/H7N9-4M at 500 mM NaCl. The mass determined in kDa of the monomeric FluPolA/H7N9-4M is indicated. **(G-I)** Mass photometry analysis of FluPolA/H7N9-4M-hANP32A interaction at 500 mM **(G)**, 300 **(H)**, and 150 mM NaCl **(I)**. The determined masses in kDa of the main species are indicated. **(J)** SDS-PAGE analysis of FluPolB at 500 mM NaCl. The molecular ladder (M) in kDa and FluPolB heterotrimer are indicated on the left of the gel. “IN” corresponds to the input. **(K-M)** SDS-PAGE analysis of FluPolB-hANP32A interaction at 500 **(K)**, 300 **(L)**, and 150 mM NaCl **(M)**. The molecular ladder (M) in kDa, FluPolB heterotrimer, and hANP32A are indicated on the left of the gel. “IN” corresponds to the input. **(N)** Superposition of size exclusion chromatography profiles of FluPolB alone at 500 mM NaCl (dark blue), and with hANP32A at 500 mM (blue), 300 mM (light blue) and 150 mM (blue-grey) NaCl. The relative absorbance at 280 nm (mAU) is on the y-axis. The elution volume (ml) is on the x-axis, graduated every 50 µl. SDS-PAGE fractions 1 to 13 corresponds to the elution volume 1.0 ml - 1.65 ml, represented as an arrow on top. **(O)** Mass photometry analysis of FluPolB at 500 mM NaCl. The determined mass in kDa of the monomeric FluPolB is indicated. **(P-R)** Mass photometry analysis of FluPolB-hANP32A interaction at 500 **(P)**, 300 **(Q)**, and 150 mM NaCl **(R)**. The determined masses in kDa of the main species are indicated.

Apo-FluPolA forms homo-dimers with a 2-fold symmetrical interface mediated by loops from the cores of each the three subunits (Chang et al., 2015; Fan et al., 2019; Kouba et al., 2023). The peripheral domains (PA endonuclease, PB2-C) remain flexible in cryo-EM structures (Fan et al., 2019; Kouba et al., 2023), but take up the replicase conformation when constrained in a crystal (Fan et al., 2019). It has been proposed that template realignment following internal initiation of cRNA to vRNA replication would be specifically facilitated by transient trimer formation involving a third apo-polymerase interacting via using the symmetrical dimer interface with the replicase moiety of the replication complex (Carrique et al., 2020; Chen et al., 2019; Fan et al., 2019). However, it has recently been shown that loop mutations that eliminate symmetrical dimerisation of FluPolA are not detrimental to polymerase activity in the minigenome assay (Krischuns et al., 2024) and indeed are selected for when virus evolves to use the usually non-permissive hANP32E, in the absence of hANP32A and hANP32B (Sheppard et al., 2023). It was concluded from the latter work that optimal virus replication requires the correct balance between competing symmetric and asymmetric polymerase dimer formation, consistent with previous results (Chen et al., 2019). To characterise structurally the FluA asymmetric replication complex, without interference from the symmetric dimer, we have used in the following analysis the monomeric A/H7N9-4M mutant (PA E349K, PA R490I, PB1 K577G, and PB2 G74R). This mutant was demonstrated to be active *in vitro* and in cells in previous studies aimed at elucidating the role of the Pol II CTD in replication (Krischuns et al., 2024).

Full-length (FL, 1-249) hANP32A is unable to bind apo-polymerase A/H7N9-4M at 500 or 300 mM NaCl, but can do so at 150 mM NaCl, resulting in a broadening and a shift in the elution profile (Figure 1A-E). Complimentary mass photometry, which however is performed at lower concentration (nM versus µM range), shows that the FluPol A/H7N9-4M remains mainly monomeric at all salt concentrations, but with a small fraction of polymerase dimers at 150 mM NaCl (Figure 1F-I). At 150 mM NaCl before gel filtration, FluPol A/H7N9-4M precipitates in the presence of the hANP32A LRR domain alone (1-149) (Figure S1B), but mass photometry, at low concentration, shows the presence of some polymerase dimer (Figure S1G). With the LRR together with half the LCAR (1-199) at 150 mM NaCl, binding, solubilisation and a small fraction of dimers is observed as for the FL hANP32A but the gel filtration profile does not shift (Figure S1C,E,H). With the LCAR alone (149-Cter) at 150 mM NaCl, binding and a shifted elution profile similar to FL hANP32A is observed, but no dimers are detected in mass photometry (Figure S1D,E,I). These results show that only below 300 mM NaCl can hANP32A bind and solubilise A/H7N9-4M polymerase and this depends on the presence of at least half the LCAR. Dimer formation requires the LRR, the LCAR alone being insufficient. Binding to FluPol of the full LCAR (either alone or with the LRR) is mainly responsible for the broadened and shifted gel filtration profile.

These results show that FL hANP32A has two functions with respect to FluPol A/H7N9-4M, firstly to act as an electrostatic chaperone preventing aggregation of apo-FluPol at physiological salt concentrations, this requiring the LCAR. Secondly, hANP32A promotes dimer formation, this requiring at least the LRR domain, however FluPolA/H7N9-4M dimers are rare under the conditions used. Below we show that these dimers correspond to the asymmetric replication complex, consistent with the fact that the monomeric FluPolA/H7N9-4M mutant does not form symmetric dimers (unlike the wild-type, see below).

For wild-type (WT) FluPol B/Memphis complementary results are obtained. Notably, whereas the polymerase is monomeric at 500 mM NaCl, it forms equally monomers and a stable dimer at 150 mM NaCl in the presence of hANP32A (with already some dimer detected at 300 mM NaCl), as indicated by mass photometry and a gel filtration profile shift (Figure 1J-R). Similar results are obtained with the LCAR alone (Figure S1M,R,N). At 150 mM NaCl with only the LRR domain, the polymerase precipitates in gel filtration but mass photometry at low concentration detects a small fraction (7%) of dimers (Figure S1K,P,N). With the LRR plus half the LCAR (1-199) partial solubilisation is achieved and only 15% dimers are formed (Figure S1L,Q,N). These results show that, as for FluPolA, the LCAR is required for apo-FluB polymerase solubilisation at physiological salt concentration, supporting the notion that hANP32A is acting as an electrostatic chaperone. Interestingly, the stable FluPolB dimer detected by mass photometry only requires the LCAR (Figure S1M,R,N). Below we show by cryo-EM that this dimer is a novel symmetric FluPolB dimer, not previously described. We also observe in cryo-EM a very small proportion of FluPolB trimers which comprise the asymmetric, hANP32A-bound replication complex with the replicase making a symmetric dimer with a third polymerase. These particles are likely too rare to be detected in the biochemical and biophysical experiments, although the FluPolB dimer with the LRR alone could correspond to the asymmetric dimer.

As previously established (Fan et al., 2019; Kouba et al., 2023) wild-type A/H7N9 FluPol forms symmetric homodimers at high salt (Figure S1S, V). However, these dimers are only stable at low salt in the presence of bound hANP32A (Figure S1T, V). On the other hand, promoter bound FluPolA is mainly monomeric at low salt even in the absence of hANP32A (Figure S1U, V). These results are consistent with the requirement of hANP32A to chaperone apo-FluPolA at physiological salt concentrations, at least *in vitro*.

### ANP32 binding to apo-influenza A and B polymerases promotes formation of the replication complex

To characterise structurally the influenza A asymmetric replication complex we analysed complexes of FluPol A/H7N9-4M with hANP32A by cryo-EM (Supp. Info. 1-2, Table 1, Table S1). The majority of particles observed are monomers of the polymerase core with the endonuclease in the replicase conformation (ENDO(R)). Two distinct such structures were determined at 2.77 (CORE-ENDO(R)-1) and 2.54 (CORE-ENDO(R)-2) Å resolution respectively, differing in the degree of polymerase core opening (Supp. Info. 1, Table S1). Consistent with the biochemical and biophysical analysis, a small fraction of particles correspond to the FluA replication complex, comprising a replicase and an encapsidase bridged by hANP32A, whose overall structure was determined at 3.25 Å resolution. The overall map resolution is limited by the flexibility between the replicase and hANP32A-encapsidase moieties as well as the presence of a preferred orientation. To alleviate this, a small number of particles have been selected to equilibrate the distribution of orientations. Focussed refinement on the separate replicase and hANP32A-encapsidase moieties then yield good quality maps of respectively 3.21 and 3.13 Å resolution allowing a relatively complete model to be built (Supp. Info. 2; Table S1). The replicase core is most similar to that in the CORE-ENDO(R)-1 structure. In the case of FluPolB, cryo-EM grids made after mixing apo-polymerase and hANP32A show a majority of dimers with 2-fold symmetric interface, which are quite distinct from those of FluPolA (Supp. Info. 3-4, Table 1, Table S2). Several different pseudo-symmetric dimer structures were determined at 2.8 to 3.1 Å overall resolution, with invariably an encapsidase paired with a variable partner (Supp. Info. 3). A minority of particles are FluPolB trimers, comprising the asymmetric replication complex with an additional polymerase, whose overall structure was determined to 3.6 Å resolution. Further refinement focussed on the replicase or hANP32A-encapsidase moieties improved the map quality and estimated resolution to 3.3 Å (Supp. Info. 5; Table 1; Table S3). The FluPolB replication complex is similar to that of FluPolA, with the replicase moiety forming a FluPolB type symmetric dimer with a third polymerase (see below).

### Structure of the FluPol A/H7N9-4M replication complex

The FluPolA replication complex comprises the asymmetric replicase-encapsidase dimer with bound hANP32A (Figure 2; Figure S2). Both replicase (R) and encapsidase (E) have the conserved polymerase core (PA-C, PB1, PB2-N), but with differently disposed PA-N and PB2-C peripheral domains (Figure 2A; Figure S2B,C). Compared to the transcription (T) conformation (Figure S2A), the replicase (R) conformation is characterised by a rotated endonuclease, against which packs the PB2 NLS domain with the C-terminal, helical NLS containing peptide extending across the endonuclease surface (Fan et al., 2019; Hengrung et al., 2015; Thierry et al., 2016)(Figure 2A; Figure S2A,B,D). The PB2 cap-binding domain (CBD) is packed against the palm domain of PB1 (Figure S2E). The PB2/627-NLS(R) double domain is in the open conformation (Delaforge et al., 2015) with the linker extended and the otherwise flexibly connected PB2/627(R) domain being held in place in the replication complex by interactions with the PB2/NLS(E) domain (Figure 2A). In the distinct encapsidase (E) conformation (Figure S2C), the endonuclease packs on the PB1 fingers domain, but has only low resolution density. The flexible 51-72 insertion of the endonuclease contacts the top of the CBD(E) (e.g. residues PA/55-57 with PB2/I461, 469-471, K482), which is not rigidly integrated into the complex either (Figure 2A; Figure S2C,F). Interestingly, this PA loop, found in FluPolA and FluPolB but not FluPolC, has been shown to be essential for replication by FluPolA (Nilsson-Payant et al., 2018). In contrast, the PB2(E) midlink domain is stabilised by an antiparallel alignment of strands 520-524 with PB2-N 126-132 (Figure S2C,G). PB2(E)-N residues 138-226, which includes the helical lid domain, are not visible in the map, having apparently been displaced to avoid clashing with the encapsidase endonuclease. There is putative density for the PB1-PB2 helical interface bundle, but no model can be built, contrary to the FluPolB case (see below). The encapsidase 627-NLS double domain is in the closed conformation with the 627-domain packing against PA-C. The NLS(E) domain makes a substantial contact with the replicase 627-domain (Figure 2A).

**Figure 2.**
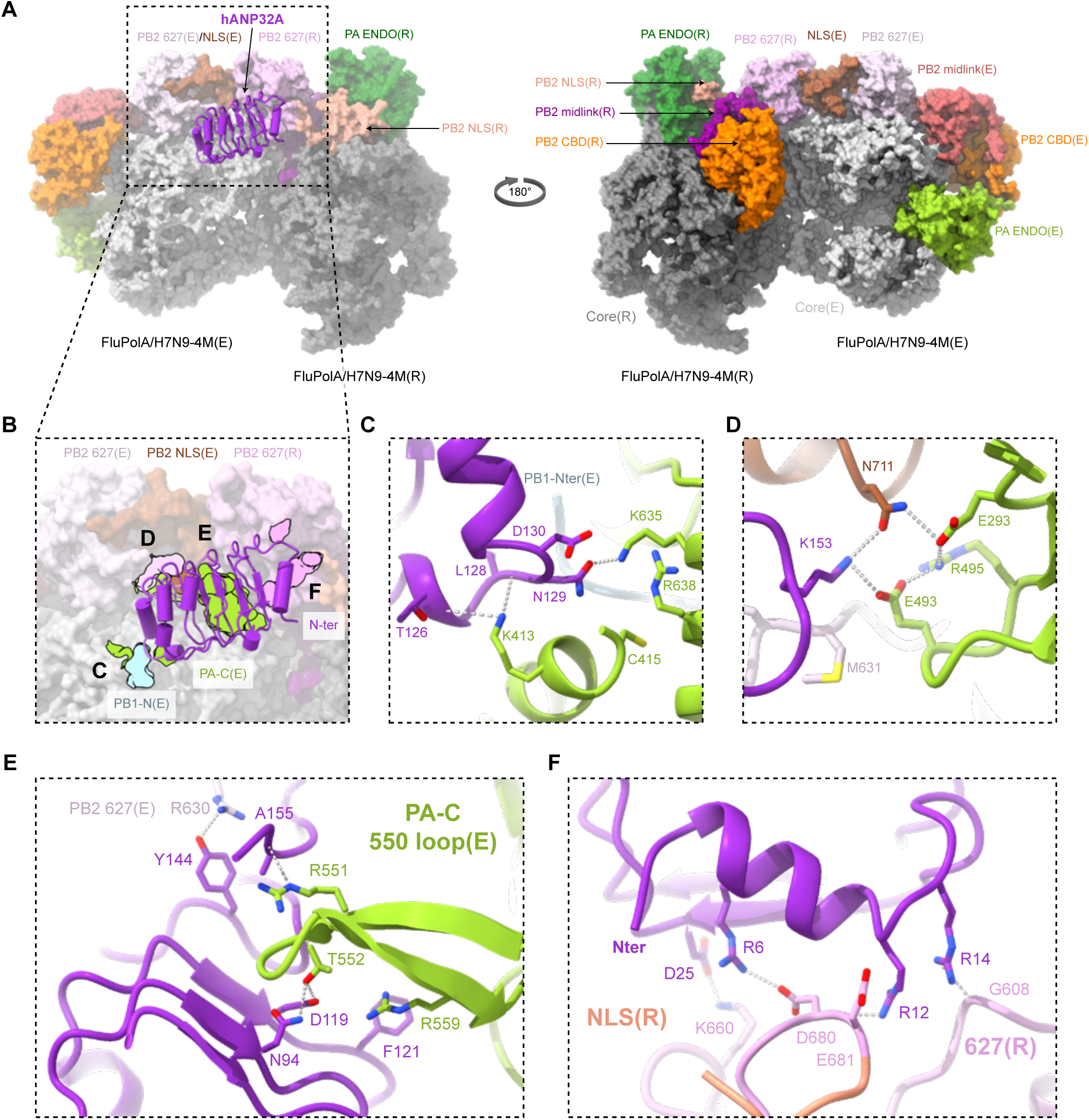
Overall structure of the FluPolA/H7N9-4M replication complex and the interactions of hANP32A. **(A)** Surface representation of the FluPolA/H7N9-4M replication complex with hANP32A displayed as a cartoon (purple). FluPolA/H7N9-4M replicase (R) core is dark grey. PA ENDO(R) is in dark green, PB2 midlink(R) magenta, PB2 CBD(R) orange, PB2 627(R) pink and PB2 NLS(R) beige. FluPolA/H7N9-4M encapsidase (E) core is light grey. PA ENDO(E) is in light green, PB2 midlink(E) salmon, PB2 CBD(E) orange, PB2 627(E) light pink and PB2 NLS(E) brown. **(B)** Close up view of hANP32A interactions with FluPolA/H7N9-4M replicase complex. Interaction surface are highlighted, main contacts are labelled from **(C)** to **(F)**, and coloured according to FluPolA/H7N9-4M interacting domains. PB1-N(E) is coloured in light blue, PA-C(E) in light green, with PB2 627(E)/NLS(E), PB2 627(R) coloured as in **(A)**. **(C)** Cartoon representation of the interaction between hANP32A 128-130 loop and FluPolA/H7N9-4M PA-C(E) and PB1-N(E). hANP32A and FluPolA/H7N9-4M domains are coloured as in **(B)**. Ionic and hydrogen bonds are shown as grey dotted lines. **(D)** Cartoon representation of the interaction between hANP32A K153 and FluPolA/H7N9-4M PA-C(E) and PB2 627(E)/NLS(E). hANP32A and FluPolA/H7N9-4M domains are coloured as in **(B)**. Ionic and hydrogen bonds are shown as grey dotted lines. **(E)** Cartoon representation of the interaction between hANP32A curved β-sheet and FluPolA/H7N9-4M PA-C(E) 550-loop. hANP32A and FluPolA/H7N9-4M domains are coloured as in **(B)**. Ionic and hydrogen bonds are shown as grey dotted lines. **(F)** Cartoon representation of the interaction between hANP32A N-terminus and FluPolA/H7N9-4M PB2 627(R)/NLS(R). hANP32A and FluPolA/H7N9-4M domains are coloured as in **(B)**. Ionic and hydrogen bonds are shown as grey dotted lines.

The interface between the replicase and the encapsidase buries a solvent accessible surface of around 3300 Å^2^, with three main zones of contact (Figure S3). The first involves PB2-N(R) β-strands (128-134, 243-250), neighbouring PA(R) 432-438 loop interacting with the PA(E) arch (N-ter side, 368-377) and the tip of the PB1(E) β-hairpin (361-364) (Figure S3A,B). The latter region is close to the encapsidase 5’ hook binding site, which however is empty in this apo-structure. Key hydrophobic contacts are made by PA(E)/I330, W368 (which changes rotamer) and M374 to PB2-N(R)/T129, M243 and T245; PB1(E)/M362 to PA(R)/P434 and I438 and PB1(E)/K363 to PB2-N(R)/F130 (Figure S3B). The impact of various mutations designed to disrupt this interface was tested using cell-based assays for the A/WSN/33 FluPol in a vRNP reconstitution assay with a reporter vRNA to assess overall transcription/replication activity and a split luciferase-based complementation assay to assess binding to hANP32A (Figure S3E). FluPol activity was significantly impaired in the presence of the PA I330A and PB2 T129A-T245A mutations, more markedly so when they were combined (Figure S3C), consistent with the described structure. Similar trends were observed when FluPol activity support by either hANP32A, hANP32B or chANP32A was determined by transient complementation in HEK-293T cells knocked out for hANP32A and hANP32B (Figure S3D). This is consistent with decreased FluPol-binding levels to either hANP32A, hANP32B, or chANP32A, as determined in a split luciferase-based complementation assay (Figure S3E).

The second zone of interaction between replicase and encapsidase involves the C-terminal β-sheet region of PB2-627(R) (residues 645, 651-657, 668-669) with PA(E) (315-316, 550-loop 547-558) (Figure S3A,F). Notable hydrophobic interactions include PB2-627(R) M645 and L668-G669 with PA(E) F315 and PB2-627(R) P654 with PA(E) L549, together with a hydrogen bond between PA(E) Q556 and PB2-627(R) N652 carbonyl oxygen (Figure S3F). The third zone of interaction is localized at the interface of PB2-627(R) (residues 585-587, 631-637 on the 627-loop, 644-646) with PB2-NLS(E) (residues 703-708, 712, 715-720) (Figure S3A,G). Notable hydrophobic contacts are made by PB2-627(R)/A587, M631, F633 S637 and R646 (Figure S3G). Using the cell-based assays described above, we found that the mutations PB2/A587K or A717E significantly reduced overall FluPol activity (Figure S3H), its dependence on either hANP32A, hANP32B or chANP32A (Figure S3I) as well as binding to hANP32A, hANP32B and/or chANP32A (Figure S3J), consistent with the structural findings. The PB2 A717K mutation had the strongest effect on FluPol activity but not on binding to hANP32A, suggesting that it impairs another FluPol function beyond the replicase-encapsidase interaction.

PB2-627(R) behaves as a rigid-body part of the encapsidase in focussed cryo-EM refinement (Supp. Info. 2), which is explained by its interfaces with the encapsidase PB2-NLS(E) and PA-C(E) domains. The particular juxtaposition of the replicase 627 domain with the encapsidase 627-NLS domains is a key feature that distinguishes both the FluPolA and B replication complexes from the previously described FluPolC complex (Figure 3). In FluPolA, the NLS(E) domain is sandwiched between the 627(E) and 627(R)-domains, with no contact between the latter two domains, whereas in FluPolC the 627-NLS(E) double domain is rotated relative to the 627(R) domain by ∼ 78° and the NLS(E) domain squeezed to one side (Figure 3A, B). Consequently, in FluPolC, the replicase and encapsidase 627-domain loops containing K649 (equivalent to E627 in FluPolA/H7N9-4M and K627 in FluPolB) are closer together and face each other with the two K649 Cα atoms ∼19 Å apart. The LCAR is proposed to pass over this interface (Carrique et al., 2020). In FluPolA/B these two loops are rotated far apart from each other with a main chain distance of respectively ∼40 Å and ∼42 Å between equivalent PB2/627 residues (Figure 3A,C). This makes a significant difference to the surface with which the LCAR of ANP32 is likely to interact in FluPolA and FluPolB (see below).

**Figure 3.**
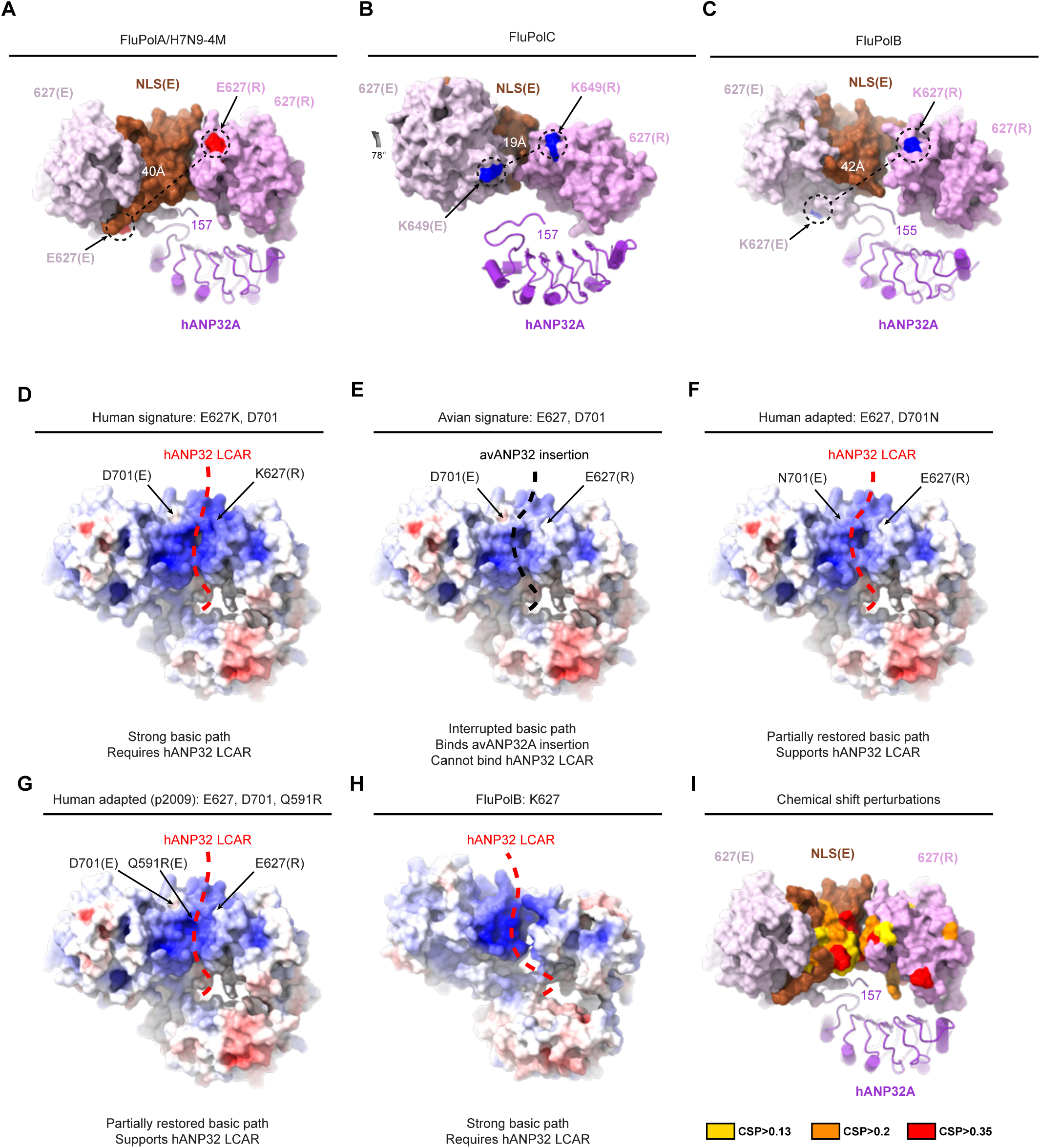
hANP32A, PB2 627-NLS(E) and PB2 627(R) domain organisation and implication in human adaptive mutations. **(A)** Surface representation of FluPolA/H7N9-4M PB2 627(E)/NLS(E) and PB2 627(R) domains. hANP32A is displayed as cartoon. PB2 E627 residue is coloured in red as surface. The distance between PB2 E627(R) and PB2 E627(E) is indicated. The last seen hANP32A C-terminal residue is annotated. hANP32A and FluPolA/H7N9-4M domains are coloured as in Figure 2. **(B)** Surface representation of FluPolC PB2 627(E)/NLS(E) and PB2 627(R) domains. FluPolA/H7N9-4M PB2 627(R) **(A)** has been used as reference to align the corresponding FluPolC PB2 627(R) domain (extracted from PDB 6XZQ). hANP32A is displayed as cartoon. PB2 K649 residue is coloured in blue as surface. The distance between PB2 K649(R) and PB2 K649(E) is indicated. FluPolC PB2 627(E)/NLS(E) 78 degree rotation compared to FluPolA/H7N9-4M PB2 627(E)/NLS(E) is indicated with an arrow. The last seen hANP32A C-terminal residue is annotated. hANP32A and FluPolC domains are coloured as FluPolA/H7N9-4M. **(C)** Surface representation of FluPolB PB2 627(E)/NLS(E) and PB2 627(R) domains. FluPolA/H7N9-4M PB2 627(R) **(A)** has been used as reference to align the corresponding FluPolB PB2 627(R) domain. hANP32A is displayed as cartoon. PB2 K627 residue is coloured in blue as surface. The distance between PB2 K627(R) and PB2 K627(E) is indicated. The last seen hANP32A C-terminal residue is annotated. hANP32A and FluPolB domains are coloured as FluPolA/H7N9-4M. **(D-G)** Surface representation of FluPolA/H7N9 627-NLS(E) and 627(R) domains, bearing human or avian adapted mutations, coloured according to the electrostatic potential: **(D)** human signature with PB2 E627K/D701 (modelled); **(E)** avian signature with PB2 E627/D701 (modelled); **(F)** human adapted avian signature with PB2 E627/D701N (obtained in this study); **(G)** human adapted (p2009) signature with PB2 E627/D701/Q591R (modelled). A strong basic path allows hANP32 LCAR (red dotted line) electrostatic interaction to FluPolA/H7N9 PB2 627(E)/NLS(E) and PB2 627(R) domains. An interrupted basic path allows avANP32A insertion (black dotted line) interaction, displayed as a black dotted line. **(H)** Surface representation of FluPolB PB2 627(E)/NLS(E) and PB2 627(R) domains coloured by electrostatic potential. The putative path of the hANP32 LCAR is shown as a red dotted line. **(I)** Chemical shift perturbations (CSPs), calculated from chemical shift differences between free and hANP32A bound forms of 627(K)-NLS from avian H5N1 A/duck/Shantou/4610/2003, were taken from Figure 2 of (Camacho-Zarco et al., 2020). CSPs were mapped onto the structure of the 627-NLS(E) and 627(R) domains from the FluPolA/H7N9-4M replication complex. CSPs from the 627 domain were designated as being associated with the replicase, while shifts from the NLS domain were associated with the encapsidase. Red corresponds to the highest CSPs (CSP>0.35), orange corresponds to intermediate CSPs (>0.2) and yellow to lower but still measurable CSPs (CSP>0.13).

### Interactions of hANP32A with the FluPolA encapsidase and replicase

Binding of the hANP32A LRR domain into the replicase-encapsidase dimer buries ∼3300 Å^2^ of solvent accessible surface of which 80 % is with the encapsidase (Figure 2B). The C-terminal end of the LRR domain, notably the 128-129 loop, packs against the encapsidase PA-C domain, burying the N-terminus of PB1 and the peptide 152-157 curves back against the LRR domain to contact the 627-NLS domains of the encapsidase (Figure 2B). In particular, hANP32A/N129 makes a key interaction with PA(E)/K635, a residue previously shown to be critical for the binding of a phosphoserine in Pol II CTD binding site 1 of both FluPolA and B (Krischuns et al., 2024; Lukarska et al., 2017). In addition, PA(E)/K413 makes multiple hydrogen bonds to the main-chain of residues 126-128 of hANP32A (Figure 2C). Consistently, when cell-based mutational analysis was performed in the A/WSN/33 FluPol background, the PA/K413A, PA/K413E and PA/K635A mutations reduced FluPol activity (Figure S4A), FluPol dependence on hANP32A, hANP32B or chANP32A (Figure S4B), and binding to hANP32A, hANP32B or chANP32A (Figure S4C). The effect is most dramatic for PA/K413E which shows FluPol activity and binding levels close to background. These data are in agreement with a previous report that the PA/K413A mutation affects replication of FluPolA, based on the observed role of the equivalent residue K391 in the FluPolC replication complex (Carrique et al., 2020) (note that this residue is not conserved in FluPolB). Similarly, it has been previously reported that the PA/K635A mutant, is not only defective for CTD binding and hence transcription activity, but also replication activity (Krischuns et al., 2024; Lukarska et al., 2017). The observation that PA/K635 is important for binding to both the Pol II CTD and to hANP32A suggests that simultaneous binding of CTD and ANP32 to PA is sterically impossible, notably in the context of the encapsidase. ANP32A/K153 also makes interactions with PB2(E)/N711 and PA(E)/E493, which are stabilised by PA(E)/R495 and E293 in a network of polar interactions (Figure 2D).

To assess the impact of hANP32A mutations on FluPolA activity, vRNP reconstitution were performed in HEK-293T cells knocked out for ANP32A and ANP32B, and transiently complemented with a wild-type or mutant hANP32A protein. Compared to the wild-type, the hANP32A mutants K153A and K153E were less efficient in supporting FluPol activity (Figure S4D). The PA/E493K mutant was also impaired in FluPol activity. Interestingly, FluPol activity was partially rescued when charge-reversal mutants PA/E493K and hANP32A/K153E (but not K153A) were combined (Figure S4D), indicating that the interaction is restored to some extent. Consistently, hANP32A/K153E and PA/E493K individually decreased FluPol-binding to hANP32A, but showed increased binding-levels when tested in combination (Figure S4E). This is in line with the interactions shown in Figure 2C,D and with previous results showing that this region of hANP32A is critical for its interactions with the polymerase (Park et al., 2021).

Another major point of contact is of the PA-C(E) 550-loop, which bends to be able to interact with the concave beta-sheet surface of hANP32A. Loop residues R551, T552 (an avian specific residue, (Mehle et al., 2012), normally serine in mammalian polymerases) and R559 make direct interactions with hANP32A/A155, D119, N94 and F121 respectively (Figure 2E). A deletion in the PA 550-loop was previously shown to affect replication in a cell-based assay (Krischuns et al., 2024; Krischuns et al., 2022). Consistent with these observations, we show that the triple mutation R551A-S552A-R559A in WSN PA affects both FluPol activity and binding to hANP32A, as does the double mutation F121A-N122A in hANP32A, although with a relatively modest effect on FluPol activity (Figure S4F-G). Beyond residue 157 there is only disjointed, low resolution density for hANP32A, so that the conformation and interactions of the LCAR cannot be visualised precisely in the FluPolA replication complex.

The interactions of the FluPolA replicase with hANP32A are more tenuous (Figure 2F). K660 from the PB2-627(R) 660-loop makes a salt bridge with hANP32A/D25. Residues 680-DE-681 from the extended replicase NLS-627(R) linker likely make electrostatic interactions with R6 and R12 of hANP32A and G608 from PB2-627(R) with R14, although the density is relatively poor in this region, due to mobility. Again, cell-based mutational analysis (PB2/K660A, hANP32A/D25A) confirmed the structural findings (Figure S4H-I). Importantly, steady-state levels of the WT and mutant PA, PB2 and hANP32A proteins used for functional studies in cell-based assays were similar as determined by western blot (Figure S4J-L).

### Correspondence with other published studies on FluPol-ANP32 interactions

There is already abundant literature on putative interactions between hANP32A and FluPolA. Residues 129-130 have been shown to be critical for defining functional (or non-functional) species and isoform specific interactions of ANP32. In mammals, these residues are generally 129-ND in ANP32A and ANP32B (but are NA and SD in mouse, ANP32A and B, respectively), all of which support influenza A replication, although mouse proteins are suboptimal (Peacock et al., 2020; Zhang et al., 2019). Avian ANP32A has 129-ND and supports influenza replication, whereas avANP32B (129-IN) and avANP32A with the single N129I mutation do not (Long et al., 2019). Human or avian ANP32E, with 129-ED, poorly support replication (Idoko-Akoh et al., 2023; Long et al., 2019; Sheppard et al., 2023). These observations are fully consistent with the FluPolA replication complex structure that shows N129 interacting with PA/K635(E) (Figure 2C) and which can be plausibly substituted by the smaller serine, as in mouse, but not by the larger isoleucine or glutamate. Virus that has evolved to use ANP32E in human cells knocked out for ANP32A and ANP32B acquires the PB1/K577E and PA/Q556R (550-loop) mutations (Sheppard et al., 2023). The PB1 mutation likely acts by weakening the competing symmetric dimer interface as proposed (Sheppard et al., 2023)(compare FluPolA/H7N9-4M, which bears the PB1-K577G substitution), whereas our replicase structure shows that the PA/Q556R mutation could make a strong salt-bridge with hANP32A E154 (which is conserved in ANP32E), thus promoting asymmetric dimer formation (Figure S5A,B). In a related experiment, when virus is selected to replicate in transgenic chickens or chicken embryos carrying mutant chANP32A (N129I, D130N) instead of WT chANP32A, escape mutants PA/E349K, Q556R, T639I, G634E, K635E, K635Q and PB2/M631L, I570L are found, with a predominance of PA/E349K and PB2/M631L. PA/349K again acts by weakening the symmetric dimer interface (Chen et al., 2019; Sheppard et al., 2023)(compare FluPolA/H7N9-4M, which also bears this substitution), whilst Q556R would strengthen the interaction with ANP32A (see above). The other PA mutations cluster around the key contact with the hANP32A 129-130 loop, plausibly making local perturbations that better accommodate 129-IN-130 (Figure S5A). PB2/M631L is at the other main contact where the polymerase interacts with hANP32A K153 and again may facilitate accommodation to the mutated chANP32A D130N (Figure S5A).

### The FluPolA replication complex structure explains avian to human host adaptations

Adaptation of avian strain polymerases (which invariably have PB2/E627) to be able to function in human cells requires PB2 mutations. Real-life evolution of circulating influenza A viruses and numerous laboratory studies show that the most effective routes to adapt avian polymerase to mammalian ANP32A or ANP32B are PB2/E627K (Subbarao et al., 1993), PB2/D701N (Chen et al., 2013; Gabriel et al., 2005) or PB2/Q591R (A/H1N1pdm09 strain) (Mehle and Doudna, 2009; Peacock et al., 2023), with other observed mutations (PA/T572S, PB2/T271A, K702R, D740N) potentially assisting to some extent.

To understand why these residues make such a difference, we calculated electrostatic surfaces using the FluPolA replication complex model with appropriate modelled substitutions in replicase and encapsidase for the four cases (Figure 3D-G): (1) typical human signature with PB2/Q591, K627, D701 (Figure 3D), (2) typical avian signature with PB2/Q591, E627, D701 (Figure 3E), (3) human adapted A/H7N9 FluPol with PB2/Q591, E627, N701 (corresponding to our structure) (Figure 3F), and (4) human adapted A/H1N1pdm09 FluPol with PB2/R591, E627, D701 (Figure 3G).

The typical human signature results in an uninterrupted positively-charged path following the PB2/NLS(E)-PB2/627(R) interface (and continuing round the back), encompassing K627(R) and skirting the acidic patch due to D701(E)(Figure 3D). The structure of the fully human adapted FluB replication complex (see below) shows a similar strong basic path (Figure 3H). Residues in FluA contributing to this basic surface are PB2-NLS(E)/K702, R703, K718, K721, K738 and R739 and PB2-627(R)/K586, R589, K627 and R630. We propose that this is likely the trajectory followed by the distal part of the complimentary acidic LCAR of hANP32A or hANP32B (e.g. residues 160-AEGYVEGLDDEEEDEDEEEYDEDAQVV-186, red here is mammalian specific), interacting in a predominantly electrostatic, multivalent fashion, since it is not clearly observed in the structure. Indeed, projecting the residues undergoing chemical shifts when hANP32A interacts with isolated 627-NLS domain, onto PB2/NLS(E)-PB2/627(R) rather than the closed PB2/627(E)-NLS(E) conformation, highlights perturbations (due to direct or indirect binding effects) at the NLS(E)-627(R) interface (Camacho-Zarco et al., 2020) (Figure 3I). The typical avian signature results in an interrupted basic track, due to the combined effect of positively charged D701(E) and E627(R) (Figure 3E). This surface is more appropriate to bind the avian ANP32A due to the 33 amino acid insertion (i.e. residues 160-AEGYVEGLDDEEEDEDVLSLVKDRDDK-186, blue here is avian specific), which could place the avian specific hexapeptide 176-VLSLVK (that strongly interacts, according to NMR(Camacho-Zarco et al., 2020)) and two other basic residues in the equivalent region.

Both human adapted signatures, as in A/H7N9 and A/H1N1pdm09 FluPols, partially restore the basic track (Figure 3F,G). Interestingly PB2/Q591R might have a dual mode of action in both replicase and encapsidase, since the simultaneous Q591R(E) mutation could enhance binding of hANP32 to the encapsidase through formation of a salt bridge with hANP32A/D151 (Figure S5C). Furthermore, the ‘third-wave’ mutation PA/N321K in A/H1N1pdm09 FluPol is thought to be an additional adaptation of a swine polymerase to human ANP32 (Elderfield et al., 2014; Peacock et al., 2020). Indeed, the mutation PA(E)/N321K could lead to a salt bridge with PB2(R)/E249 (Figure S5D), thus strengthening the encapsidase-replicase dimer, a mechanism suggested to compensate for a sub-optimal ANP32 interaction (Sheppard et al., 2023).

It has also been shown that mammalian ANP32A and ANP32B proteins preferentially drive different adaptive mutations in avian FluPol, respectively PB2/D701N or PB2/E627K, and this ability maps to the significantly different LCAR (Peacock et al., 2023)(Supp. Info. 6). One possible explanation is that hANP32B is considerably more acidic than hANP32A in the region 176-190, with an insertion of 5 extra acidic residues and substitution of three non-charged residues by acidic residues (Supp. Info. 6). This hyper-acidic stretch of hANP32B may require the more basic track resulting from the E627K mutation to bind in a functional way.

Importantly, the FluA replication complex structure reveals a significant asymmetry in the positioning of the encapsidase PB2/627 and 701 residues and their counterparts in the replicase. Only the 627(R) and 701(E) residues are in the putative pathway of the LCAR. This would suggest that the nature of these residues, whether E/K or D/N, should only exert their influence in human cells via the replicase or encapsidase, respectively. Whereas the relevant position of PB2/701 has not been analysed, several studies have addressed the effect of making the PB2/E627K substitution only in the encapsidase or the replicase (Manz et al., 2012; Nilsson-Payant et al., 2022; Swann et al., 2023). These studies are based, firstly, on a cRNA stabilisation assay involving infection of human 293T cells in the presence of pre-expressed NP and PB2-627E or -627K polymerase with an inactive PB1, in the presence of actinomycin D or cycloheximide to prevent transcription/translation by the incoming vRNPs. The incoming virus thus provides the vRNA to cRNA replicase, whilst the pre-expressed polymerase acts as encapsidase. Results indicate that both incoming avian 627E and 627K viruses produce stable cRNPs in infected cells, whether the pre-expressed PB2 is 627E or 627K (Manz et al., 2012; Nilsson-Payant et al., 2022; Swann et al., 2023). In a second assay, performed in the presence of pre-expressed NP and polymerase with an active PB1 (i.e. a replication assay), Manz *et al*. found that only 627K viruses could produce functional cRNPs in human cells. In contrast, Nilsson-Payant *et al*. and Swan *et al*. found that: (i) cRNPs produced by 627E and 627K viruses can both serve as a template for cRNA to vRNA synthesis, provided that the pre-expressed PB2 (which now is part of the replicase) is 627K; and (ii) the impaired vRNA synthesis when the pre-expressed PB2 is 627E can be restored by pre-expressing chicken ANP32A. Taken together, these observations indicate that in human cells, the PB2-627E polymerase is functional as a replicase to perform vRNA to cRNA synthesis and as an encapsidase, but is impaired as a replicase to co-opt human ANP32 and perform cRNA to vRNA synthesis. This agrees with the structure showing that only the replicase 627 residues is part of the likely LCAR trajectory, however why this restriction only affects cRNA to vRNA replication remains unexplained.

### Structure of the FluPolB symmetrical dimer

The biochemical and biophysical analysis revealed that a stable apo-FluPolB dimer is formed in the presence of hANP32A at physiological salt concentrations (Figure 1). Indeed, the majority of particles in the FluPolB-hANP32A cryo-EM analysis are dimers or monomers, consistent with the biophysical analysis, together with a minority of trimers (Supp. Info. 2-5). The dimers have a 2-fold symmetrical interface, although the peripheral domains of each monomer can be in quite different conformations (Figure 4, Table 1, Table S3, Supp. Info. 3-4). The symmetrical interface involves the PA-arch residues 375-385 of one monomer contacting PA/332-338 and 361-364 of the second monomer, and *vice versa*. In addition the tips of the PB1 β-hairpin of each monomer (residues 360-363, closely associated with the PA-arch), interact with each other across the 2-fold axis (Figure 4A,B). The core dimer interface buries ∼2200 Å^2^ of solvent accessible surface (less than the ∼3600 Å^2^ for the FluPolA homodimer) and involves both hydrophobic (e.g. PA/F335, Y361, W364, I375, M376, V379) and polar and salt-bridge interactions (e.g. PA/D382 with Y361 and K338, E378 with K338 and K358) (Figure 4C). PA peptides 357-372 (notably aromatics PA/Y361 and W364) and 504-513 (including PA/H506, which stacks on nucleotide 11 of the 5’ hook) are refolded in the apo-form compared to their configuration in the 5’ hook bound form of FluPolB (Figure 4D), suggesting that for FluPolB, symmetrical dimer formation and 5’ hook binding are mutually exclusive. Biochemically and biophysically this is found to be the case since FluPolB bound to the vRNA 5’ hook is soluble and monomeric at 150 mM NaCl and, furthermore, does not bind hANP32A (Figure S6). This highlights the fact that hANP32A is only required to chaperone apo-polymerase, even though we see no density corresponding to it in the symmetrical dimer structures. We confirmed these observations by structure determination of a monomeric form of the FluPolB encapsidase bound to nucleotides 1-12 of the 5’ hook of cRNA (Figure 4E, Table 1, Table S4, Supp. Info. 7), which can serve as a model for the vRNA replication product-bound encapsidase. We conclude that 5’ hook binding disassociates the FluPolB symmetric dimer or, conversely, under certain circumstances, formation of a symmetrical dimer could perhaps eject the bound 5’ end (see discussion).

**Figure 4.**
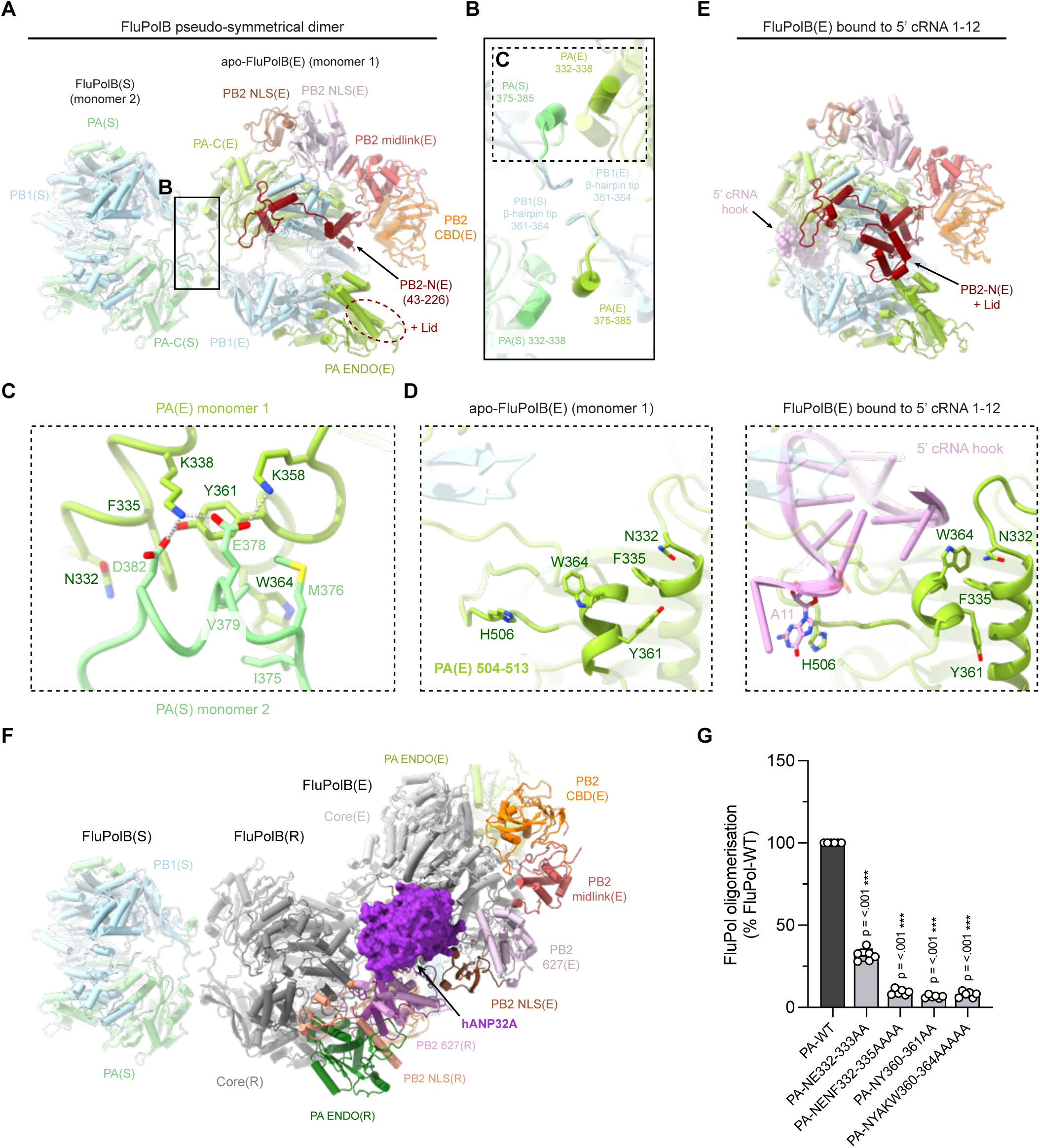
Apo-FluPolB pseudo-symmetrical dimer and 5’ cRNA bound FluPolB encapsidase. **(A)** Cartoon representation of the most abundant apo-FluPolB pseudo-symmetrical dimer. The monomer 1 is in an encapsidase conformation FluPolB(E). PA-C(E) is coloured in light green, PB1(E) in light blue, PB2-N(E) (43-226) in dark red, PB2 midlink(E) in salmon, PB2 CBD(E) in orange, PB2 627(E) in light pink, PB2 NLS(E) in brown. Putative PB2-N(E) lid density are located next to PA-ENDO(E) and indicated by a dotted ellipse. The symmetrical monomer 2 FluPol(S) takes multiple conformation. Here, only PA(S) and PB1(S) subunits are shown, and respectively coloured in light green and light blue. Both monomers core form a symmetrical interface highlighted by a dotted rectangle, with a close-up view shown in **(B)**. **(B)** Close-up view on the FluPolB symmetrical dimer interface. The main interaction is mediated by PA(E) 332-338 interacting with PA(S) 375-385. PB1 β-hairpin tip from both monomers interact with each other across the 2-fold axis. One of the symmetrical dimer interface is highlighted with a dotted rectangle, corresponding to **(C)**. **(C)** Close-up view on the main FluPolB symmetrical dimer interface between PA(E) 332-338 and PA(S) 375-385. Domains are coloured as in **(A-B)**. Ionic and hydrogen bonds are shown as grey dotted lines. **(D)** Structural rearrangement of PA(E) upon 5’ cRNA hook binding. Residues undergoing the biggest movement are displayed. The 5’ cRNA hook (nts 1-12) is coloured in plum with nucleotides represented as stubs. **(E)** Cartoon representation of the 5’ cRNA bound FluPolB encapsidase structure. Domains are coloured as in **(A)**. The 5’ cRNA hook (nts 1-12) is coloured in plum, and atoms are displayed as spheres. The PB2-N(E) lid is observed when FluPol(E) is bound to the 5’ cRNA hook. **(F)** Cartoon representation of the complete FluPolB trimer, composed of the replication complex (FluPolB(R)+hANP32A+FluPolB(E)), with FluPolB(R) core forming a pseudo-symmetrical dimer with a third FluPolB(S). FluPolB(S) is coloured as in **(A)**. hANP32A is displayed as surface, coloured in purple. FluPolB(R) core is coloured in dark grey. PA ENDO(R) in dark green, PB2 midlink(R) in magenta, PB2 CBD(R) in orange, PB2 627(R) in pink, PB2 NLS(R) in beige. FluPolB(E) core is coloured in light grey, PA ENDO(E) in light green, PB2 midlink(E) in salmon, PB2 CBD(E) in orange, PB2 627(E) in light pink, PB2 NLS(E) in brown. **(G)** Cell-based split-luciferase complementation assay to assess B/Memphis/13/2003 FluPol self-oligomerisation for the indicated PA mutants. HEK-293T cells were co-transfected with plasmids encoding PB2, PA, PB1-luc1 and PB1-luc2 (Chen et al., PMID: 31581279). Luminescence signals due to luciferase reconstitution are represented as a percentage of PA-WT (mean ± SD, n=6, ***p < 0.001, one-way ANOVA; Dunnett’s multiple comparisons test).

The cryo-EM analysis shows that most FluPolB symmetric dimers exhibit a complete encapsidase conformation in one monomer (monomer 1 in Figure 4A). In the FluPolB encapsidase, the PB2 lid domain is disordered (as in FluPolA), but interestingly, is observed in its normal position in the 5’ cRNA hook bound form of the encapsidase, due to subtle displacements of domains (compare Figure 4A and 4E). For the symmetric partner monomer (monomer 2 in Figure 4A), denoted FluPolB(S), a variety of conformations are observed including with the endonuclease in either the replicase (ENDO(R)), encapsidase (ENDO(E)) or transcriptase (ENDO(T)) orientations, with the PB2-C domains usually exhibiting only weak density. Even the core conformations vary due to different openings of the polymerase. Given that this encapsidase-containing dimer requires the hANP32A LCAR for stabilisation, we suggest that the LCAR in fact stabilises the encapsidase conformation. This is consistent with the fact that the encapsidase conformation has only ever been visualised in the presence of ANP32, as here, or for FluPolC (Carrique et al., 2020). As shown below, the FluPolB replication complex cryo-EM maps reveal the likely pathway of the extended LCAR, providing a rationale for how it stabilises the encapsidase structure.

We used the split luciferase assay to show that mutations in the FluPolB homodimer interface significantly reduce self-oligomerisation in cells (Figure 4G), suggesting that this dimer likely exists under physiological conditions.

### Structure of the influenza B replication complex

The overall architecture of the influenza B replication complex is similar to that of FluA, with most domains in the equivalent location, but with a few significant differences (Figure 5; Figure S7). In the FluPolB replicase there is a rotation of ∼15° of the ENDO(R) towards the 627(R) domain compared to FluPolA (after alignment of PB1(R)), allowing the endonuclease 63-73 insertion to contact the 627(R) domain in the region of W575 (contact not observed in FluPolA) (Figure S7D). The encapsidase moiety of the FluPolB replication complex adopts a very similar conformation to that seen in the FluPolB symmetrical dimer. The encapsidase endonuclease, ENDO(E) is rotated by ∼48° away from the cap-binding domain compared to FluPolA (after alignment of PB1(E)), so there is no longer contact with the endonuclease 63-73 loop as in FluPolA (Figure S7E). The lid domain of PB2(E) is also disordered in FluPolB encapsidase, although there is some suggestive, but low resolution, density close to the endonuclease. Interestingly, there is unambiguous density for the encapsidase PB1(Cter)-PB2(Nter) helical bundle swung away from its normal position (Figure S7E). This structural element, together with the tip of the encapsidase PB1 β-ribbon (residues 194-198), makes a significant new interface with the top of the replicase cap-binding domain (PB2/466-474), which considerably reinforces the asymmetric dimer (Figure 5A; Figure S8A,F). No equivalent interaction is observed in the FluPolA structure. This extra contact largely accounts for the fact that the total buried surface between encapsidase and replicase in FluB is ∼ 5100 Å^2^ (mainly resulting from the three encapsidase subunits each contacting PB2(R)) considerably more than the ∼3300 Å^2^ for FluA (Figure S8A-D). Interestingly, the PA/609-loop, a specific FluPolB insertion that is important for Pol II CTD binding (Krischuns et al., 2022) makes special contacts within the complex. The encapsidase PA/609 loop interacts with the encapsidase cap-binding domain 420-loop (not shown) and the replicase PA/609 loop interacts with the encapsidase PA arch (Figure S8E).

**Figure 5.**
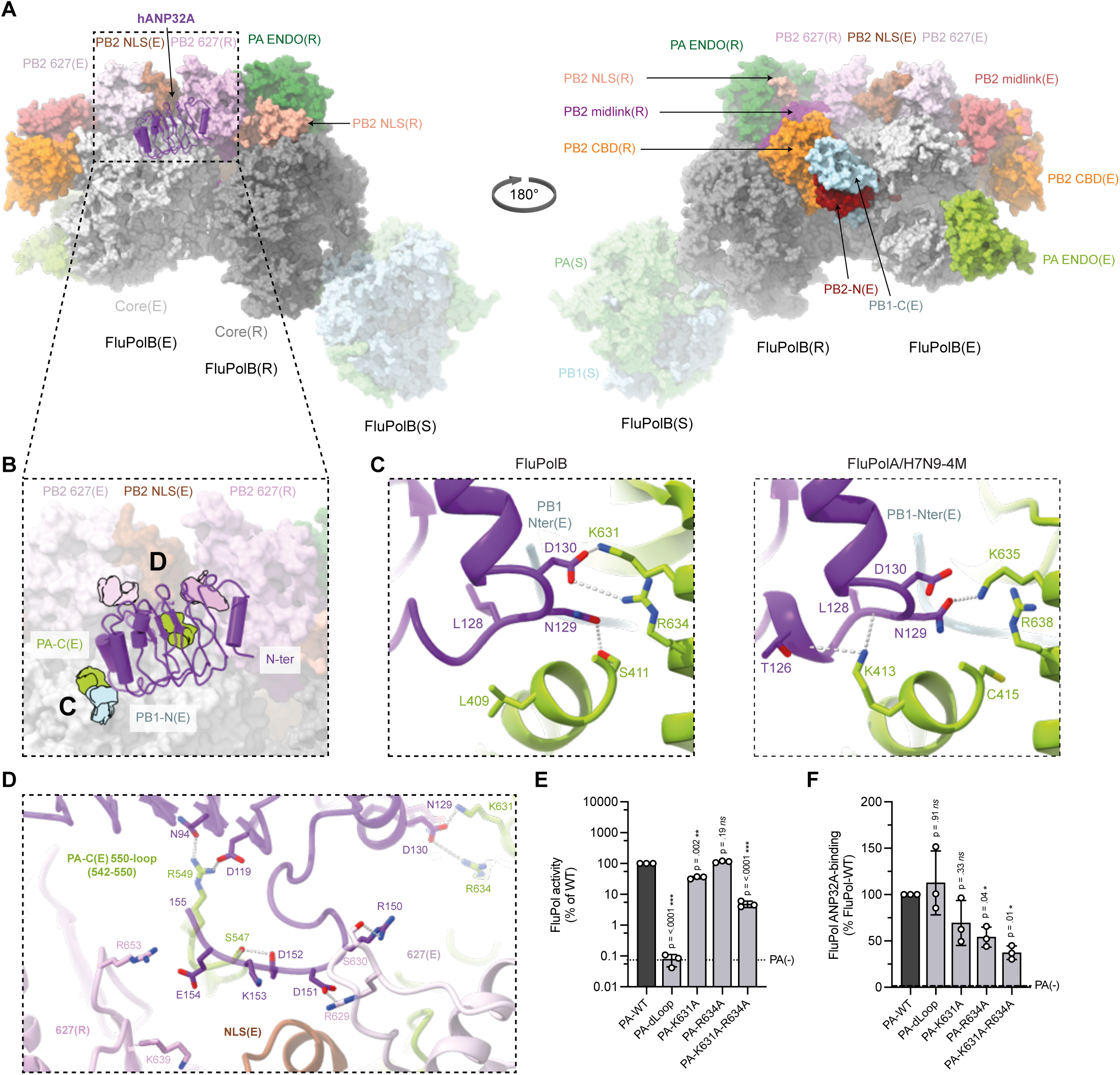
Overall FluPolB trimer replication complex and hANP32A interactions. **(A)** Surface representation of the FluPolB trimer replication complex. hANP32A is displayed as cartoon, coloured in purple. FluPolB replicase (R) core is coloured in dark grey. PA ENDO(R) in dark green, PB2 midlink(R) in magenta, PB2 CBD(R) in orange, PB2 627(R) in pink, PB2 NLS(R) in beige. FluPolB encapsidase (E) core is coloured in light grey, PA ENDO(E) in light green, PB2 midlink(E) in salmon, PB2 CBD(E) in orange, PB2 627(E) in light pink, PB2 NLS(E) in brown. PB1-C(E) and PB2-N(E), respectively coloured in blue and dark red, interacts with PB2 CBD(R) bridging FluPolB(R) and FluPolB(E). The symmetrical FluPolB(S) is coloured as in Figure 4. **(B)** Close up view on hANP32A interaction with FluPolB(R) and FluPolB(E). Interaction surface are highlighted, main contacts are labelled from **(C)** to **(D)**, and coloured according to FluPolB interacting domains. PB1-N(E) is coloured in light blue, PA-C(E) is coloured in light green, PB2 627(E)/NLS(E), PB2 627(R) are coloured as in **(A)**. **(C)** Comparison of the interaction between hANP32A 128-130 loop with FluPolB PA-C(E)/PB1-N(E) and FluPolA/H7N9-4M PA-C(E)/PB1-N(E). hANP32A and FluPol domains are coloured as in **(B)**. Ionic and hydrogen bonding are shown as grey dotted lines. **(D)** Cartoon representation of the interaction between hANP32A curved β-sheet and LRR C-terminus with FluPolB PA-C(E) 550-loop, PB2 627(E)/NLS(E), PB2 627(R). hANP32A and FluPolB domains are coloured as in **(B)**. Ionic and hydrogen bonding are shown as grey dotted lines. **(E)** Cell-based assay of B/Memphis/13/2003 FluPol activity for the indicated PA mutants. HEK-293T cells were co-transfected with plasmids encoding PB2, PB1, PA, NP with a model vRNA encoding the Firefly luciferase. Luminescence was normalised to a transfection control and is represented as a percentage of PA-WT (mean ± SD, n=3, **p < 0.002, ***p < 0.001, one-way ANOVA; Dunnett’s multiple comparisons test). **(F)** Cell-based assay of B/Memphis/13/2003 FluPol binding to ANP32A for the indicated PA mutants. HEK-293T cells were co-transfected with plasmids encoding PB2, PA, PB1-luc1 and hANP32A-luc2. Luminescence signals due to luciferase reconstitution are represented as a percentage of PA-WT (mean ± SD, n=3, *p < 0.033, one-way ANOVA; Dunnett’s multiple comparisons test).

### FluPolB trimer

The FluB replicase-encapsidase complex is only seen as part of a trimer (determined at 3.57 Å resolution overall), with an additional monomer, FluPolB(S), making a symmetrical FluB-type dimer interface with the replicase (Figure 4F, 5A, Table 1, Supp. Info. 5). The third polymerase is less well ordered with only the core visible and not the peripheral domains, likely because it is a mixture of conformations. The encapsidase component of the replication complex cannot simultaneously make a symmetrical dimer as it uses the same interface (the PA-arch and PB1 β-hairpin) to make the asymmetric dimer with the replicase. This shows that the encapsidase component of the symmetric apo-dimer would have to disassociate to be able to form the replication complex. Consistent with this and the biophysical data, a significant number of monomeric apo-encapsidases are observed (Supp. Info. 4). A speculative biological role for the replication complex-containing FluPolB trimer is mentioned in the discussion.

### Interactions of hANP32A within the FluB replication complex

hANP32A binds in the same position and orientation to the FluPolB encapsidase as in FluPolA, but due to sequence divergence the interactions are not necessarily conserved (Figure 5A,B). The overall buried solvent accessible surface upon hANP32A binding to the FluB replication complex is ∼2200 Å^2^ only 66% that for FluPolA. As for FluPolA, the main anchor point remains the C-terminal end of the LRR domain wedged against the PA-C(E) domain in the vicinity of the N-terminus of PB1(E) (Figure 5A,B). On the other hand, the N-terminal end of the LRR domain does not make contact with either polymerase and consequently has less well ordered density due to mobility. A critical contact is again made by hANP32A 129-ND, but in FluB it is D130 that directly interacts with CTD-binding residue PA(E)/K631 (in FluA N129 contacts the equivalent PA/K635) with also a slightly more distant salt-bridge to R634, whereas N129 hydrogen bonds to PA(E)/S411 (Figure 5C). The equivalent of FluPolA PA/K413 is L409 in FluPolB and is close but does not interact (Figure 5C). It has previously been shown that the substitution N129E that occurs in hANP32E is responsible for the limited ability of hANP32E to support FluB replication (Zhang et al., 2020), emphasising the importance of this contact point. Furthermore, we show that mutations PA/K631A and R634A together reduce FluPolB activity (Figure 5E), as previously described (Krischuns, 2022 #2204), as well as hANP32A binding to FluPolB in a cell-based split luciferase assay (Figure 5F). In FluPolA PA(E)/R638 is further away and its mutation less impacts replication (Lukarska et al., 2017). The PA/550-loop of FluPolB is four residues shorter than that of FluA and does not reach so far onto the β-sheet surface of hANP32A (Figure 5B,D), which is consistent with the fact that no decreased binding of FluPolB PA/550-loop to hANP32A was observed (Figure 5F), despite some contacts being observed (PA(E)/S547 to hANP32A D152 and the carbonyl oxygen of K153 and PA(E)/R549 to ANP32A/N84 and D119). Other polar interactions are made to hANP32A/R150 and D151 by PB2(E)/S630 and R629, respectively, and to E154 by PB2(R)/R653 and possibly K639 (Figure 5D).

### Path of the hANP32A LCAR in the FluB replication complex

Additional pseudo-continuous density is present in the FluB replication complex maps that we interpret as tracing the path of the hANP32A LCAR extending beyond residue 155, the last of the folded part of the protein core, and wrapping around the encapsidase (Figure 6A,B). Although no model can be built into this density, it could correspond to an extended chain of at least 50 residues (i.e. beyond hANP32A residue 200). The path follows a clearly defined positively-charged electrostatic track, created by a number of basic residues that would point towards the acidic LCAR, in order along the pathway: PB2-NLS(E)/K721, K742, R741, K703, K734, PB2-627(R)/R629, K588 then PB1(E)/K566, K570 and PA(E)/K298, K301, K475, H506, K374, then PB1(E)/R196 and R203 (close to the PB1 NLS on the long β ribbon (Hutchinson et al., 2011)) and finally PB1(E)/R135, K353 and PB2(E)/R40 (Figure 6C). The LCAR passes over the PB2-NLS(E)/PB2-627(R) interface, then over the 3’ end secondary binding loop (PA/K298, K301), round the tip of the PB1(E) β ribbon (PB1/R203) and parallel to the β ribbon with the PA(E) arch on the other side (PA/K374) (Figure 6B,C). These observations plausibly explain how the LCAR stabilises the encapsidase conformation by electrostatic complementation.

**Figure 6.**
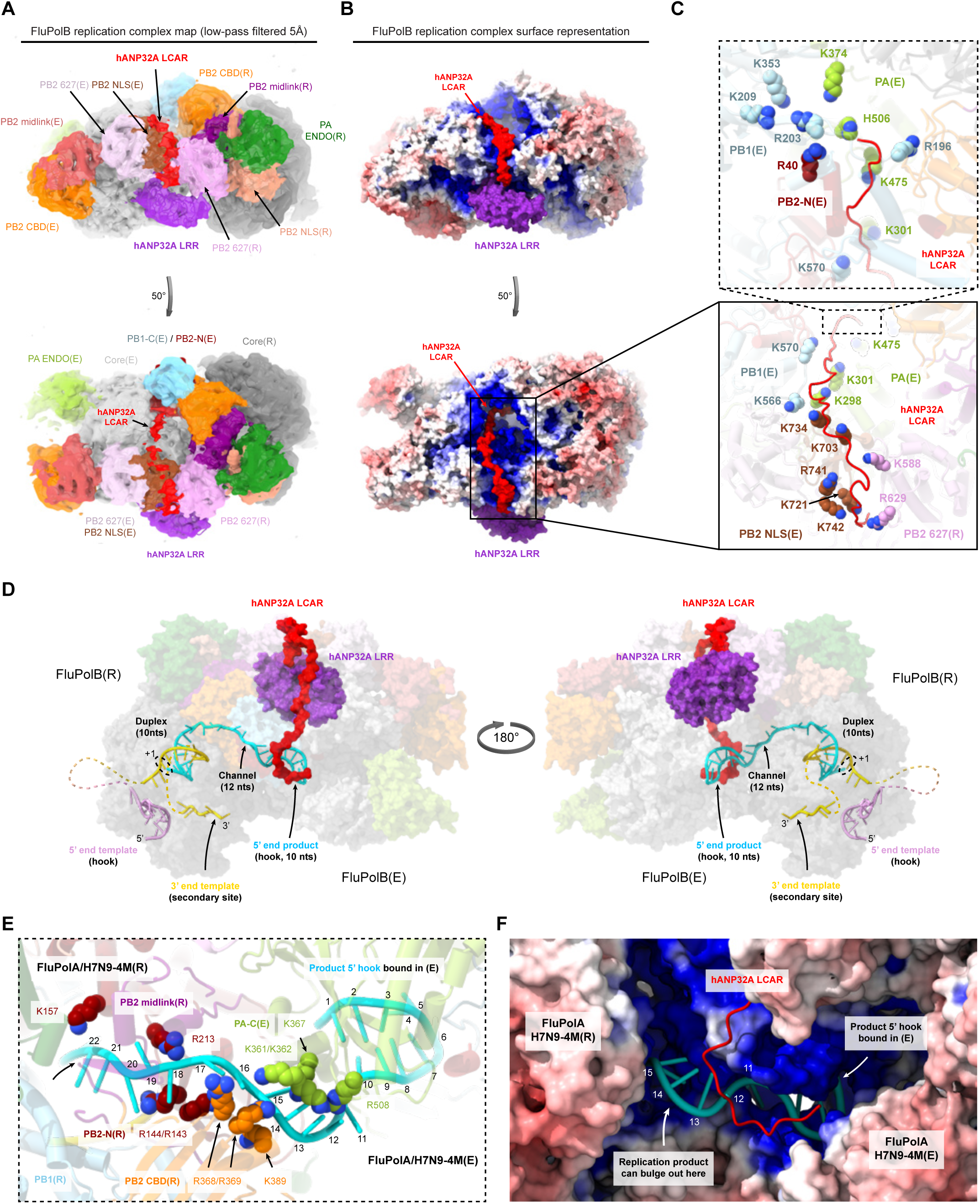
hANP32A LCAR density and putative RNA path within FluPolA/H7N9-4M and FluPolB replication complexes. **(A)** Low-pass filtered FluPolB trimeric replication complex map at 5Å resolution (threshold = 0.12). hANP32A and FluPolB domains are coloured as in Figure 5. Density assigned to the hANP32A LCAR, in red, pass over the NLS(E) and 627(R) interface and extends over FluPolB(E). **(B)** Surface representation of the FluPolB trimer replication complex coloured by electrostatic potential. hANP32A is coloured as in **(A)**. The hANP32A LCAR follows a positively charged electrostatic path. **(C)** Close-up view on the positively-charged electrostatic track followed by hANP32A LCAR. Domains are coloured as in **(A)**. Atoms are displayed as spheres, shown as non-transparent. hANP32A LCAR is displayed as cartoon. **(D)** Model of template and product RNA binding to the FluPolB replication complex in the early-elongation state. The complex is shown as a transparent surface with hANP32A LRR coloured in purple and the LCAR in red. The template 5’ hook is coloured in plum and the 3’ end gold. The replication product is coloured in cyan. The template extremities and product-template duplex bound to the replicase are modelled according to PDB 6QCT. The replication product is modelled to extend through a channel between the replicase PB2-N and midlink/cap-binding domains into the 5’ end hook binding site of the encapsidase (modelled using the cRNA hook-bound FluPol(E) structure). The 3’ end template binds back to the secondary site. Un-modelled RNAs are displayed as dotted lines. The C-terminal end of the modelled LCAR **(A-C)** would clash with the product RNA entering the hook-binding site and therefore would have to be displaced. **(E)** Close-up view of the RNA product exit channel from the replicase to the encapsidase. Domains are coloured as in **(A)**, with PB1(R) in blue, PB2 midlink(R) in magenta, PB2-N(R) in dark red, PB2 CBD(R) in orange, PA-C(E) in green. Modelled RNA product is displayed, coloured as in **(D)**. Nucleotides 1-22 are numbered from the 5’ end to the first nucleotide exiting the duplex in the replicase. Nucleotides 1-10 corresponds to the 5’ hook product, bound to the encapsidase. Nucleotides 11-22 corresponds to the minimal product in the channel formed by PB2-N/midlink/CBD(R) and PA-C(E). Basic residues are displayed with atoms represented as spheres. **(F)** Surface representation of FluPolA/H7N9-4M replication complex coloured according to the electrostatic potential, with the FluPolB modelled RNA product and hANP32A LCAR superimposed. The RNA product and hANP32A LCAR are coloured as in **(E)**. Nucleotides are numbered from the 5’ end. The distal end of the hANP32A LCAR clashes with the RNA product. White arrows indicate the 5’ hook bound to the encapsidase and where the growing replication product could bulge out.

### Model of the RNA bound replication complex

We have determined the structure of the putative FluPolA and B replication complexes in the absence of viral RNA. We therefore sought to model how template and product RNA could bind to an active replication complex (Figure 6D-F). In doing this we have not taken into account the expected conformational changes that are known to accompany promoter binding and the initiation to elongation transition (e.g. different degrees of polymerase opening and extrusion of the priming loop) (Kouba et al., 2019; Wandzik et al., 2020). The template extremities and product-template duplex bound to the replicase were modelled by superposing on PB1 an A/H7N9 elongation state (e.g. PDB 8PNQ). The 3’ end of the template binds back to the secondary site. The replication product was manually extended from the top of the duplex through a channel into the 5’ end hook binding site of the encapsidase (modelled using the cRNA 5’ hook-bound FluPol(E) structure). From the 3’ end at the +1 position in the replicase active site to the 5’ end in the encapsidase hook-binding site, a minimal 32 nucleotides of the product are modelled (10 in the duplex, 12 in the channel, 10 in the 5’ hook) (Figure 6D,E). In the FluPolA structure, the putative product exit channel linking replicase to encapsidase passes between the replicase PB2-N and cap-binding domains and then has PA(E) on one side, while the other side is solvent exposed and could be where the extending product bulges out to be bound by NP (Figure 6E,F). The channel is lined by basic residues notably PB2(R)/R143, R144, K157, R213, R368, R369, K389 and PA(E)/K361, K362, K367, R508 yielding a positively charged passage rich in flexible arginines that are able to interact with both bases and backbone (Figure 6E,F). Not unexpectedly, the product exit channel partially overlaps with the capped transcription primer entrance channel, where it has already been shown how some of the same arginines can adapt to interact with different RNA configurations (Kouba et al., 2023). The RNA model is transferable between the FluPolA and FluPolB replication complexes, without modification. In FluPolB, one can see that the distal, C-terminal end of the modelled LCAR, would clash with the product RNA entering the encapsidase hook-binding site (Figure 6D,F) and therefore would have to be displaced as the RNA emerges. This could have the dual effect of preventing non-specific RNA binding to the encapsidase and releasing the LCAR for interaction with NP when the product RNA emerges.

## Discussion

*De novo* synthesis of the influenza anti-genomic and genomic RNA (cRNA and vRNA), respectively from parental vRNPs and cRNPs, with concomitant packaging of the product RNA into a progeny RNP, is a highly complex process that we are only just beginning to get a grasp off. Two key elements have been established, firstly that the host factor ANP32 plays critical and probably multiple roles in the process, and secondly, that an asymmetric FluPol dimer comprising replicase (integrated into the parental RNP) and encapsidase (a newly synthesised and initially apo-FluPol) is fundamental to nucleate co-replicational assembly of the progeny RNP. The proposed roles of ANP32 so far include formation and stabilisation of the ternary replicase-ANP32-encapsidase replication complex (Carrique et al., 2020) and secondly, through interactions of the LCAR with apo-NP, successive recruitment of NPs to package the growing replicate (Camacho-Zarco et al., 2023; Wang et al., 2022). A further intriguing aspect that needs a full mechanistic explanation is why certain specific mutations, mainly in PB2, are required to overcome the restriction of avian strain polymerases to replicate in human cells, given that this depends on a systematic difference in avian ANP32A (a 33 aa insertion in the LCAR) compared to human ANP32A and ANP32B (Long et al., 2016).

Here we present evidence based on *in vitro* biochemical and structural analysis of complexes of FluPolA and FluPolB with hANP32A, that ANP32 may have a third important role that is to act as an electrostatic chaperone/disaggregase (Huang et al., 2021), that solubilises and stabilises apo-FluPolA or FluPolB in a principally dimeric form at physiological salt concentrations, this depending principally on the LCAR. We speculate that this may have been the primordial role of ANP32 in the nuclear lifecycle of influenza-like viral polymerases, since it is highly conserved in all potential eukaryotic hosts of orthomyxo- and orthomyxo-like viruses. Later, it would have acquired an active role in replication. Interestingly, the FluPolA and FluPolB apo-dimers both have 2-fold symmetric core interfaces (although the peripheral domains need not be disposed symmetrically), but they are structurally quite different. The FluPolA apo-dimer is stable at high salt without ANP32 (Fan et al., 2019; Kouba et al., 2023) but requires ANP32 at physiological salt (Fig S1S-V), whereas FluPolB is monomeric at high salt (Figure 1). One monomer of the FluPolB apo-dimer is observed to be preferentially in the encapsidase conformation, which appears specifically to be stabilised by the LCAR of ANP32. These initial studies led us to perform cryo-EM structural studies on mixtures of apo-FluPolA and FluPolB with hANP32A, which yielded multiple structures including the replication complex of each FluPol. Electrostatic calculations for the FluPolA replication structure suggest that human adaptive mutations restore a coherent positively charged pathway at the NLS(E)- 627(R) interface, able to bind the negatively charged distal LCAR of hANP32A/B, whereas the more mixed surface of avian FluPol more appropriately binds the avian LCAR, which because of the 33 aa insertion has more basic and hydrophobic residues (Figure 3). The FluPolA structure also explains why PB2/E627 is only restrictive in human cells in the replicase. The FluPolB replication complex exhibits density for the extended LCAR following a basic pathway around the encapsidase, thus providing a plausible explanation of how it stabilises this particular FluPol conformation (Figure 6A-C). Both the FluA and FluB replication complexes exhibit a positively-charge channel for the product RNA to exit the replicase and enter the 5’ hook binding site of the encapsidase (Figure 6D-F).

For both FluA and FluB, an ANP32-bound apo-FluPol symmetric dimer is brought together with a replication-competent RNP during the replication initiation process. We therefore propose a generalised trimer model of replication (Figure 7). In each case, one half of the dimer would become the ANP32-bound encapsidase within the asymmetric replication complex. For FluPolA, the third polymerase has been proposed to play a role in template realignment specifically during initiation of cRNA to vRNA synthesis by forming a transient symmetrical interface with the asymmetric replication complex dimer (Fan et al., 2019). However, such a trimer has not been observed structurally. For FluPolB, based on the observed trimer structure, where the third FluPolB forms a different symmetrical interface with the replicase, we speculate that it could have a distinct role. It might assist in replication termination by binding the replicase and releasing the 5’ hook of the template so it can be copied. It is possible that both types of symmetric dimer exist for both FluPolA and B under certain conditions, but this has not been shown yet. However, given that FluPolA mutated to be monomeric appears to be functional (Krischuns et al., 2024; Sheppard et al., 2023), it could be that these third polymerases are not essential but merely increase efficiency of replication.

**Figure 7.**
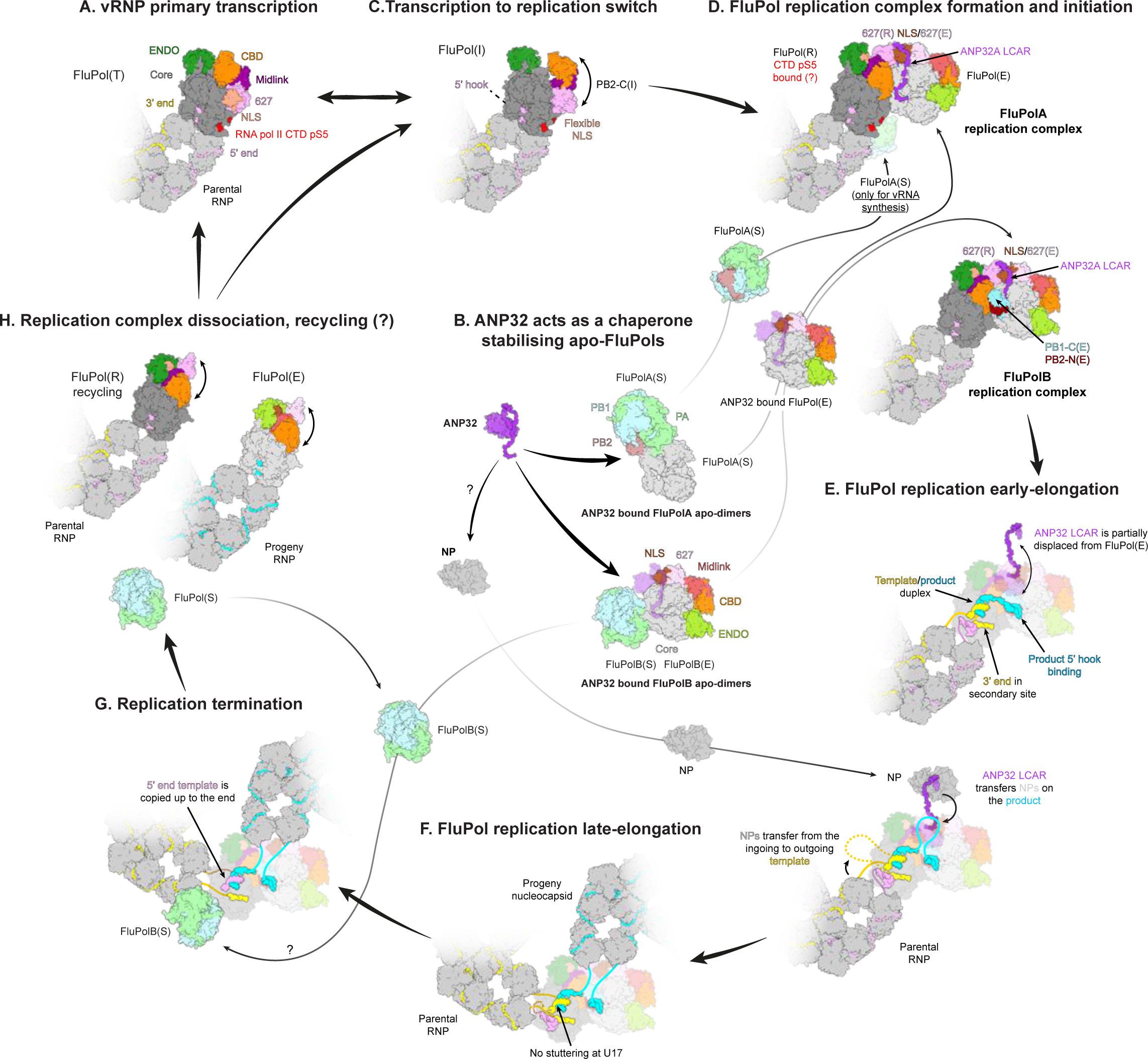
Trimer model of FluPol replication. **(A)** vRNPs bound to cellular Pol II with the FluPol in the transcriptase conformation (FluPol(T)) perform primary transcription leading to the synthesis of capped/poly-adenylated viral mRNAs. These are translated by the host machinery to yield new viral proteins, including the apo-FluPols and NPs that are required for replication. **(B)** ANP32 acts as a chaperone at physiological salt concentration, stabilising apo-FluPol through electrostatic interactions via the LCAR. ANP32-bound apo-FluPolA and apo-FluPolB form distinct dimers with a 2-fold symmetric interface. For FluPolB at least, one of the monomers preferentially takes up the encapsidase conformation (FluPol(E)). **(C)** Within the parental vRNP, due to domain flexibility, the FluPol(T) conformation can transiently adopt the intermediate conformation FluPol(I), but only in the presence of ANP32 bound apo-FluPol, possibly co-localised by interaction with the Pol II CTD, can it be locked into the stable replicase conformation upon formation of the ANP32-FluPol(E)-FluPol(R) complex. The FluPol(E) must derive from dissociation of the symmetric dimer (which is probably in equilibrium with monomeric forms, at least for FluPolB), bringing with it ANP32. While the ANP32 LCAR stabilises FluPol(E), the LRR domain bridges FluPol(E) to FluPol(R). **(D)** Replication initiation may start concomitantly with replication complex formation. For cRNA synthesis, initiation is terminal but for vRNA synthesis, it is internal. For FluPolA, one FluPol from the apo-dimer (FluPol(S)) is proposed to form a symmetric dimer with FluPol(R), allowing template realignment in the case of vRNA synthesis (Fan et al., 2019). A similar mechanism may occur for FluPolB, even though no FluPolA-like symmetric dimers have been described for FluPolB so far. **(E)** FluPol(R) synthesises the complementary replication product. In early elongation, the newly synthetized 5’ end binds to the FluPol(E) hook binding site, accessible from the replicase through a protected channel. Simultaneously, the ANP32 LCAR, where may initially prevent non-specific RNA binding, is partially displaced from FluPol(E). It becomes available for recruiting apo-NPs that successively will bind the replication product bulging out of the replication complex. Concomitantly, NPs are transferred from the ingoing to outgoing template in the parental replicase RNP. **(F)** FluPol(R) processively copies the template until it reaches nucleotide 17 from the 5’ end whereupon further template translocation is normally resisted by the tight binding of the 5’ hook. In the case of transcription of the vRNA template, this leads to poly-adenylation through repeated copying of U17. However, in replication, FluPol(R) manages to read-through to the end of the template. At this stage, a progeny nucleocapsid has formed, packaging the almost full-length replication product. **(G)** Replication termination. To synthetize a full-length complementary replication product, the 5’ end of the template must be released from its binding site. The mechanism for this is unknown. However, the trimeric replication complex structures observed for FluPolB suggests that a FluPol(S) from the apo-dimer could interact with FluPol(R), forcing release of the 5’ hook and stabilising the replicase while it copies the last nucleotides up to the template 5’ end. **(H)** Replication complex dissociation and recycling. Once replication is terminated, FluPol recycling can occur as previously proposed for the transcription cycle (Wandzik et al., 2020). FluPol(R), within the parental RNP, can perform another round of replication (or potentially transcription). The erstwhile FluPol(E), now part of the progeny RNP, can become a replicase or transcriptase (in the case of a progeny vRNP).

To elaborate in more detail the mechanism of replication it will be important to determine structural snapshots of replication in action, as has recently been done for transcription (Kouba et al., 2019; Wandzik et al., 2020). This would visualise the trajectory of the nascent RNA replicate from the replicase into the encapsidase and subsequent conformational changes that might occur as elongation proceeds. Ultimately, one would like to validate the NP recruitment model leading to RNP assembly.

## Acknowledgments

We thank Aldo R. Camacho-Zarco for providing hANP32A truncated constructs proteins. We thank Wojtek Galej, Sarah Schneider and Romain Linares for access to the Glacios at EMBL Grenoble; Aymeric Peuch for support using the joint EMBL-IBS computer cluster and Caroline Mas help with the Mass Photometer measurements. We acknowledge the European Synchrotron Radiation Facility and Grenoble Partnership for Structural Biology for access to the Titan Krios CM01 and Romain Linares and Gregory Effantin for assistance with data collection. We thank Matthias Budt and Thorsten Wolff (Robert Koch Institute, Berlin) for providing the HEK-293T ANP32AB KO cells and Sylvain Paisant (Institut Pasteur) for helping with plasmid mutagenesis and purification.

## Author contributions

SC and BA conceived the project. MP and BA performed cloning. PD did initial biochemical studies on FluPol-hANP32A complexes. BA performed all other *in vitro* biochemical, biophysical and cryo-EM analyses. SC did model building and refinement. Discussions with MB led to inclusion of Figure 3I. TK, supervised by NN, performed all cellular assays. SC and BA prepared the manuscript with input from all authors, especially TK and NN.

## Data availability

The data that support this study are available from the corresponding authors upon reasonable request. The coordinates and EM maps generated in this study have been deposited in the Protein Data Bank and the Electron Microscopy Data Bank (summarised in Table 1). Source data are provided in a Supplemental file associated with this paper.

## Methods

### Construction of expression plasmids for influenza A/Zhejiang/DTID-ZJU01/2013(H7N9) and B/Memphis/2003 polymerases

The previously described pFastBac Dual vector encoding for the influenza A/Zhejiang/DTID-ZJU01/2013(H7N9) polymerase subunits, PA (Uniprot: M9TI86), PB1 (Uniprot: M9TLW3), and PB2 (Uniprot: X5F427), with the mutations PA/E349K and R490I, PB1/K577G and PB2/G74R (FluPolA/H7N9-4M) (Krischuns et al., 2024) was used as a starting point. The PA N-terminal His-tag was removed by a combination of PCRs and Gibson assembly. The PB2 C-terminal Twin-strep-tag was kept. The previously described pKL vector encoding the self-cleavable poly-protein B/Memphis/2003 influenza polymerase subunits, PA (Uniprot: Q5V8Z9_9INFB), PB1 (Uniprot: Q5V8Y6_9INFB), and PB2 (Uniprot: Q5V8X3_9INFB) (Reich et al., 2014) was used as a starting point. Each FluPolB subunit was amplified and inserted in a pLIB plasmid by a combination of PCRs and Gibson assembly resulting in FluPolB subunits being under control of distinct polyhedrin promoters. The PA N-terminal His-tag was removed, whilst the PB2 C-terminal Twin-strep-tag was retained to enable purification. All plasmid sequences were confirmed by Sanger sequencing for each polymerase subunit.

Removal of all N- or C-terminal tags (except the PB2 C-terminal purification tag) was essential to allow replication complex formation. An N-terminal PA tag prevents formation of the full replicase conformation, which requires close contact of the cap-binding domain to the N-terminus of PA. An N-terminal PB1 extension impedes close packing against PA-C of ANP32 in the vicinity of residues 129-130, which buries the PB1 N-terminus. In the FluB replication complex, an N-terminal PB2 extension would prevent the observed contact between the encapsidase PB1-C/PB2-N helical bundle with the replicase cap-binding domain.

### FluPolA and FluPolB expression and purification

FluPolA/H7N9-WT was expressed and purified as previously described (Kouba et al., 2019). FluPolA/H7N9-4M and FluPolB were produced using the baculovirus expression system in *Trichoplusia ni* High 5 cells. For large-scale expression, cells at 0.8-10^6^ cells/mL concentration were infected by adding 1% of virus. Expression was stopped 72 to 96 h after the day of proliferation arrest and cells were harvested by centrifugation (1000g, 20 min at 4°C). Cells were disrupted by sonication for 5 min (5 s ON, 20 s OFF, 40% amplitude) on ice in lysis buffer (50 mM HEPES pH 8, 500 mM NaCl, 2 mM TCEP, 5% glycerol) with cOmplete EDTA-free Protease Inhibitor Cocktail (Roche). After lysate centrifugation at 48,000g for 45 min at 4°C, ammonium sulphate was added to the supernatant at 0.5 g/mL final concentration. The recombinant protein was then collected by centrifugation (45 min, 4°C at 70.000g), re-suspended in the lysis buffer, and the procedure was repeated. FluPol was purified using strep-tactin affinity purification beads (IBA, Superflow). Bound proteins were eluted using the lysis buffer supplemented by 2.5 mM d-desthiobiotin and protein-containing fractions were pooled and diluted with an equal volume of buffer (50 mM HEPES pH 8, 2 mM TCEP, 5% glycerol) before loading on an affinity column HiTrap Heparin HP 5 mL (Cytiva). A continuous gradient of lysis buffer supplemented with 1 M NaCl was applied over 15 CV, and FluPol was eluted as single species at ∼800 mM NaCl. Pure and acid nucleic free FluPol were dialysed overnight in a final buffer (50 mM HEPES pH 8, 500 mM NaCl, 2 mM TCEP, 5% glycerol), concentrated with Amicon Ultra-15 (50 kDa cutoff), flash-frozen and stored at - 80°C for later use.

### Human Acidic Nuclear Phosphoprotein 32A (hANP32A)

Human ANP32A (hANP32A) was cloned and expressed as previously described (Krischuns et al., 2024). The N-terminal His-tagged hANP32A construct was expressed in BL21(DE3) *E.coli* cells. Expression was induced with 1 mM IPTG, for 4 h at 37°C. Cells were harvested by centrifugation (1000g, 20 min at 4°C), disrupted by sonication for 5 min (5 s ON, 15 s OFF, 50% amplitude) on ice in lysis buffer (50 mM HEPES pH 8, 150 mM NaCl, 5 mM beta-mercaptoethanol (BME)) with cOmplete EDTA-free Protease Inhibitor Cocktail (Roche). After lysate centrifugation at 48,000g for 45 min at 4°C, the soluble fraction was loaded on a HisTrap HP 5 mL column (Cytiva). Bound proteins were subjected to a wash step using the lysis buffer supplemented by 50 mM imidazole. Remaining bound protein was eluted using the lysis buffer supplemented by 500 mM imidazole. Fractions containing hANP32A were dialysed overnight in the lysis buffer (50 mM HEPES pH 8, 150 mM NaCl, 5 mM BME) together with N-terminal his-tagged TEV protease (ratio 1:5 w/w). Tag-cleaved hANP32A protein was subjected to a Ni-sepharose affinity column to remove the TEV protease, further concentrated with Amicon Ultra-15 (3 kDa cutoff) and subjected to a Size-Exclusion Chromatography using a Superdex 200 Increase 10/300 GL column (Cytiva) in a final buffer containing 50 mM HEPES pH 8, 150 mM NaCl, 2 mM TCEP. Fractions containing exclusively hANP32A were concentrated with Amicon Ultra-15 (3 kDa cutoff), flash-frozen and stored at -80°C for later use.

Truncated hANP32A constructs (1-199 and 144-249) were generated, expressed and purified as previously described (Camacho-Zarco et al., 2020). The hANP32A 1-149 construct was a gift from Cynthia Wolberger (Addgene plasmid # 67241, (Huyton and Wolberger, 2007)) and was expressed and purified as previously described (Camacho-Zarco et al., 2020).

### Analytical size exclusion chromatography

Size exclusion chromatography (SEC) experiments for FluPolA/H7N9-4M and FluPolB were performed on a Superdex 200 Increase 3.2/300 (Cytiva) at 4°C, in a final buffer containing 50 mM HEPES pH 8, 150/300/500 mM NaCl, 2 mM TCEP. Depending on the experiment, 5 µM FluPol, 15 µM hANP32A (full-length, “1-149”, “1-199”, “144-Cter”), and 10 µM 5’ vRNA 1-12 (5’-pAGU AGU AAC AAG-3’) were used. Resulting mixtures were incubated 1h on ice before injection onto the column. SEC fractions of interest were loaded on 4-20% Tris-glycine gel (ThermoFisher) and stained with Coommassie Blue.

Size exclusion chromatography (SEC) experiments for FluPolA/H7N9-WT were performed on a Superdex 200 Increase 3.2/300 (Cytiva) at 4°C, in a final buffer containing 25 mM HEPES pH 8, 200/650 mM NaCl, 2 mM TCEP. Depending on the experiment, 10 µM FluPol, 5 µM hANP32A, and 10 µM 5’ vRNA 1-14 (5’-pAGU AGU AAC AAG AG)/ 3’ 1-18 (5’-UAU ACC UCU GCU UCU GCU -3’) were used. Resulting mixtures were incubated 1h on ice before injection onto the column. SEC fractions of interest were loaded on 4-20% Tris-glycine gel (ThermoFisher) and stained with Coommassie Blue.

### Mass photometry analysis

Mass photometry measurements were performed on a OneMP mass photometer (Refeyn). Coverslips (No. 1.5H, 24x50mm, VWR) were washed with water and isopropanol before being used as a support for silicone gaskets (CultureWellTM 423 Reusable Gaskets, Grace Bio-labs). Contrast/mass calibration was realized using native marker (Native Marker unstained protein 426 standard, LC0725, Life Technologies) with a medium field of view and monitored during 60 sec using the AcquireMP software (Refeyn). For each condition, 18 µl of buffer (50 mM HEPES pH 8, 150/300/500 mM NaCl, 2 mM TCEP) were used to find the focus. Using diluted SEC inputs, 2 µl of sample were added to reach a final FluPol concentration of 50 nM. Movies of 60 sec were recorded, processed, and mass estimation was determined automatically using the DiscoverMP software (Refeyn).

### Electron microscopy

#### FluPol A/Zhejiang/DTID-ZJU01/2013(H7N9) and B/Memphis/2003 replication complexes sample preparation

FluPolA/H7N9-4M and FluPolB replication complexes were trapped by mixing 1.15 µM FluPol with 5.75 µM hANP32A (molar ratio 1:5) in a final buffer containing 50 mM HEPES pH 8, 150 mM NaCl, 2 mM TCEP. Mix were incubated for 1 h at 4°C, centrifuged for 5 min at 11.000g and kept at 4°C before proceeding to grids freezing. For grid preparation, 1.5 µl of sample was applied on each sides of plasma cleaned (Fischione 1070 Plasma Cleaner: 1 min 10 s, 90% oxygen, 10% argon) grids (UltrAufoil 1.2/1.3, Au 300). Excess solution was blotted for 3 sec, blot force 0, 100% humidity, at 10°C, with a Vitrobot Mark IV (ThermoFisher) before plunge freezing in liquid ethane.

#### FluPol B/Memphis/2003 bound to 5’ cRNA sample preparation

The FluPolB encapsidase bound to 5’ cRNA structure was trapped by mixing 1.15 µM FluPolB with 5.75 µM hANP32A and 1.72 µM 5’ cRNA 1-12 (5’-AGC AGA AGC AGA -3’) (molar ratio 1:5:1.5) in a final buffer containing 50 mM HEPES pH 8, 150 mM NaCl, 2 mM TCEP. The mix was incubated for 1 h at 4°C, centrifuged for 5 min at 11000g and kept at 4°C before proceeding to grid freezing. For grid preparation, 1.5 µl of sample was applied on each sides of plasma cleaned (Fischione 1070 Plasma Cleaner: 1 min 10 s, 90% oxygen, 10% argon) grids (UltrAufoil 1.2/1.3, Au 300). Excess solution was blotted for 3 sec, blot force 0, 100% humidity, at 10°C, with a Vitrobot Mark IV (ThermoFisher) before plunge freezing in liquid ethane.

### Cryo-EM data collection

#### FluPol A/Zhejiang/DTID-ZJU01/2013(H7N9) and B/Memphis/2003 replication complexes

Automated data collections were performed on a TEM Titan Krios G3 (ThermoFisher) operated at 300 kV equipped with a K3 direct electron detector camera (Gatan) and a BioQuantum energy filter (Gatan), using EPU (ThermoFisher). Coma and astigmatism correction were performed on a carbon grid. Micrographs were recorded in counting mode at a ×105,000 magnification giving a pixel size of 0.84 Å with defocus ranging from −0.8 to −2.0 µm. Gain-normalised movies of 40 frames were collected with a total exposure of ∼40 e^−^/Å^2^.

#### FluPol B/Memphis/2003 bound to 5’ cRNA sample preparation

Automated data collection was performed on a TEM Glacios (ThermoFisher) operated at 200 kV equipped with a F4i direct electron detector camera (ThermoFisher) and a SelectrisX energy filter (ThermoFisher), using EPU (ThermoFisher). Coma and astigmatism correction were performed on a carbon grid. Micrographs were recorded in counting mode at a ×130,000 magnification giving a pixel size of 0.878 Å with defocus ranging from −0.8 to −2.0 µm. EER movies were collected with a total exposure of ∼40 e^−^/Å^2^.

### Image processing

#### FluPol A/Zhejiang/DTID-ZJU01/2013(H7N9) structure determination

For the FluPolA TEM Titan Krios dataset, 14,001 movies were collected. Movie drift correction was performed using Relion’s Motioncor implementation, with 7x5 patch, using all movie frames (Zheng et al., 2017). All additional initial image processing steps were performed in CryoSPARC v4.3 (Punjani et al., 2017). CTF parameters were determined using “Patch CTF estimation”. Realigned micrographs were then manually inspected and low-quality images were manually discarded resulting in 13,328 micrographs kept. Particles were automatically picked using a circular blob with a diameter ranging from 110 to 130 Å, and extracted using a box size of 420 x 420 pixels^2^, Fourier cropped to 210 x 210 pixels^2^. Successive 2D classifications were used to eliminate particles displaying poor structural features, and coarsely separate monomers from dimers. Monomers were subjected to a “heterogeneous refinement” job. Particles displaying PA-ENDO in the replicase conformation (PA-ENDO(R)), the rest of them displaying a dislocated FluPol core, were Fourier uncropped and subjected to a “non-uniform refinement” job. Based on the estimated particle angles and shifts, a “3D classification” job was performed. For each relevant FluPol conformation, particles were grouped and subjected to a final “non-uniform refinement”. FluPolA/H7N9-4M asymmetric dimers were first subjected to a “heterogeneous refinement” job. Particles assigned to the 3D class displaying well-defined secondary structures were used for model training and picking using Topaz (Bepler et al., 2019). The resulting picked particles were extracted and subjected to 2D classification. All asymmetric dimers particles were merged, the duplicates removed, Fourier uncropped, and then subjected to a “non-uniform refinement” job. To alleviate the preferential orientation problem of the FluPolA/H7N9-4M replication complex, a “3D classification” job was used. Particles displaying a proper view distribution equilibrium were used and subjected to a “non-uniform refinement”. Based on this consensus map, particle subtraction around “FluPol(R) minus 627(R)” and “FluPol(E)-hANP32A-627(R)” was performed. The subtracted particles were finally subjected to local refinement to improve subtracted particle angles and shifts estimation. Post-processing was performed in CryoSPARC using an automatically or manually determined B-factor. For each final map, reported global resolution is based on the FSC 0.143 cut-off criteria. Local resolution variations were estimated in CryoSPARC. The detailed image processing pipeline is shown in Supp. Info. 1-2.

#### FluPol B/Memphis/2003 structure determination

For the FluPolB TEM Titan Krios dataset, 15,650 movies were collected. Movie drift correction was performed using Relion’s Motioncor implementation, with 7x5 patch, using all movie frames (Zheng et al., 2017). All additional initial image processing steps were performed in cryoSPARC v4.3 (Punjani et al., 2017). CTF parameters were determined using “Patch CTF estimation”, realigned micrographs were then manually inspected and low-quality images were manually discarded resulting in 15,234 micrographs kept. Particles were automatically picked using a circular blob with a diameter ranging from 110 to 140 Å and extracted using a box size of 480 x 480 pixels^2^, Fourier cropped to 200 x 200 pixels^2^. Successive 2D classifications using a circular mask of 210 Å were used to eliminate particles displaying poor structural features. Following initial 2D classifications, all particles were re-extracted at a larger box size (512 x 512 pixels^2^, Fourier cropped to 200 x 200 pixels^2^) and subjected to multiple 2D classifications using a circular mask of 280 Å to coarsely separate dimers, monomers, and trimers. For the FluPolB symmetrical dimers, following an “ab-initio” reconstruction job, particles displaying one FluPolB(E) were Fourier uncropped and subjected to a “non-uniform refinement” job, followed by respective FluPolB symmetrical (FluPolB(S)) and FluPolB(E) signal subtraction. After subsequent local refinements, “3D classification” jobs were performed to separate the different FluPolB states. 3D classes displaying a complete FluPol(E) conformation were grouped, locally refined, and subjected to a final “non-uniform refinement” using the un-subtracted particles. A similar approach was used for the different FluPol(S) conformations (core, ENDO(R), ENDO(E) or ENDO(T)) (Supp. Info. 3). Dimers displaying two FluPolB core were Fourier uncropped and subjected to a “non-uniform refinement” job followed by a “3D classification”. Particles displaying one FluPolB with PA-ENDO in a transcriptase conformation (ENDO(T)) were grouped and subjected to a final “non-uniform refinement” job (Supp. Info. 4). For the FluPolB monomers, particles were subjected to an “ab-initio” reconstruction followed by a “non-uniform refinement”. Subsequent “3D classification” allowed isolation of monomeric apo-FluPolB(E). Particles were Fourier uncropped and subjected to a final “non-uniform refinement” job (Supp. Info. 4). For the FluPolB trimers (FluPolB replication complex plus one FluPol(S)), particles were subjected to an “ab-initio” reconstruction job. The few particles displaying a well-defined FluPolB replication complex were Fourier uncropped and subjected to a “non-uniform refinement” job. Particle subtraction was performed on “FluPolB(S) + FluPolB(E)”, “FluPolB(S)+FluPolB(R)” and “FluPolB(S)” moieties, followed by local refinements to improve subtracted particle angles and shifts estimation. (Supp. Info. 5). Post-processing was performed in CryoSPARC using an automatically or manually determined B-factor. For each final map, reported global resolution is based on the FSC 0.143 cut-off criteria. Local resolution variations were estimated in CryoSPARC. The detailed image processing pipeline is shown in Supp. Info. 3-5.

#### 5’ cRNA bound FluPol B/Memphis/2003 structure determination

For the TEM Glacios dataset, 2,451 movies were collected. Movie drift correction was performed using Relion’s Motioncor implementation, with 7x5 patch, using all movie frames (Zheng et al., 2017). All additional initial image processing steps were performed in CryoSPARC v4.3 (Punjani et al., 2017). CTF parameters were determined using “Patch CTF estimation”, realigned micrographs were then manually inspected and low-quality images were manually discarded resulting in 2,353 micrographs kept. Particles were automatically picked using a circular blob with a diameter ranging from 110 to 140 Å and extracted using a box size of 380 x 380 pixels^2^, Fourier cropped to 240 x 240 pixels^2^. Successive 2D classifications using a circular mask of 210 Å were used to eliminate particles displaying poor structural features. Remaining particles were subjected to a “heterogeneous refinement” job. Particles belonging to the class corresponding to 5’ cRNA bound FluPolB(E) were subjected to a “non-uniform refinement” job, followed by “3D classification”. A final “non-uniform refinement” has been done with particles displaying a complete FluPol(E) conformation. Post-processing was performed in CryoSPARC using an automatically or manually determined B-factor. For each final map, reported global resolution is based on the FSC 0.143 cut-off criteria. Local resolution variations were estimated in CryoSPARC. The detailed image processing pipeline is shown in Supp. Info. 7.

### Model building and refinement

Atomic models were constructed by iterative rounds of manual model building with COOT (Emsley and Cowtan, 2004) and real-space refinement using Phenix, with Ramachandran restraints (Afonine et al., 2018). For model building of the replicase-moiety of the FluPolB replication complex, the previously determined replicase-like structure (PDB: 5EPI)(Thierry et al., 2016) was used as starting point. The FluPolB encapsidase conformation was initially constructed from the higher resolution symmetrical dimer map and transferred to the replicase complex. For FluPolA replication complex structure building, a variety of previous A/H7N9 structures were used as starting models.

Validation was performed using Phenix. Model resolution according to the cryo-EM map was estimated at the 0.5 FSC cutoff. Structural analysis was performed in Coot and Chimera (Pettersen et al., 2004). Electrostatic potential surfaces were calculated using the APBS-PDB2PQR software suite (Jurrus et al., 2018). Buried solvent accessible surfaces were calculated using PISA (Krissinel and Henrick, 2007) at the PDBe. Figures were generated using ChimeraX (Goddard et al., 2018).

### Cells

HEK-293T cells (ATCC CRL-3216) and HEK-293T (ATCC CRL-11268) ANP32AB KO cells (Krischuns et al., 2024) were grown in complete Dulbecco’s modified Eagle’s medium (DMEM, Gibco) supplemented with 10% fetal bovine serum (FBS) and 1% penicillin-streptomycin (Gibco). Cell cultures were PCR-tested regularly to ensure absence of mycoplasma contamination. Cells (3E04/well) were seeded in 96-well white plates (Greiner Bio-One) the day before transfection with polyethyleneimine (PEI-max, #24765-1 Polysciences Inc).

### Plasmids used in cell-based assays

The pcDNA3.1-hANP32A-FLAG, A/WSN/33 (WSN) pcDNA3.1-PB2, -PB1, -PA, pCI-NP and B/Memphis/13/2003 (Memphis) pcDNA3.1-PB2, -PB1, -PA, -NP plasmids were described previously (Lukarska et al. PMID: PMID: 28002402, Reich et al. (Camacho-Zarco et al., 2023; Lukarska et al., 2017; Reich et al., 2014). Plasmids used for vRNP reconstitution assays and the WSN pCI-PB1-luc1, Memphis pCI-PB1-luc1, pCI-hANP32A-luc2, pCI-chANP32A-luc2 plasmids used for split-luciferase-based complementation assays were described previously (Krischuns et al., 2024; Krischuns et al., 2022). The pCI-hANP32B-luc2 plasmid was constructed by replacing the hANP32A sequence in the pCI-hANP32A-luc2 plasmid. pcDNA3.1-hANP32B-FLAG, -chANP32A-FLAG were constructed by replacing the hANP32A sequence in the pcDNA3.1-hANP32A-FLAG plasmid. All mutations were introduced by an adapted QuikChange site-directed mutagenesis (Agilent Technologies) protocol (Zheng et al., 2004). ORFs were verified by Sanger sequencing and primer and plasmid sequences are available with annotations as a supplementary data file.

### vRNP reconstitution assays

HEK-293T cells were co-transfected with plasmids encoding the vRNP protein components (PB2, PB1, PA, NP), a pPolI-Firefly plasmid encoding a negative-sense viral-like RNA expressing the Firefly luciferase and the pTK-Renilla plasmid (Promega) as an internal control. For FluPol activity rescue experiments in ANP32AB KO cells, a plasmid encoding either the wild-type or mutant hANP32A, hANP32B or chANP32A protein was co-transfected. Mean relative light units (RLUs) produced by the Firefly and Renilla luciferase, reflecting the viral polymerase activity and transfection efficiency, respectively, were measured using the Dual-Glo Luciferase Assay System (Promega) on a Centro XS LB960 microplate luminometer (Berthold Technologies, MikroWin Version 4.41) at 48 hours post-transfection (hpt). Firefly luciferase signals were normalised with respect to Renilla luciferase signals. At least three independent experiments (each in technical duplicates) were performed, and each biological replicate is represented as a dot in the graphs. Plasmid combinations, orientations of tags as well as plasmid amounts used for transfections in a given experiment are available as a source data file.

### Protein complementation assays

HEK-293T cells were co-transfected with plasmids encoding the FluPol subunits (PB2, PB1-G1, PA) and an ANP32A protein (hANP23A-G2, hANP32B-G2 or chANP32A-G2). Cells were lysed 20-24 hpt in Renilla lysis buffer (Promega) for 45 min at room temperature under steady shaking. RLUs produced by the reconstituted Gaussia princeps luciferase, reflecting the FluPol-ANP32 interaction, were measured on a Centro XS LB960 microplate luminometer (Berthold Technologies, MikroWin Version 4.41) using a reading time of 10 s upon injection of 50 µl Renilla luciferase reagent (Promega). Three independent experiments (each in technical triplicates) were performed, and each biological replicate is represented as a dot in the graphs. Plasmid combinations, orientations of tags as well as plasmid amounts used for transfections in a given experiment are available as a source data file.

### Antibodies and immunoblots

Total cell lysates were prepared in RIPA cell lysis buffer as described (Krischuns et al., 2018). Proteins were separated by SDS-PAGE using NuPAGE™ 4-12% Bis-Tris gels (Invitrogen) and transferred to nitrocellulose membranes which were incubated with primary antibodies directed against PA ((Da Costa et al., 2015), 1:2500), PB2 (GTX125925, GeneTex, 1:5,000), Gaussia princeps luciferase (New England Biolabs, #E8023, 1:5,000), Histone H3 (Cell Signaling Technology, #9715, 1:1,000), Tubulin (B-5-1-2, Sigma Aldrich, 1:10,000) and subsequently with HRP-tagged secondary antibodies (Jackson Immunoresearch, 1:10,000). Membranes were revealed with the ECL2 substrate according to the manufacturer’s instructions (Pierce). Chemiluminescence signals were acquired using the ChemiDoc imaging system (Bio-Rad, Image Lab Touch Software 2.4.0.03) and analysed with ImageLab (Bio-Rad, Image Lab 6.0.1 build 34). Uncropped gels are provided as a source data file.

## Supplemental information

**Tables S1-S4. Cryo-EM data collection, refinement and validation statistics**

**Figures S1-S8**

**Supplemental Information 1-5 and 7. Cryo-EM data processing pipelines**

**Supplemental Information 6. ANP32 multiple sequence alignment.**

**Table S1.** Cryo-EM data collection, refinement and validation statistics for A/H7N9-4M polymerase with hANP32A.

| Sample | A/H7N9-4M polymerase with hANP32 |  |  |  |  |
| --- | --- | --- | --- | --- | --- |
| Structure | Endo(R) core1 | Endo(R) core2 | Focus replicase minus 627(R) | Focus Encapsidase plus 627(R) hANP32 | Complete replication complex |
| PDB ID | PDB ID 8RMP | PDB ID 8RMQ | PDB ID 8RMS | PDB ID 8RN0 | PDB ID 8RMR |
| EMDB ID | EMD-19366 | EMD-19367 | EMD-19369 | EMD-19382 | EMD-19368 |
| Data collection and processing |  |  |  |  |  |
| Microscope | ThermoFisher Krios TEM |  |  |  |  |
| Voltage (kV) | 300 |  |  |  |  |
| Camera | Gatan K3 direct electron detector mounted on a Gatan Bioquantum LS/967 energy filter |  |  |  |  |
| Magnification | 105,000 |  |  |  |  |
| Nominal defocus range (μm) | -0.8 / -2 |  |  |  |  |
| Electron exposure (e−/Å²) | 40 |  |  |  |  |
| Pixel size (Å) | 0.84 |  |  |  |  |
| Initial micrographs (no.) | 14,001 |  |  |  |  |
| Final micrographs (no.) | 13,328 |  |  |  |  |
| Refinement |  |  |  |  |  |
| Particles per class (no.) | 56,572 | 277,339 | 18,596 |  |  |
| Map resolution (Å), 0.143 FSC | 2.77 | 2.54 | 3.21 | 3.13 | 3.25 |
| Model resolution (Å), 0.5 FSC | 3.0 | 2.6 | 3.76 | 3.60 | 3.80 |
| Map sharpening B factor (Å²) | -75 | -87 | -56 | -56 | -55 |
| Map versus model cross-correlation (CCmask) | 0.8241 | 0.8462 | 0.7118 | 0.8241 | 0.8462 |
| Model composition |  |  |  |  |  |
| Non-hydrogen atoms | 13818 | 13676 | 16190 | 18082 | 34215 |
| Protein residues | 1717 | 1699 | 2019 | 2262 | 4273 |
| Water | - | 7 | - | - | - |
| Ligands | 1 Mg | 2 Mg | 1 Mg | - | 2 Mg |
| B factors (Å²) |  |  |  |  |  |
| Protein | 79.96 | 49.91 | 99.99 | 116.09 | 100.81 |
| Water | - | 21.56 | - | - | - |
| RMS deviations |  |  |  |  |  |
| Bond lengths (Å) | 0.003 | 0.003 | 0.002 | 0.002 | 0.002 |
| Bond angles (°) | 0.512 | 0.484 | 0.466 | 0.482 | 0.502 |

| Validation |  |  |  |  |  |
| --- | --- | --- | --- | --- | --- |
| MolProbity score | 1.76 | 1.65 | 1.58 | 1.76 | 1.77 |
| All-atom clash score | 4.49 | 4.57 | 7.06 | 8.05 | 8.85 |
| Poor rotamers (%) | 2.95 | 2.45 | 0.17 | 0.15 | 0.26 |
| Ramachandran plot |  |  |  |  |  |
| Favored (%) | 97.00 | 97.38 | 96.85 | 95.44 | 95.73 |
| Allowed (%) | 2.94 | 2.62 | 3.15 | 4.43 | 4.20 |
| Outliers (%) | 0.06 | 0.0 | 0.0 | 0.13 | 0.07 |

**Table S2.**
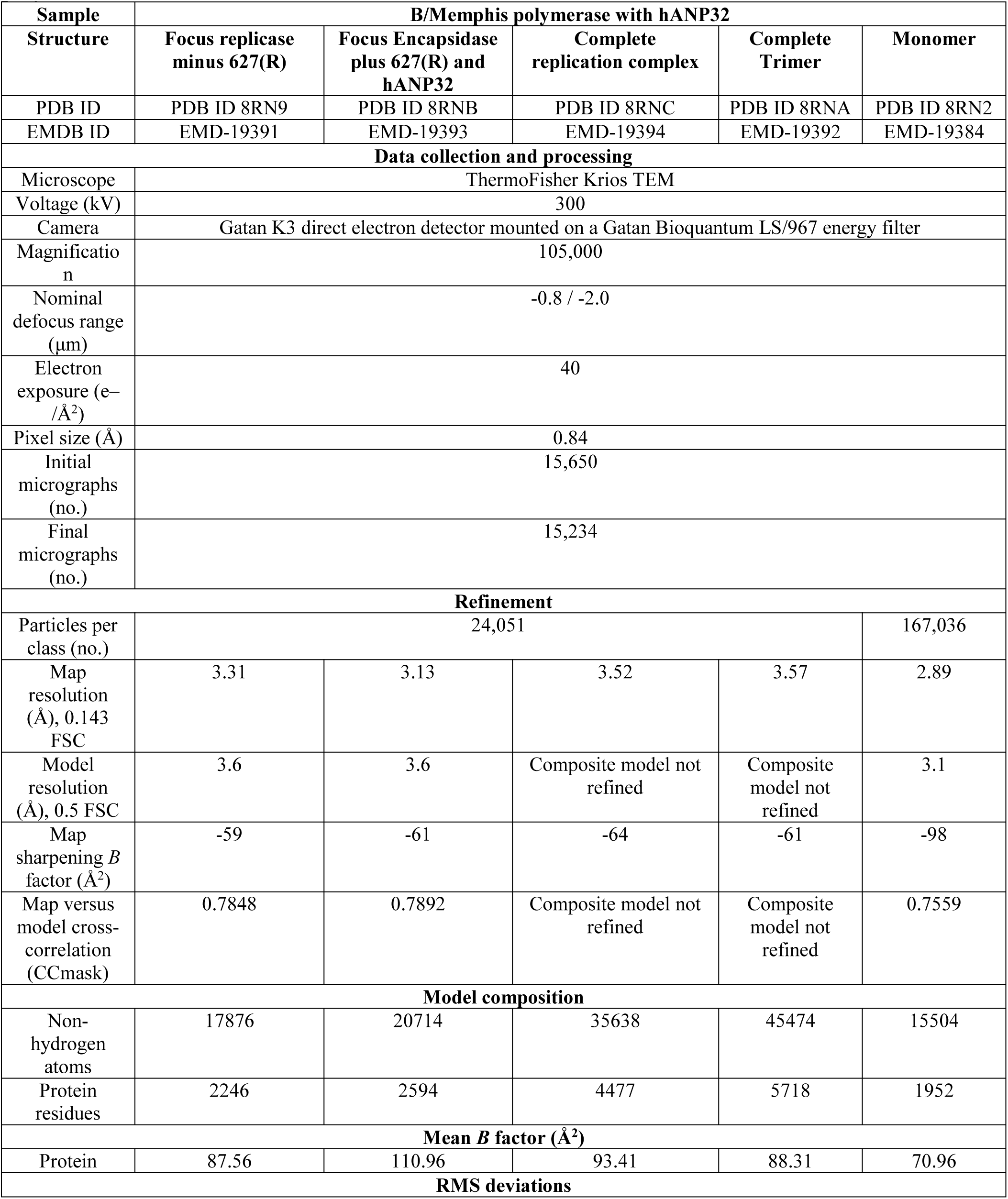
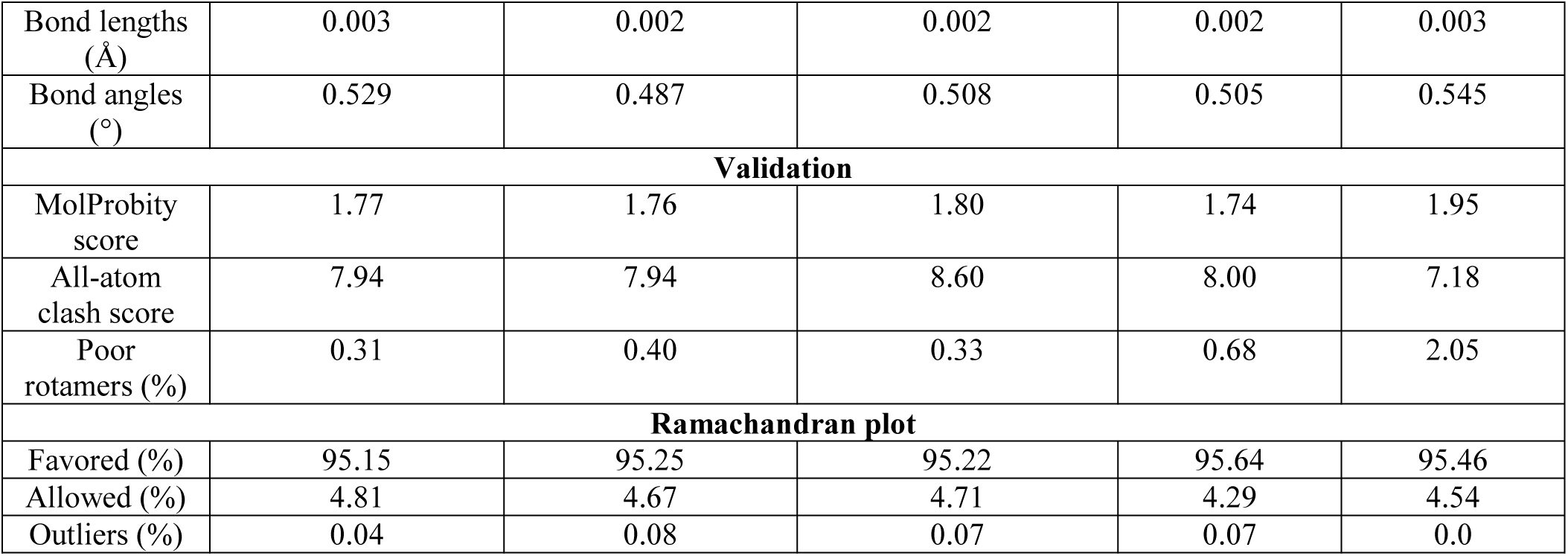
Cryo-EM data collection, refinement and validation statistics for B/Memphis polymerase with hANP32A.

**Table S3.**
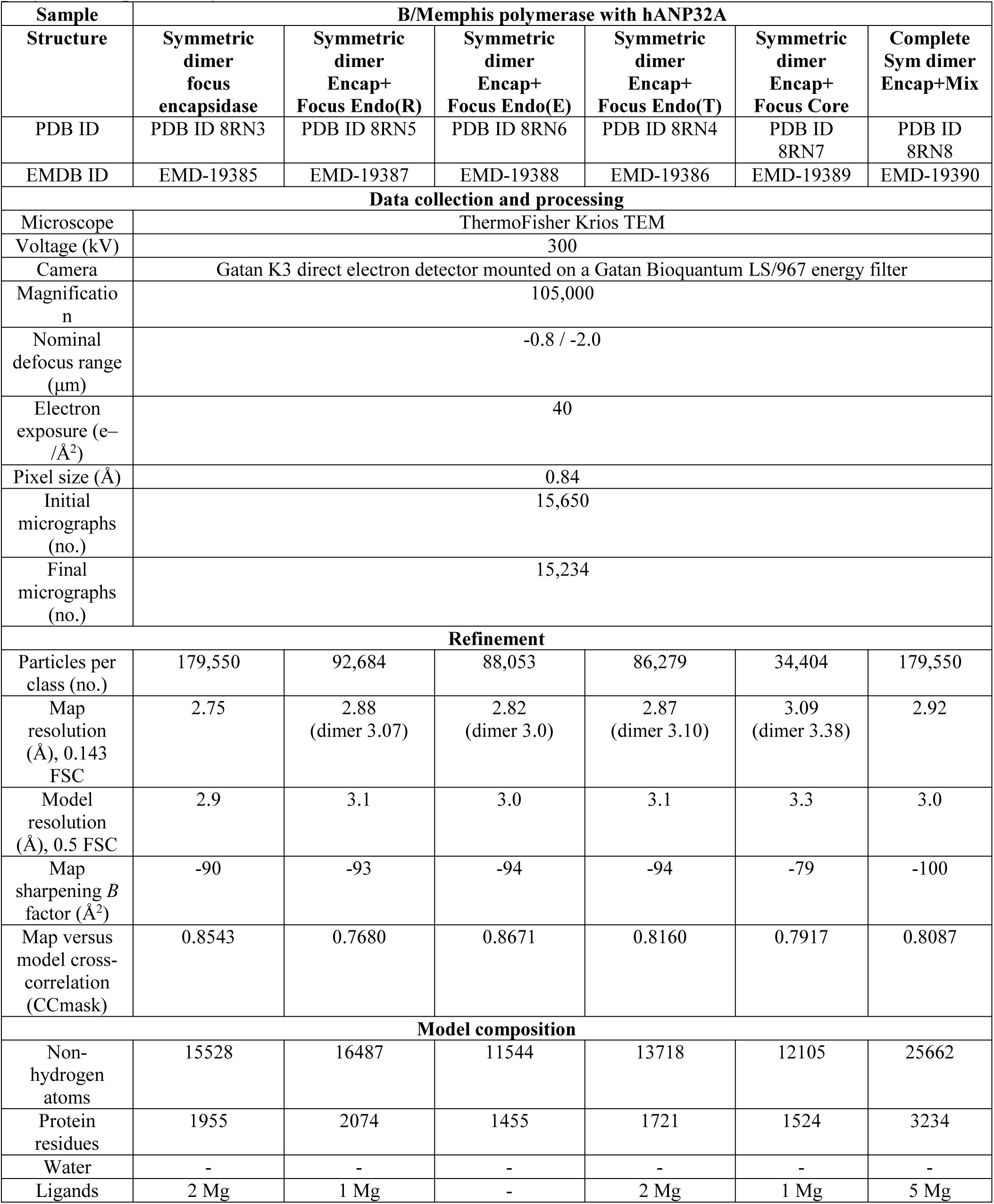
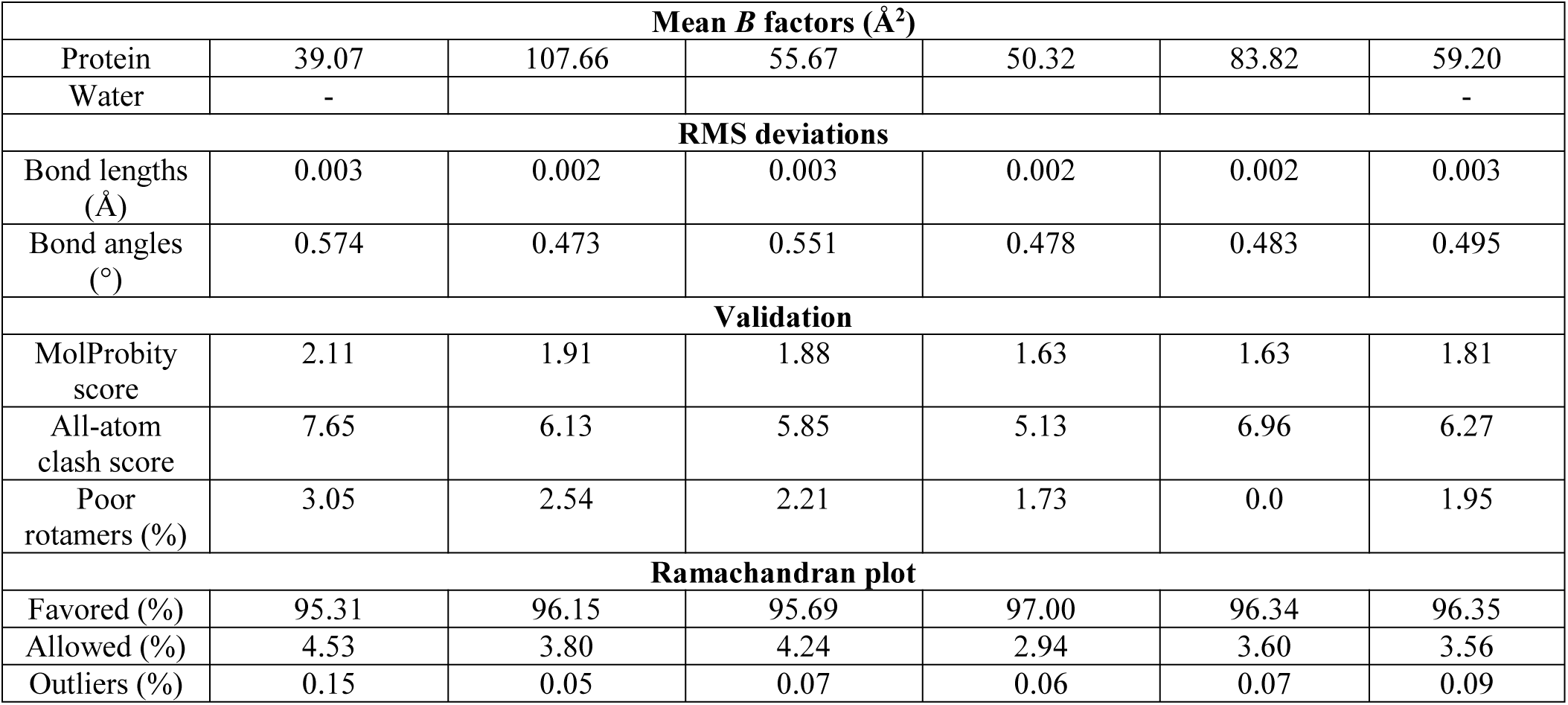
Cryo-EM data collection, refinement and validation statistics for B/Memphis polymerase pseudo-symmetric dimer.

| Sample | B/Memphis polymerase with hANP32A |  |  |  |  |  |
| --- | --- | --- | --- | --- | --- | --- |
| Structure | Symmetric dimer focus encapsidase | Symmetric dimer Encap+ Focus Endo(R) | Symmetric dimer Encap+ Focus Endo(E) | Symmetric dimer Encap+ Focus Endo(T) | Symmetric dimer Encap+ Focus Core | Complete Sym dimer Encap+Mix |
| PDB ID | PDB ID 8RN3 | PDB ID 8RN5 | PDB ID 8RN6 | PDB ID 8RN4 | PDB ID 8RN7 | PDB ID 8RN8 |
| EMDB ID | EMD-19385 | EMD-19387 | EMD-19388 | EMD-19386 | EMD-19389 | EMD-19390 |
| Data collection and processing |  |  |  |  |  |  |
| Microscope | ThermoFisher Krios TEM |  |  |  |  |  |
| Voltage (kV) | 300 |  |  |  |  |  |
| Camera | Gatan K3 direct electron detector mounted on a Gatan Bioquantum LS/967 energy filter |  |  |  |  |  |
| Magnification | 105,000 |  |  |  |  |  |
| Nominal defocus range (μm) | -0.8 / -2.0 |  |  |  |  |  |
| Electron exposure (e−/Å²) | 40 |  |  |  |  |  |
| Pixel size (Å) | 0.84 |  |  |  |  |  |
| Initial micrographs (no.) | 15,650 |  |  |  |  |  |
| Final micrographs (no.) | 15,234 |  |  |  |  |  |
| Refinement |  |  |  |  |  |  |
| Particles per class (no.) | 179,550 | 92,684 | 88,053 | 86,279 | 34,404 | 179,550 |
| Map resolution (Å), 0.143 FSC | 2.75 | 2.88 (dimer 3.07) | 2.82 (dimer 3.0) | 2.87 (dimer 3.10) | 3.09 (dimer 3.38) | 2.92 |
| Model resolution (Å), 0.5 FSC | 2.9 | 3.1 | 3.0 | 3.1 | 3.3 | 3.0 |
| Map sharpening B factor (Å²) | -90 | -93 | -94 | -94 | -79 | -100 |
| Map versus model cross-correlation (CCmask) | 0.8543 | 0.7680 | 0.8671 | 0.8160 | 0.7917 | 0.8087 |
| Model composition |  |  |  |  |  |  |
| Non-hydrogen atoms | 15528 | 16487 | 11544 | 13718 | 12105 | 25662 |
| Protein residues | 1955 | 2074 | 1455 | 1721 | 1524 | 3234 |
| Water | - | - | - | - | - | - |
| Ligands | 2 Mg | 1 Mg | - | 2 Mg | 1 Mg | 5 Mg |

| Mean *B* factors (Å<sup>2</sup>) |  |  |  |  |  |  |
| --- | --- | --- | --- | --- | --- | --- |
| Protein | 39.07 | 107.66 | 55.67 | 50.32 | 83.82 | 59.20 |
| Water | - |  |  |  |  | - |
| RMS deviations |  |  |  |  |  |  |
| Bond lengths (Å) | 0.003 | 0.002 | 0.003 | 0.002 | 0.002 | 0.003 |
| Bond angles (°) | 0.574 | 0.473 | 0.551 | 0.478 | 0.483 | 0.495 |
| Validation |  |  |  |  |  |  |
| MolProbity score | 2.11 | 1.91 | 1.88 | 1.63 | 1.63 | 1.81 |
| All-atom clash score | 7.65 | 6.13 | 5.85 | 5.13 | 6.96 | 6.27 |
| Poor rotamers (%) | 3.05 | 2.54 | 2.21 | 1.73 | 0.0 | 1.95 |
| Ramachandran plot |  |  |  |  |  |  |
| Favored (%) | 95.31 | 96.15 | 95.69 | 97.00 | 96.34 | 96.35 |
| Allowed (%) | 4.53 | 3.80 | 4.24 | 2.94 | 3.60 | 3.56 |
| Outliers (%) | 0.15 | 0.05 | 0.07 | 0.06 | 0.07 | 0.09 |

**Table S4.**
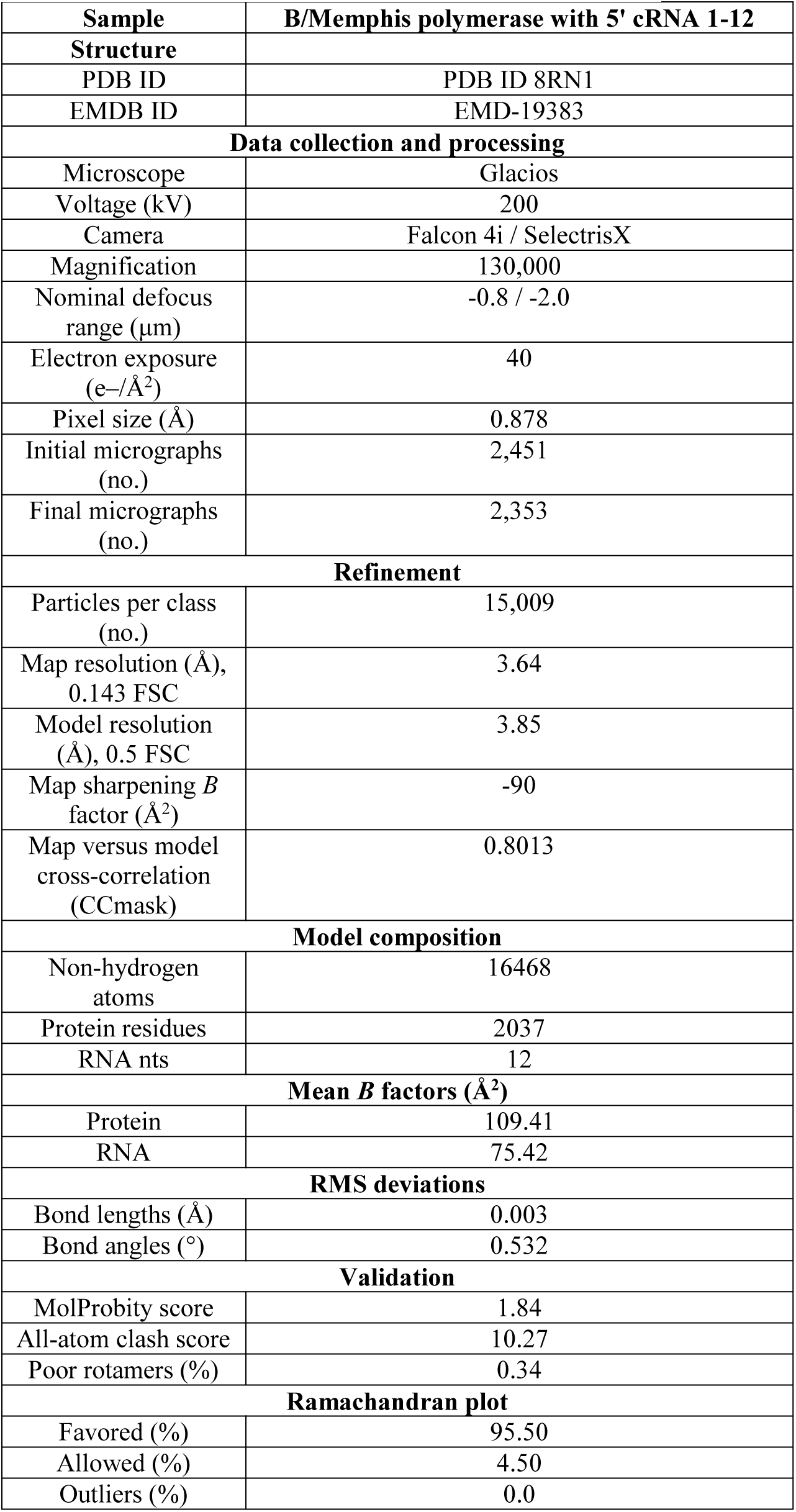
Cryo-EM data collection, refinement and validation statistics for B/Memphis encapsidase bound to 5’ cRNA 1-12.

**Figure S1.**
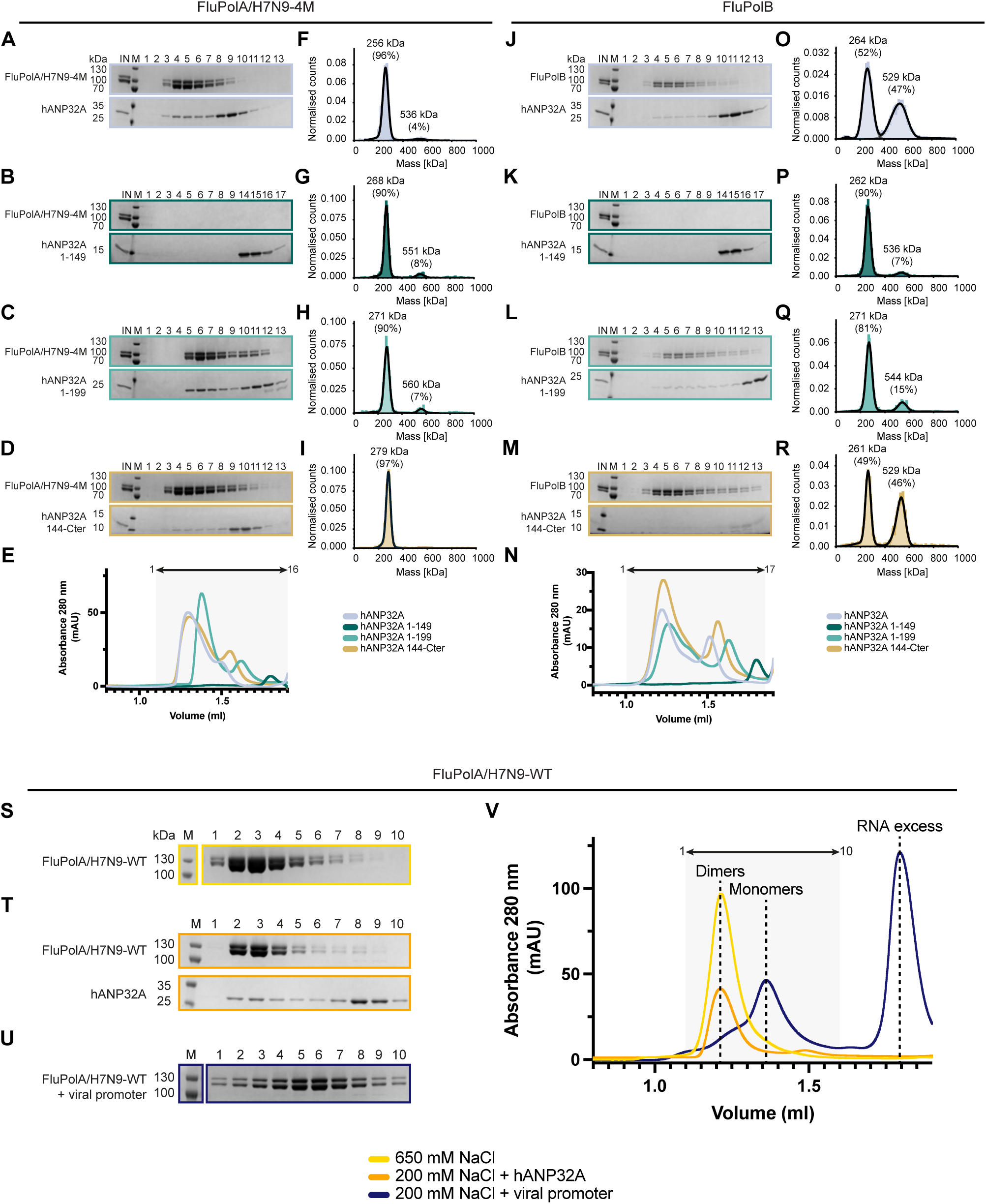
Biochemical analysis of the interaction of FluPolA/H7N9-4M and FluPolB with different hANP32A constructs. **(A)** SDS-PAGE analysis of FluPolA/H7N9-4M-hANP32A at 150 mM NaCl. The molecular ladder (M) in kDa, FluPolA/H7N9-4M heterotrimer, and hANP32A are indicated on the left of the gel. “IN” corresponds to the input. This data is also presented in Figure 1D. **(B-D)** SDS-PAGE analysis of FluPolA/H7N9-4M interaction with hANP32A 1-149 (LRR domain alone) **(B)**, hANP32A 1-199 (LRR domain with half the LCAR) **(C)**, and hANP32A 144-C-terminus (LCAR alone) **(D)** at 150 mM NaCl. The molecular ladder (M) in kDa, FluPolA/H7N9-4M heterotrimer, and hANP32A constructs are indicated on the left of the gel. “IN” corresponds to the input. **(E)** Superposition of size exclusion chromatography profiles of FluPolA/H7N9-4M with hANP32A (blue-grey), hANP32A 1-149 (dark blue), hANP32A 1-199 (green) and hANP32A 144-C-terminus (yellow) at 150 mM NaCl. SEC profile of FluPolA/H7N9-4M with hANP32A is presented in Figure 1E. The relative absorbance at 280 nm (mAU) is on the y-axis. The elution volume (ml) is on the x-axis, graduated every 50 µl. SDS-PAGE fractions 1 to 13 corresponds to the elution volume 1.1 ml - 1.75 ml, represented as an arrow on top. **(F)** Mass photometry analysis of FluPolA/H7N9-4M-hANP32A interaction at 150 mM NaCl. The determined masses in kDa of the main species are indicated. This data is also presented in Figure 1I. **(G-I)** Mass photometry analysis of FluPolA/H7N9-4M interaction with hANP32A 1-149 (LRR domain alone) **(G)**, hANP32A 1-199 (LRR domain with half the LCAR) **(H)**, and hANP32A 144-C-terminus (LCAR alone) **(I)**, at 150 mM NaCl. The determined masses in kDa of the main species are indicated. **(J)** SDS-PAGE analysis of FluPolB-hANP32A interaction at 150 mM NaCl. The molecular ladder (M) in kDa, FluPolB heterotrimer and hANP32A are indicated on the left of the gel. “IN” corresponds to the input. This data is presented in Figure 1M. **(K-M)** SDS-PAGE analysis of FluPolB interaction with hANP32A 1-149 (LRR domain alone) **(K)**, hANP32A 1-199 (LRR domain with half the LCAR) **(L)**, and hANP32A 144-C-terminus (LCAR alone) **(M)**, at 150 mM NaCl. The molecular ladder (M) in kDa, FluPolB heterotrimer, and hANP32A constructs are indicated on the left of the gel. “IN” corresponds to the input. (N) Superposition of size exclusion chromatography profiles of FluPolB with hANP32A (blue-grey), hANP32A 1-149 (dark blue), hANP32A 1-199 (green) and hANP32A 144-C-terminus (yellow) at 150 mM NaCl. SEC profile of FluPolB with hANP32A is presented in Figure 1N. The relative absorbance at 280 nm (mAU) is on the y-axis. The elution volume (ml) is on the x-axis, graduated every 50 µl. SDS-PAGE fractions 1 to 13 corresponds to the elution volume 1.0 ml - 1.65 ml, represented as an arrow on top. (O) Mass photometry analysis of FluPolB-hANP32A interaction at 150 mM NaCl. The determined masses in kDa of the main species are indicated. This data is presented in Figure 1R. **(P-R)** Mass photometry analysis of FluPolB interaction with hANP32A 1-149 (LRR domain alone) **(P)**, hANP32A 1-199 (LRR domain with half the LCAR) **(Q)**, and hANP32A 144-C-terminus (LCAR alone) **(R)**, at 150 mM NaCl. The determined masses in kDa of the main species are indicated. **(S)** SDS-PAGE analysis of FluPolA/H7N9-WT at 650 mM NaCl. The molecular ladder (M) in kDa, FluPolA/H7N9-WT heterotrimer, are indicated on the left of the gel. **(T)** SDS-PAGE analysis of FluPolA/H7N9-WT interaction with hANP32A at 200 mM NaCl. The molecular ladder (M) in kDa, FluPolA/H7N9-WT heterotrimer, and hANP32A are indicated on the left of the gel. **(U)** SDS-PAGE analysis of FluPolA/H7N9-WT in complex with vRNA promoter bound. The molecular ladder (M) in kDa, FluPolA/H7N9-WT heterotrimer are indicated on the left of the gel. **(V)** Superposition of size exclusion chromatography profiles of FluPolA/H7N9-WT at 650 mM NaCl (yellow), 200 mM NaCl with hANP32A (orange), 200 mM NaCl with vRNA promoter bound (dark blue). The relative absorbance at 280 nm (mAU) is on the y-axis. The elution volume (ml) is on the x-axis, graduated every 50 µl. SDS-PAGE fractions 1 to 10 corresponds to the elution volume 1.1 ml - 1.6 ml, represented as an arrow on top.

**Figure S2.**
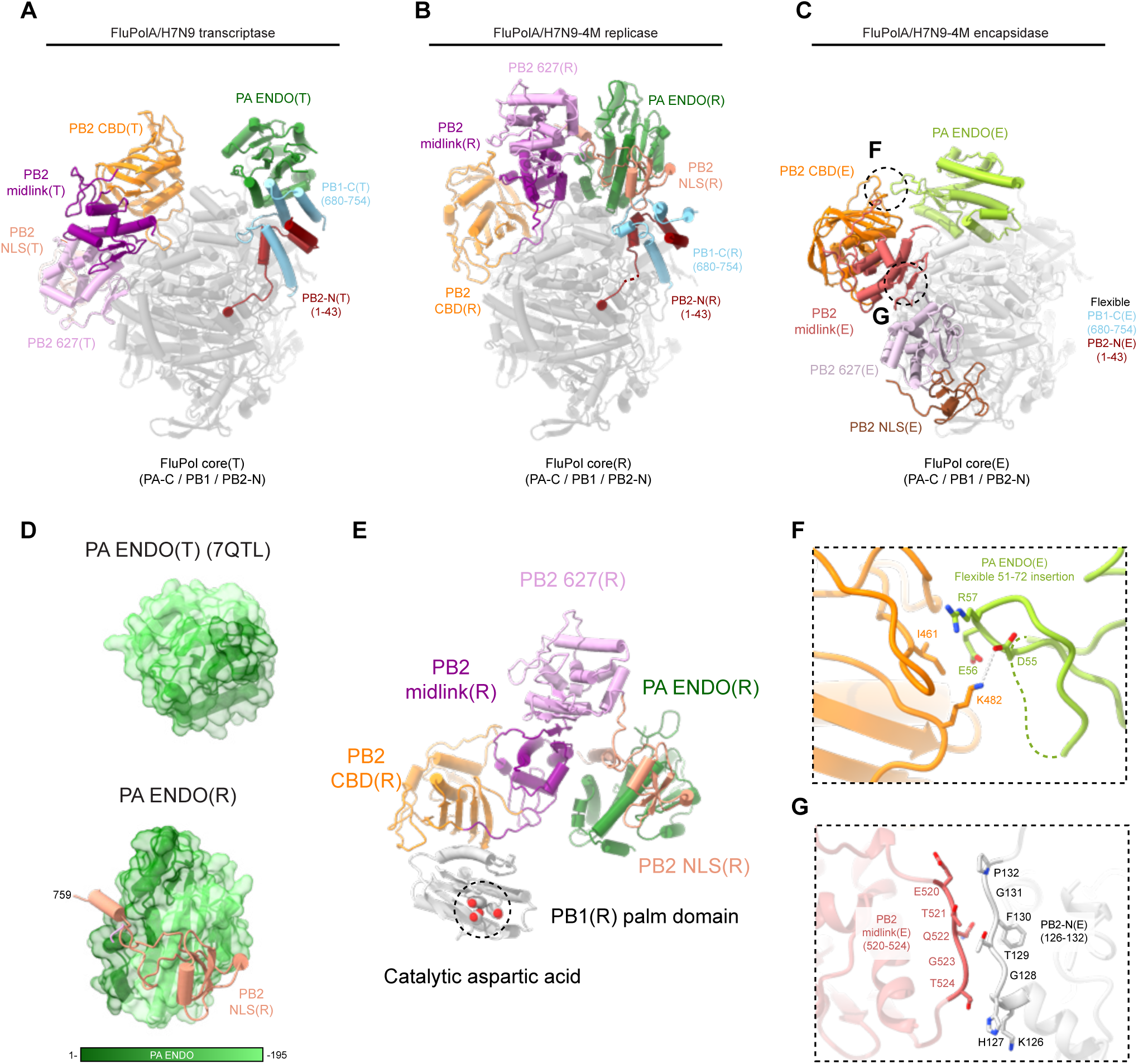
Structural comparison of FluPolA/H7N9 transcriptase, replicase and encapsidase conformations. **(A)** Cartoon representation of FluPolA/H7N9 in the transcriptase conformation (FluPolA/H7N9(T)) (PDB 7QTL). FluPolA/H7N9(T) core is dark grey, shown in transparency, PA ENDO(T) in dark green, PB1-C(T) in blue, PB2-N(T) in red, PB2 midlink(T) in magenta, PB2 CBD(T) in orange, PB2 627(T) in pink, PB2 NLS(T) in beige. **(B)** Cartoon representation of FluPolA/H7N9-4M in the replicase conformation (FluPolA/H7N9-4M(R)), extracted from the replication complex and aligned on FluPolA/H7N9(T) PB1 subunit. FluPolA/H7N9-4M(R) core is in dark grey, shown in transparency, PA ENDO(R) in dark green, PB1-C(R) in blue, PB2-N(R) in red, PB2 midlink(R) in magenta, PB2 CBD(R) in orange, PB2 627(R) in pink, PB2 NLS(R) in beige. **(C)** Cartoon representation of FluPolA/H7N9-4M in the encapsidase conformation (FluPolA/H7N9-4M(E)), extracted from the replication complex and aligned on FluPolA/H7N9(T) PB1 subunit. FluPolA/H7N9-4M(E) core is in light grey, shown in transparency, PA ENDO(E) in light green, PB2 midlink(E) in salmon, PB2 CBD(E) in orange, PB2 627(E) in light pink, PB2 NLS(E) in brown. PB1-C(E) and PB2-N(E) are flexible. Specific interactions within FluPolA/H7N9-4M(E) are annotated with dotted black circles, and refer to panels **(F)** and **(G)**. **(D)** PA-endonuclease (ENDO) conformation comparison between FluPolA/H7N9(T) and FluPolA/H7N9-4M(R). ENDOs are displayed as transparent surface, coloured from the N-terminus to the C-terminus from dark to light green. ENDO(R) rotates and interacts with PB2 NLS(R), represented as cartoon and coloured in beige. **(E)** Cartoon representation of PB2(R) C-terminal domains and PB1(R) palm domain. PB2(R) C-terminal domains are coloured as in **(B)**. PB2 CBD(R) interacts with PB1 palm domain, in light grey. Catalytic aspartic acids are shown with atoms as spheres, circled with a dotted line. **(F)** Close-up view of PA ENDO(E) flexible insertion (51-72) interacting with PB2 CBD(E). Domains are coloured as in **(C)**. PA ENDO(E) residues 67-72 are flexible, represented as a dotted line. Ionic and hydrogen bonds are shown as grey dotted lines. **(G)** Close-up view of the interaction between PB2-N(E) and PB2 midlink(E). Domains are coloured as in **(C)**. Interacting residues are displayed.

**Figure S3.**
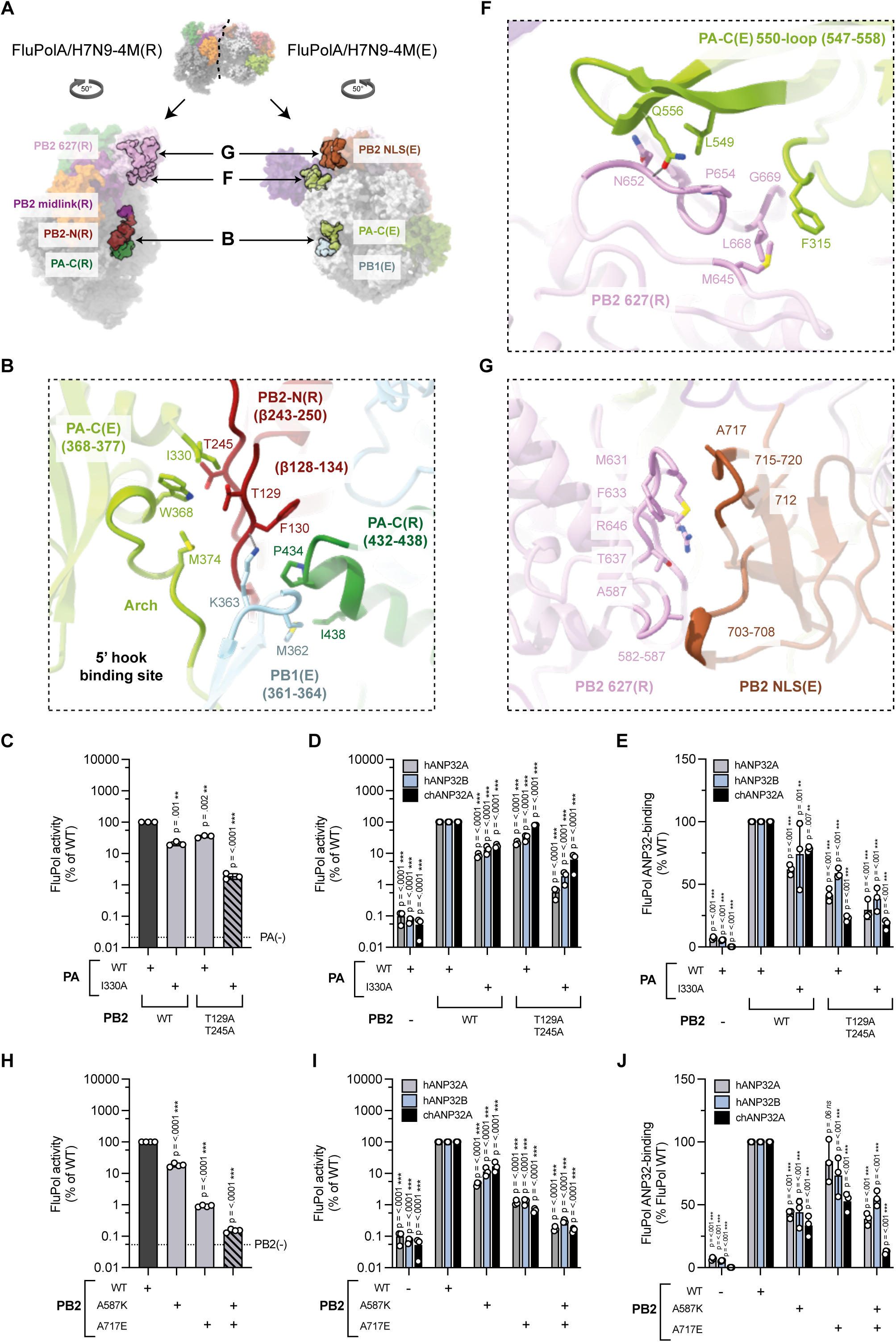
Interface between FluPolA/H7N9-4M(R) and FluPolA/H7N9-4M(E). **(A)** Overview of the interacting domains between FluPolA/H7N9-4M(R) and FluPolA/H7N9-4M(E). Both replicase and encapsidase moiety were split and rotated of 50 degree. Interacting surface is shown as non-transparent. Most domains are coloured as in Figure 2, with PA(R) in dark green, PB2-N(R) in red. For the three main interfaces, a close-up view is shown in panels **(B)**, **(F)**, **(G)**. **(B)** FluPolA/H7N9-4M PA-C(R), PB2-N(R) and FluPolA/H7N9-4M PA(E) arch interaction. Domains are coloured as in **(A).** Interacting residues are displayed, shown as non-transparent. Ionic and hydrogen bonds are shown as grey dotted lines. **(C)** Mutational analysis of the PB2-N(R) - PA(E) arch interaction shown in **(B)**. Cell-based assay of WSN FluPol activity for the indicated PA and PB2 mutants and combinations thereof. HEK-293T cells were co-transfected with plasmids encoding PB2, PB1, PA, NP with a model vRNA encoding the Firefly luciferase. Luminescence was normalised to a transfection control and is represented as a percentage of FluPol WT (mean ± SD, n=3, **p < 0.002, ***p < 0.001, one-way ANOVA; Dunnett’s multiple comparisons test). **(D)** Mutational analysis of the PB2-N(R) - PA(E) arch interaction shown in **(B)**. Cell-based assay of WSN FluPol activity for the indicated PA and PB2 mutants and combinations thereof. HEK-293T in which hANP32A and hANP32B were knocked out were transfected as in **(C)** and transiently complemented by co-transfection of plasmids encoding hANP32A, hANP32B or chANP32A. Luminescence was normalised to a transfection control and is represented as a percentage of FluPol WT (mean ± SD, n=3, ***p < 0.001, two-way ANOVA; Dunnett’s multiple comparisons test). **(E)** Mutational analysis of the PB2-N(R) - PA(E) arch interaction as shown in **(B)**. Cell-based assay of WSN FluPol binding to ANP32 for the indicated PA and PB2 mutants and combinations thereof. HEK-293T cells were co-transfected with plasmids encoding PB2, PA, PB1-luc1 and either hANP32A-luc2, hANP32B-luc2 or chANP32A-luc2. Luminescence signals due to luciferase reconstitution are represented as a percentage of FluPol WT (mean ± SD, n=3, **p < 0.002, ***p < 0.001, two-way ANOVA; Dunnett’s multiple comparisons test). **(F)** FluPolA/H7N9-4M PB2 627(R) C-terminal β-sheet and FluPolA/H7N9-4M PA-C 550-loop(E) interaction. Domains are coloured as in **(A)**. Interacting residues are displayed, shown as non-transparent. Ionic and hydrogen bonds are shown as grey dotted lines. **(G)** FluPolA/H7N9-4M PB2 627(R) and FluPolA/H7N9-4M PB2 NLS(E) interaction. Domains are coloured as in **(A)**. Most of interacting residues are displayed, shown as non-transparent. **(H)** Mutational analysis of the PB2-627(R) - PB2-NLS(E) interaction as shown in **(G)**. Cell-based assay of WSN FluPol activity for the indicated PB2 mutants and combinations thereof as described in **(C)** (mean ± SD, n=4, ***p < 0.001, one-way ANOVA; Dunnett’s multiple comparisons test). **(I)** Mutational analysis of the PB2-627(R) - PB2-NLS(E) interaction as shown in **(G)**. Cell-based assay of WSN FluPol activity for the indicated PB2 mutants and combinations thereof as described in **(D)** (mean ± SD, n=3, ***p < 0.001, two-way ANOVA; Dunnett’s multiple comparisons test). **(J)** Mutational analysis of the PB2-627(R) - PB2-NLS(E) interaction as shown in **(G)**. Cell-based assay of WSN FluPol binding to ANP32 for the indicated PB2 mutants and combinations thereof as described in **(E)** (mean ± SD, n=3, ***p < 0.001, two-way ANOVA; Dunnett’s multiple comparisons test).

**Figure S4.**
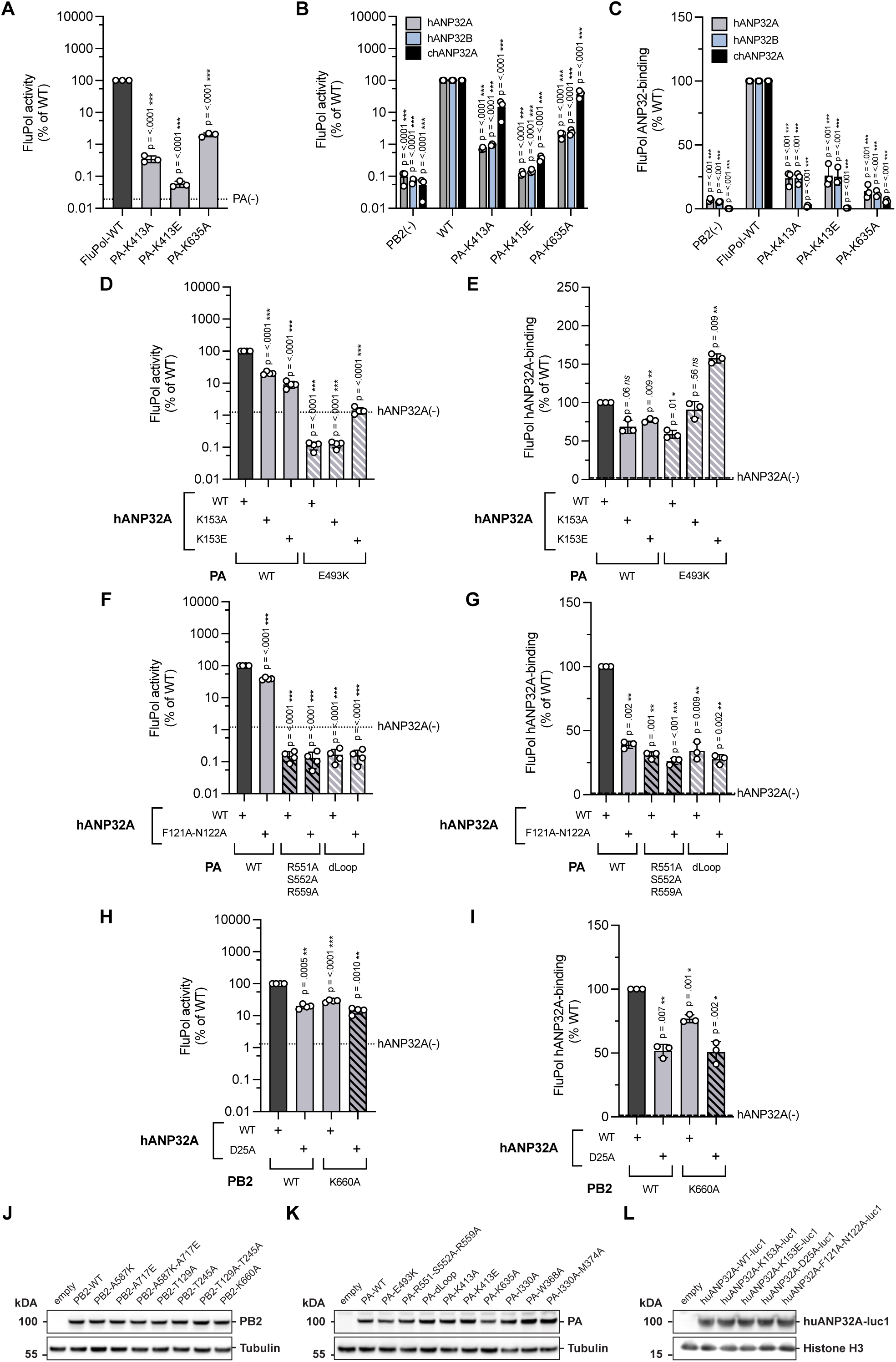
Interface between FluPolA/H7N9-4M and hANP32A. **(A)** Mutational analysis of the PA(E)-hANP32A interaction shown in Fig. 2C. Cell-based assay of WSN FluPol activity for the indicated PA mutants. HEK-293T WT cells were co-transfected with plasmids encoding PB2, PB1, PA, NP with a model vRNA encoding the Firefly luciferase. Luminescence was normalised to a transfection control and is represented as a percentage of PA-WT (mean ± SD, n=3, ***p < 0.001, one-way ANOVA; Dunnett’s multiple comparisons test). **(B)** Mutational analysis of the PA(E)-hANP32A interaction shown in Fig. 2C. Cell-based assay of WSN FluPol activity for the indicated PA mutants. HEK-293T in which hANP32A and hANP32B were knocked out were transfected as in **(A)** and transiently complemented by co-transfection of plasmids encoding hANP32A, hANP32B or chANP32A. Luminescence was normalised to a transfection control and is represented as a percentage of FluPol WT (mean ± SD, n=3, ***p < 0.001, two-way ANOVA; Dunnett’s multiple comparisons test). **(C)** Mutational analysis of the PA(E)-hANP32A interaction shown in Fig. 2C. Cell-based assay of WSN FluPol binding to ANP32 for the indicated PA mutants. HEK-293T cells were co-transfected with plasmids encoding PB2, PA, PB1-luc1 and either hANP32A-luc2, hANP32B-luc2 or chANP32A-luc2. Luminescence signals due to luciferase reconstitution are represented as a percentage of FluPol WT (mean ± SD, n=3, ***p < 0.001, two-way ANOVA; Dunnett’s multiple comparisons test). **(D)** Mutational analysis of the PA(E)-hANP32A interaction shown in Fig. 2D. Cell-based assay of WSN FluPol activity for the indicated PA and hANP32A mutants. HEK-293T in which hANP32A and hANP32B were knocked out were transfected as in **(A)** and transiently complemented by co-transfection of plasmids encoding hANP32A-WT or the indicated hANP32A mutants. Luminescence was normalised to a transfection control and is represented as a percentage of WT (mean ± SD, n=4, ***p < 0.001, one-way ANOVA; Dunnett’s multiple comparisons test). **(E)** Mutational analysis of the PA(E)-hANP32A interaction shown in Fig. 2D. Cell-based assay or WSN FluPol binding to ANP32 for the indicated PA and hANP32A mutants as described in **(C)** (mean ± SD, n=3, *< 0.033, **p < 0.002, one-way ANOVA; Dunnett’s multiple comparisons test). **(F)** Mutational analysis of the PA(E)-hANP32A interaction shown in Fig. 2E. Cell-based assay of WSN FluPol activity for the indicated PA and hANP32A mutants as described in **(D)** (mean ± SD, n=4, ***p < 0.001, one-way ANOVA; Dunnett’s multiple comparisons test). **(G)** Mutational analysis of the PA(E)-hANP32A interaction shown in Fig. 2E. Cell-based assay of WSN FluPol binding to ANP32 for the indicated PA and hANP32A mutants as described in **(C)** (mean ± SD, n=3, **p < 0.002, ***p < 0.001, one-way ANOVA; Dunnett’s multiple comparisons test). **(H)** Mutational analysis of the PB2(R)-hANP32A interaction shown in Fig. 2F. Cell-based assay of WSN FluPol activity for the indicated PB2 and hANP32A mutants as described in **(D)** (mean ± SD, n=4, **p < 0.002, ***p < 0.001, one-way ANOVA; Dunnett’s multiple comparisons test). **(I)** Mutational analysis of the PB2(R)-hANP32A interaction shown in Fig. 2F. Cell-based WSN FluPol ANP32-binding assays of the indicated PB2 and hANP32A mutants as described in **(C)** (mean ± SD, n=3, *< 0.033, **p < 0.002, one-way ANOVA; Dunnett’s multiple comparisons test). **(J-K)** HEK-293T cells were co-transfected with expression plasmids for WSN PB1, PB2 and PA with the indicated PB2 **(J)** or PA **(K)** mutations. Cell lysates were analysed by western blot using antibodies specific for PB2, PA and tubulin. Uncropped gels are provided as a source data file. **(L)** HEK-293T cells were transfected with expression plasmids for hANP32A-luc1 with the indicated mutations. Cell lysates were analysed by western blot using antibodies specific for Gaussia luciferase and Histone H3. Uncropped gels are provided as a source data file.

**Figure S5.**
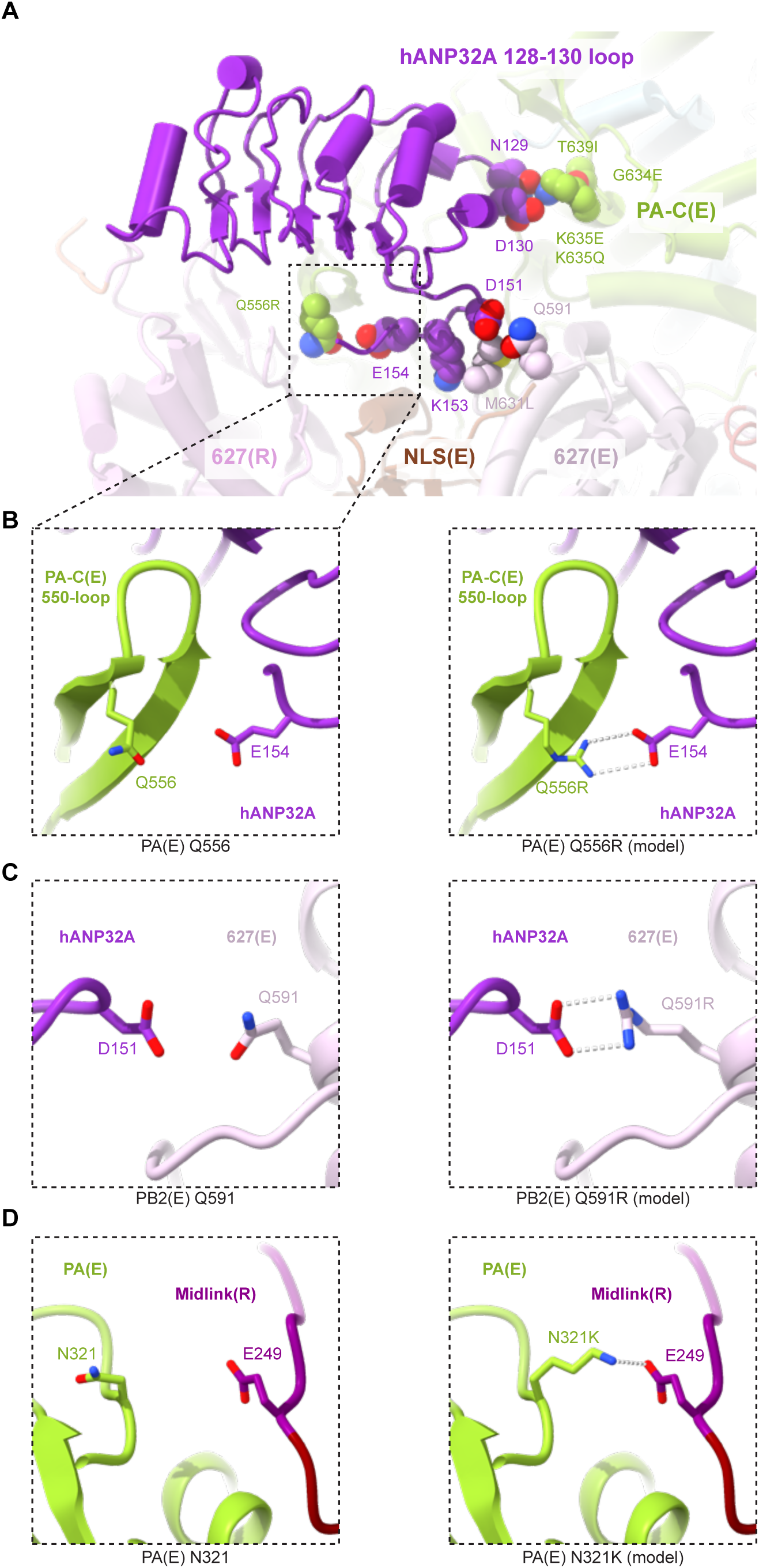
Human adaptive mutations mapped onto the FluPolA/H7N9-4M replication complex. **(A)** Overview of hANP32A interaction with FluPolA/H7N9-4M(R) and FluPolA/H7N9-4M(E). Domains are coloured as in Figure 2. FluPolA/H7N9-4M adaptive mutations are annotated (PA Q556R, G634E K635E/Q, T639I and PB2 I570L, Q591R, M631L). Corresponding residues of the FluPolA/H7N9-4M replication complex structure are displayed. Atoms are shown as spheres. **(B)** Close-up view of PA-C(E) showing the effect of the Q556R mutation. Left: PA-C(E) Q556 residue as built in the FluPolA/H7N9-4M replication complex structure. Right: Modelled PA-C(E) Q556R mutation is likely to make a salt-bridge with hANP32A E154. Ionic bonds are shown as grey dotted lines. **(C)** Close-up view of PB2(E) showing the effect of the Q591R mutation. Left: PB2(E) Q591 residue as built in the FluPolA/H7N9-4M replication complex structure. Right: Modelled PB2(E) Q591R mutation is likely to make a salt-bridge with hANP32A D151. Ionic bonds are shown as grey dotted lines. **(D)** Close-up view of PA(E) showing the effect of the N321K mutation. Left: PA(E) N321 residue as built in the FluPolA/H7N9-4M replication complex structure. Right: Modelled PA(E) N321K mutation is likely to reinforce the replicase-encapsidase interface, by interacting with PB2 midlink(R) E249. Ionic bonds are shown as grey dotted lines.

**Figure S6.**
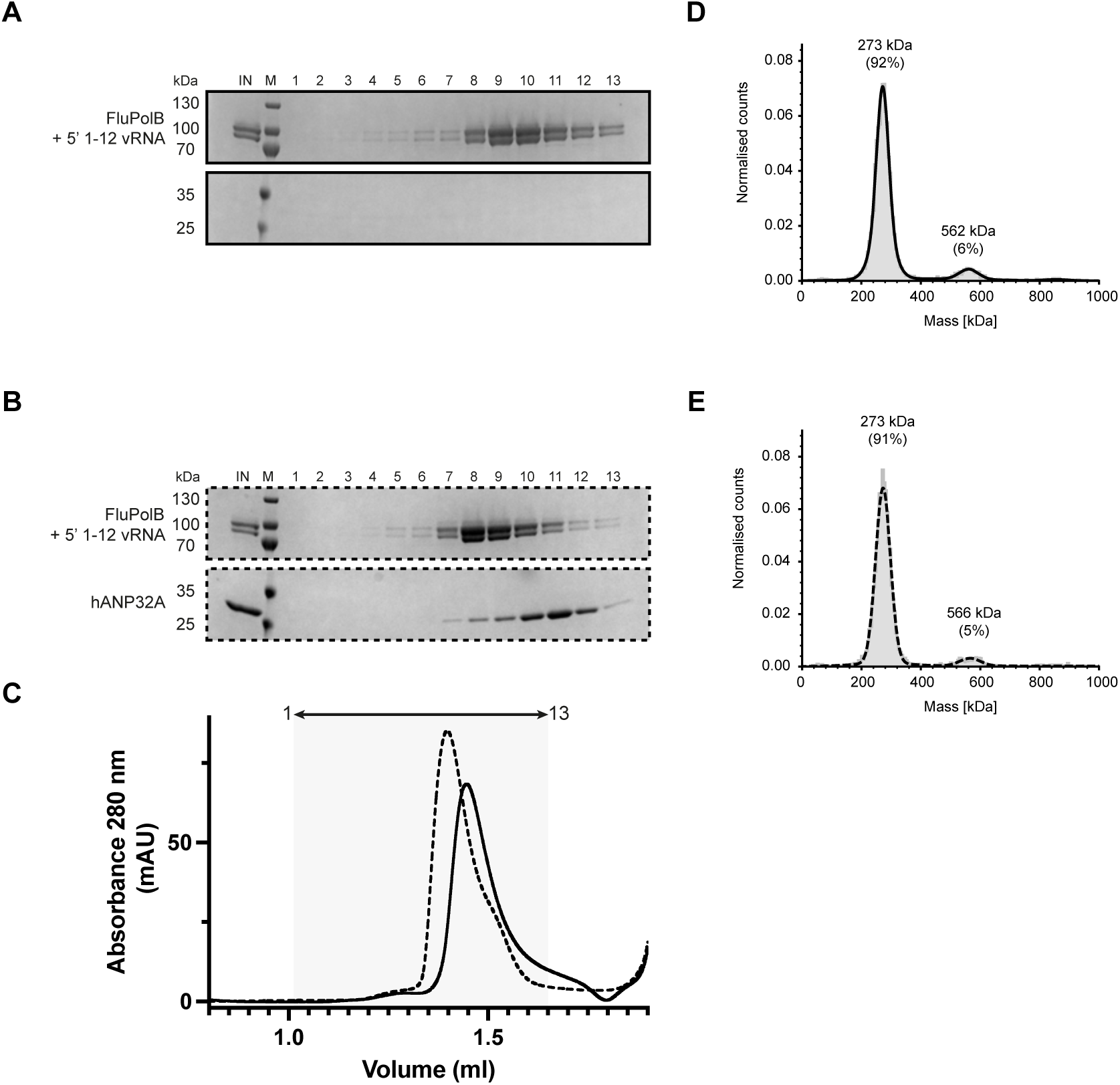
5′ vRNA end binding dissociates FluPolB dimer. **(A)** SDS-PAGE analysis of FluPolB bound to the 5′ vRNA end (nts 1-12) at 150 mM NaCl. The molecular ladder (M) in kDa and FluPolB heterotrimer are indicated on the left of the gel. “IN” corresponds to the input. **(B)** SDS-PAGE analysis of FluPolB bound to the 5′ vRNA end (1-12) with excess of hANP32A at 150 mM NaCl. The molecular ladder (M) in kDa, FluPolB heterotrimer and hANP32A are indicated on the left of the gel. “IN” corresponds to the input. **(C)** Superposition of size exclusion chromatography profiles of FluPolB bound to 5′ vRNA end (1-12) (solid line), and with hANP32A (dotted line), at 150 mM NaCl. The relative absorbance at 280 nm (mAU) is on the y-axis. The elution volume (ml) is on the x-axis, graduated every 50 µl. SDS-PAGE fractions 1 to 13 corresponds to the elution volume 1.0 ml - 1.65 ml. **(D)** Mass photometry analysis of FluPolB bound to the 5′ vRNA end (1-12) at 150 mM NaCl. The determined masses in kDa of the main species are indicated. **(E)** Mass photometry analysis of FluPolB bound to the 5′ vRNA end (1-12) with excess of hANP32A at 150 mM NaCl. The determined masses in kDa of the main species are indicated.

**Figure S7.**
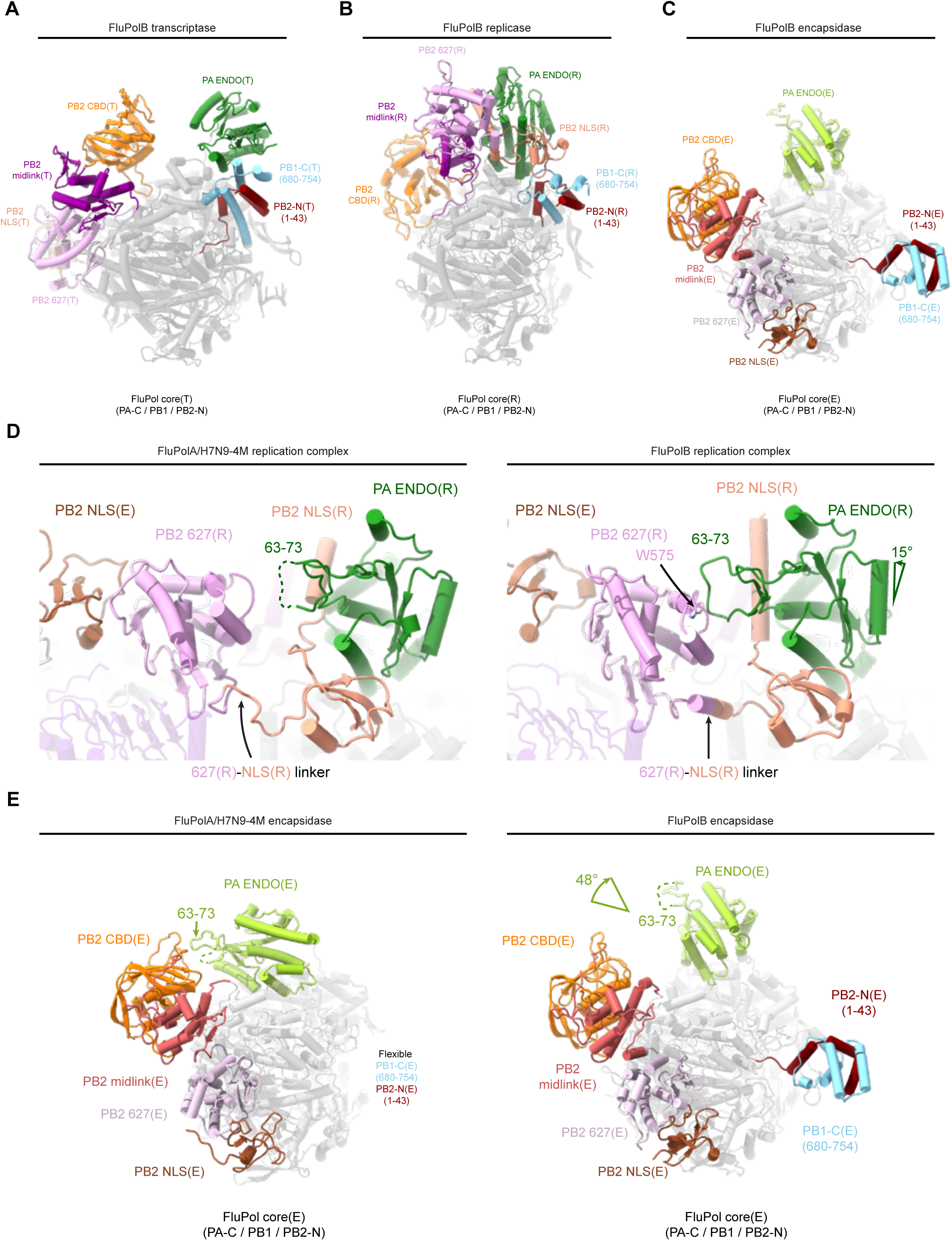
Structural comparison of FluPolB transcriptase, replicase, encapsidase conformations and between FluPolA/H7N9-4M, FluPolB replication complexes. **(A)** Cartoon representation of FluPolB in a transcriptase conformation (FluPolB(T)) (PDB 4WSA). FluPolB(T) core is dark grey, shown in transparency, PA ENDO(T) in dark green, PB1-C(T) in blue, PB2-N(T) in red, PB2 midlink(T) in magenta, PB2 CBD(T) in orange, PB2 627(T) in pink, PB2 NLS(T) in beige. **(B)** Cartoon representation of FluPolB in a replicase conformation (FluPolB(R)), extracted from the replication complex and aligned on FluPolB(T) PB1 subunit. FluPolB(R) core is in dark grey, shown in transparency, PA ENDO(R) in dark green, PB1-C(R) in blue, PB2-N(R) in red, PB2 midlink(R) in magenta, PB2 CBD(R) in orange, PB2 627(R) in pink, PB2 NLS(R) in beige. **(C)** Cartoon representation of FluPolB in an encapsidase conformation (FluPolB(E)), extracted from the replication complex and aligned on FluPolB(T) PB1 subunit. FluPolB(E) core is in light grey, shown in transparency, PA ENDO(E) in light green, PB1-C(R) in blue, PB2-N(R) in red, PB2 midlink(E) in salmon, PB2 CBD(E) in orange, PB2 627(E) in light pink, PB2 NLS(E) in brown. PB1-C(E) and PB2-N(E) helical bundle swung away, interacting with FluPolB(R) (shown in **Figure S8F**). **(D)** Structural comparison between FluPolA/H7N9-4M and FluPolB replication complexes. Domains are coloured as in **(B-C)**. FluPolA/H7N9-4M PA ENDO(R) 63-73 loop is flexible, shown as a dotted line. FluPolB(R) equivalent interacts with FluPolB PB2 627(R), next to W575, displayed and indicated. FluPolB PA ENDO(R) compared to FluPolA/H7N9-4M PA ENDO(R) undergoes a 15 degree rotation, indicated with an arrow. PB2 627(R)/NLS(R) linkers are indicated, taking up an α-helical conformation in FluPolB replication complex. **(E)** Structural comparison between FluPolA/H7N9-4M(E) and FluPolB(E). Domains are coloured as in **(C).** FluPolA/H7N9-4M PA ENDO(E) 63-73 insertion interacts with PB2 CBD(E) (as seen in **Figure S2F**). FluPolB PA ENDO(E), compared to FluPolA/H7N9-4M PA ENDO(E), undergoes a 48 degree rotation, indicated an arrow. FluPolB PA ENDO(E) 63-73 loop is flexible, represented as a dotted line.

**Figure S8.**
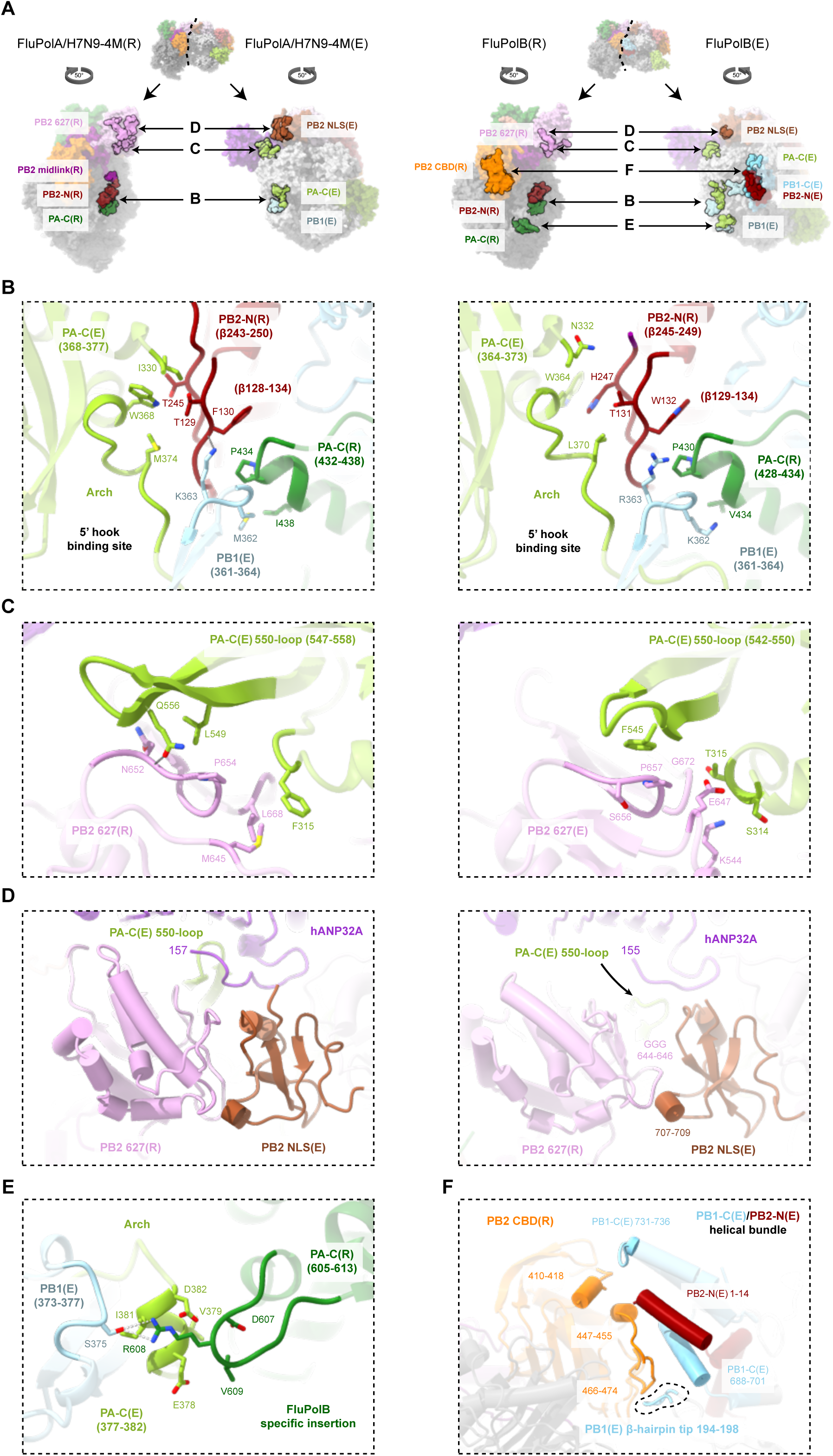
Structural comparison between FluPolA/H7N9-4M and FluPolB replication complexes interfaces. **(A)** Overview of the interacting domains between FluPols(R) and FluPols(E). Both replicase and encapsidase moiety were split and rotated by 50 degree. Interacting surface is shown as non-transparent. Most domains are coloured as in Figure 2, with PA(R) in dark green, PB2-N(R) in red. For the three main conserved interfaces between FluPolA/H7N9-4M and FluPolB replication complexes, a close-up view is shown in panels **(B)**, **(C)**, **(D)**. **(B)** PA-C(R), PB2-N(R) and PA(E) arch interaction. Domains are coloured as in **(A)**. Interacting residues are displayed, shown as non-transparent. Ionic and hydrogen bonds are shown as grey dotted lines. **(C)** PB2 627(R) C-terminal β-sheet and PA-C 550-loop(E) interaction. Domains are coloured as in **(A)**. Interacting residues are displayed, shown as non-transparent. Ionic and hydrogen bonds are shown as grey dotted lines. **(D)** PB2 627(R) and PB2 NLS(E) interaction. Domains are coloured as in **(A)**. Interacting residues are displayed, shown as non-transparent. **(E)** FluPolB PA-C(R), PA-C(E) and PB1(E) specific interface. FluPolB PA-C(R) specific insertion 605-613 interacts with PA-C(E) 377-382 and PB1(E) 373-377. Domains are coloured as in **(A)**. Interacting residues are displayed, shown as non-transparent. **(F)** FluPolB PB1-C(E)/PB2-N(E) helical bundle interacts with PB2 CBD(R). Interacting Domains are coloured as in **(A)** and shown as non-transparent.

**Supplemental information 1.**
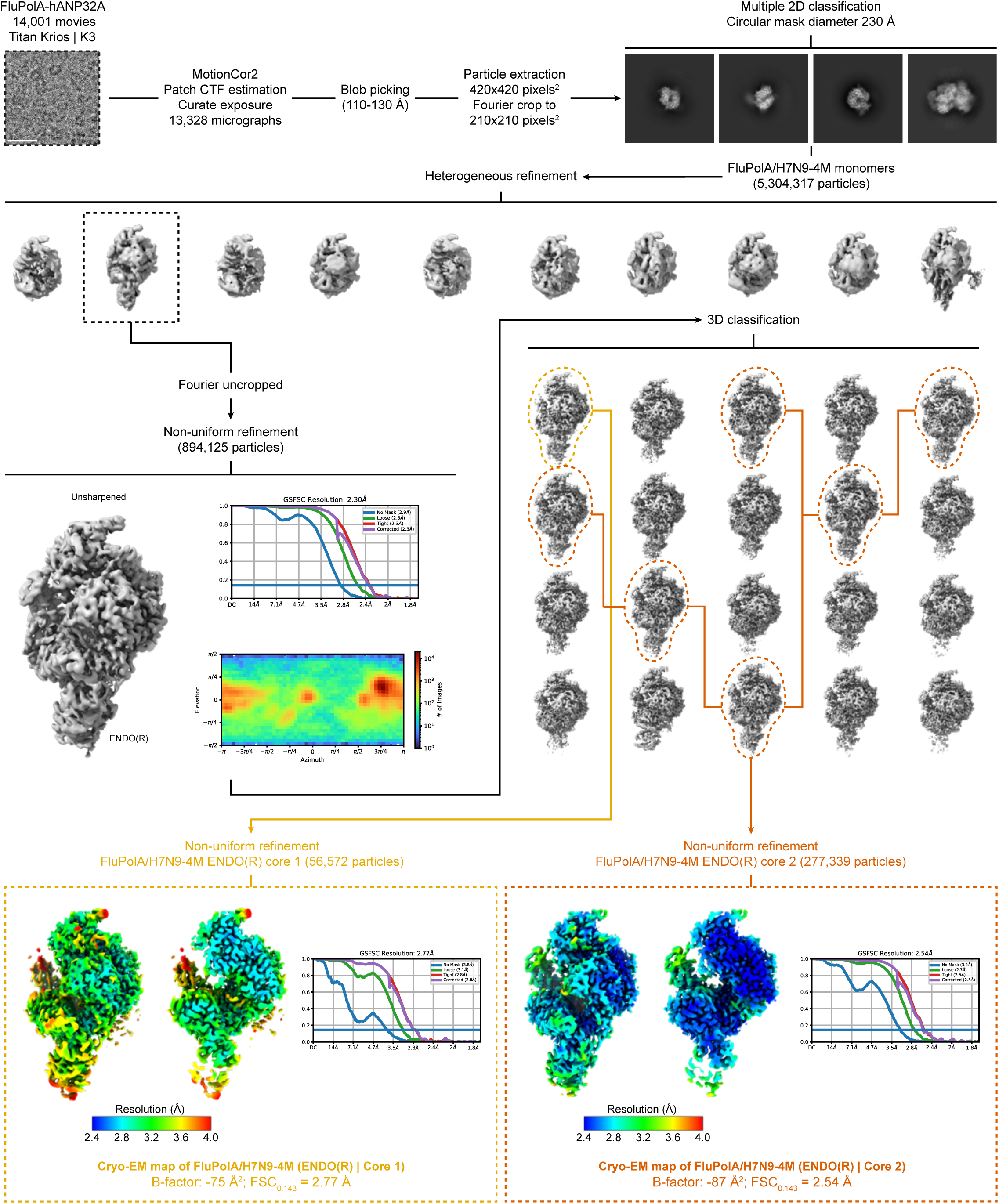
Cryo-EM image processing strategy applied to obtain FluPolA/H7N9-4M ENDO(R) core1 and core2 structures. Schematics of the image processing strategy used with the data collected on a TEM Titan Krios equipped with a Gatan K3 direct electron detector mounted on a Gatan Bioquantum energy filter. Representative cropped micrograph, 2D class averages and 3D classes are displayed. Full and cutaway views of each local resolution filtered EM maps are shown. Fourier shell correlation curves (FSC) are displayed. Scale bar = 200 Å.

**Supplemental information 2.**
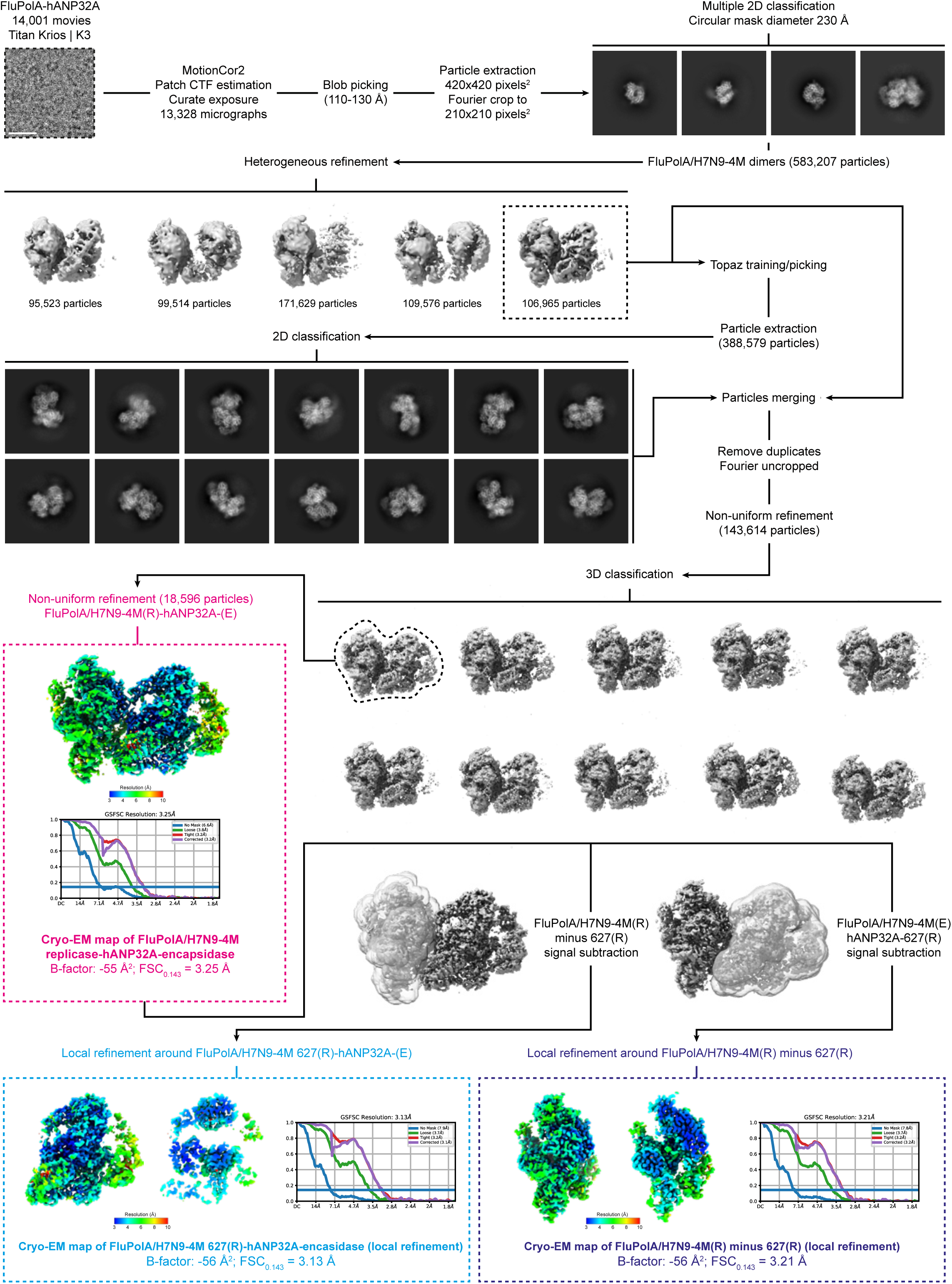
Cryo-EM image processing strategy applied to obtain FluPolA/H7N9-4M replication complex structures. Schematics of the image processing strategy used with the data collected on a TEM Titan Krios equipped with a Gatan K3 direct electron detector mounted on a Gatan Bioquantum energy filter. Representative cropped micrograph, 2D class averages and 3D classes are displayed. Full and cutaway views of each DeepEMhancer (Sanchez-Garcia et al., 2021) filtered EM maps are shown. Fourier shell correlation curves (FSC) are displayed. Scale bar = 200 Å.

**Supplemental information 3.**
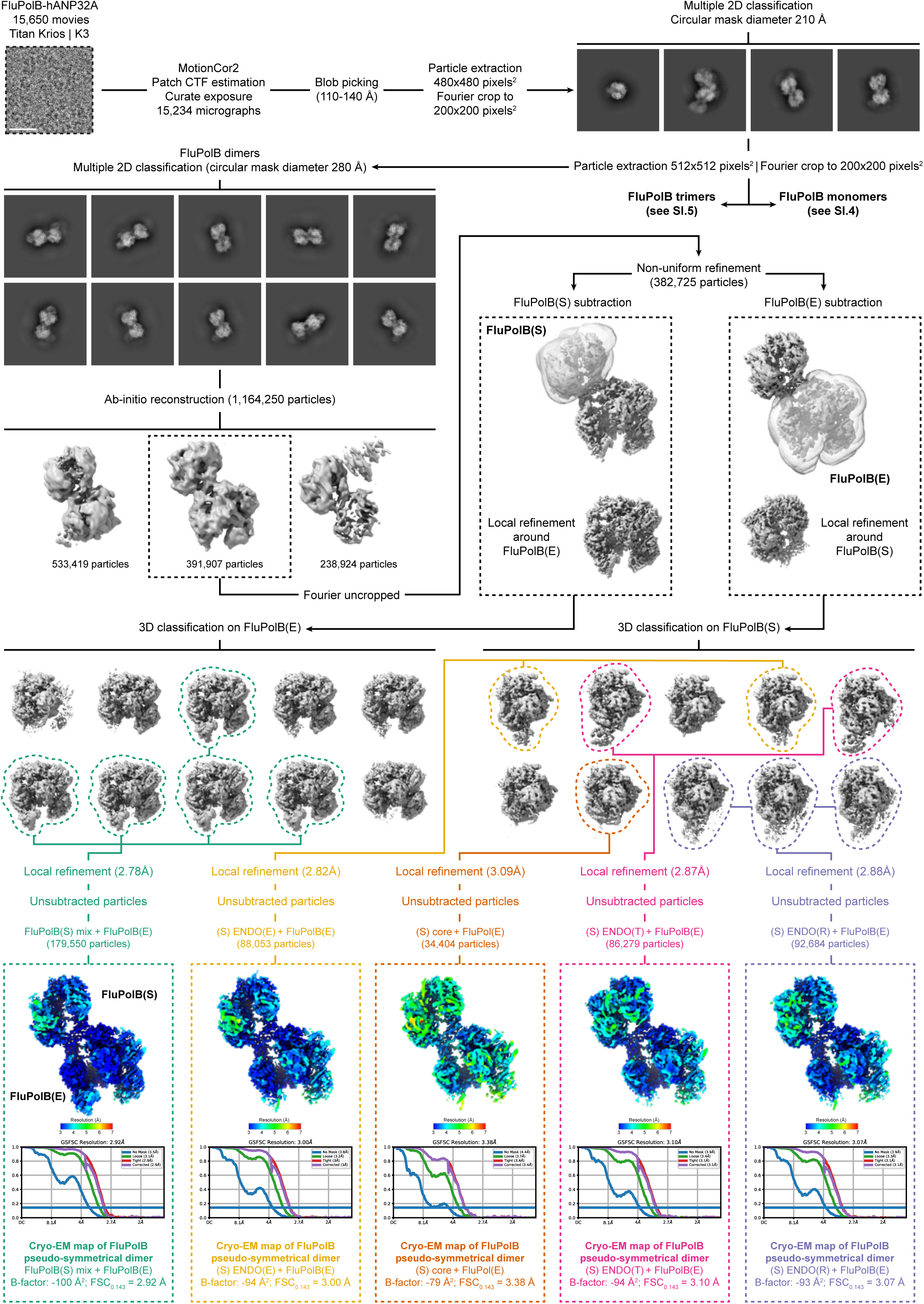
Cryo-EM image processing strategy applied to obtain FluPolB pseudo-symmetrical dimer structures, with one moiety being an encapsidase. Schematics of the image processing strategy used with the data collected on a TEM Titan Krios equipped with a Gatan K3 direct electron detector mounted on a Gatan Bioquantum energy filter. Representative cropped micrograph, 2D class averages and 3D classes are displayed. Full and cutaway views of each local resolution filtered EM maps are shown. Fourier shell correlation curves (FSC) are displayed. Scale bar = 200 Å.

**Supplemental information 4.**
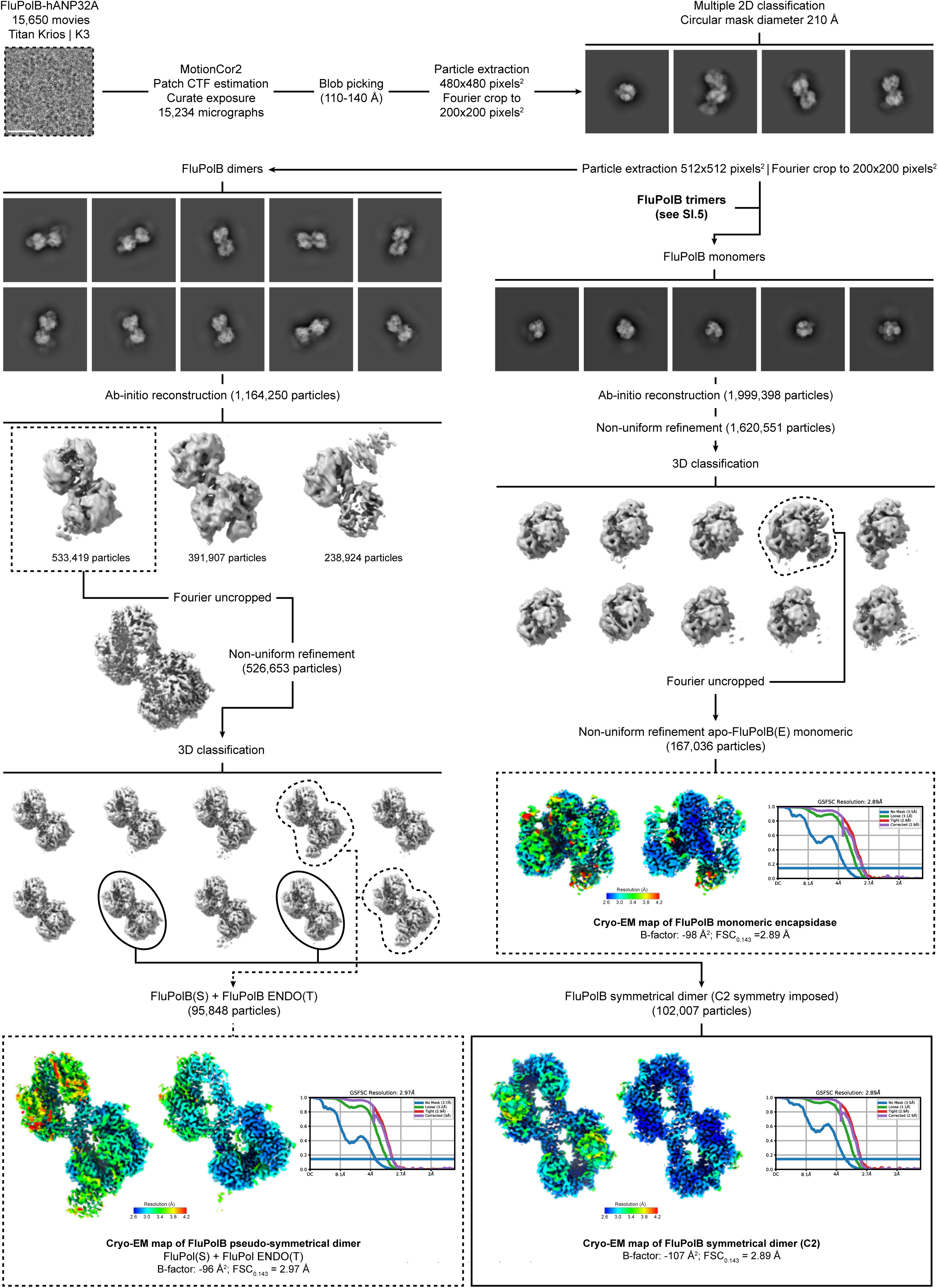
Cryo-EM image processing strategy applied to obtain FluPolB (pseudo-)symmetrical dimer structures and monomeric apo-FluPolB encapsidase. Schematics of the image processing strategy used with the data collected on a TEM Titan Krios equipped with a Gatan K3 direct electron detector mounted on a Gatan Bioquantum energy filter. Representative cropped micrograph, 2D class averages and 3D classes are displayed. Full and cutaway views of each local resolution filtered EM maps are shown. Fourier shell correlation curves (FSC) are displayed. Scale bar = 200 Å.

**Supplementary figure 5.**
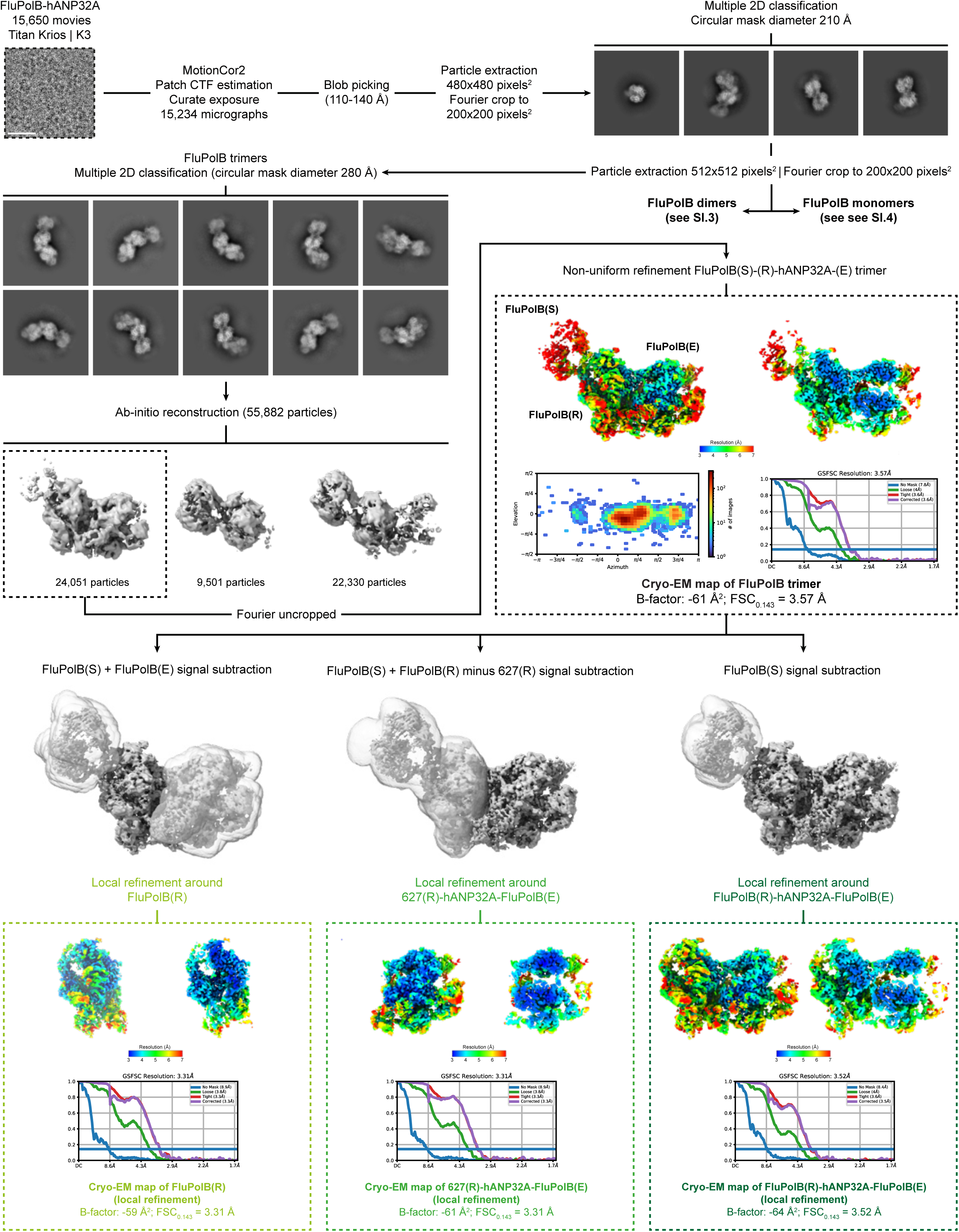
Cryo-EM image processing strategy applied to obtain FluPolB replication complex structures. Schematics of the image processing strategy used with the data collected on a TEM Titan Krios equipped with a Gatan K3 direct electron detector mounted on a Gatan Bioquantum energy filter. Representative cropped micrograph, 2D class averages and 3D classes are displayed. Full and cutaway views of each DeepEMhancer (Sanchez-Garcia et al., 2021) filtered EM maps are shown. Fourier shell correlation curves (FSC) are displayed. Scale bar = 200 Å.

**Supplemental information 6.**
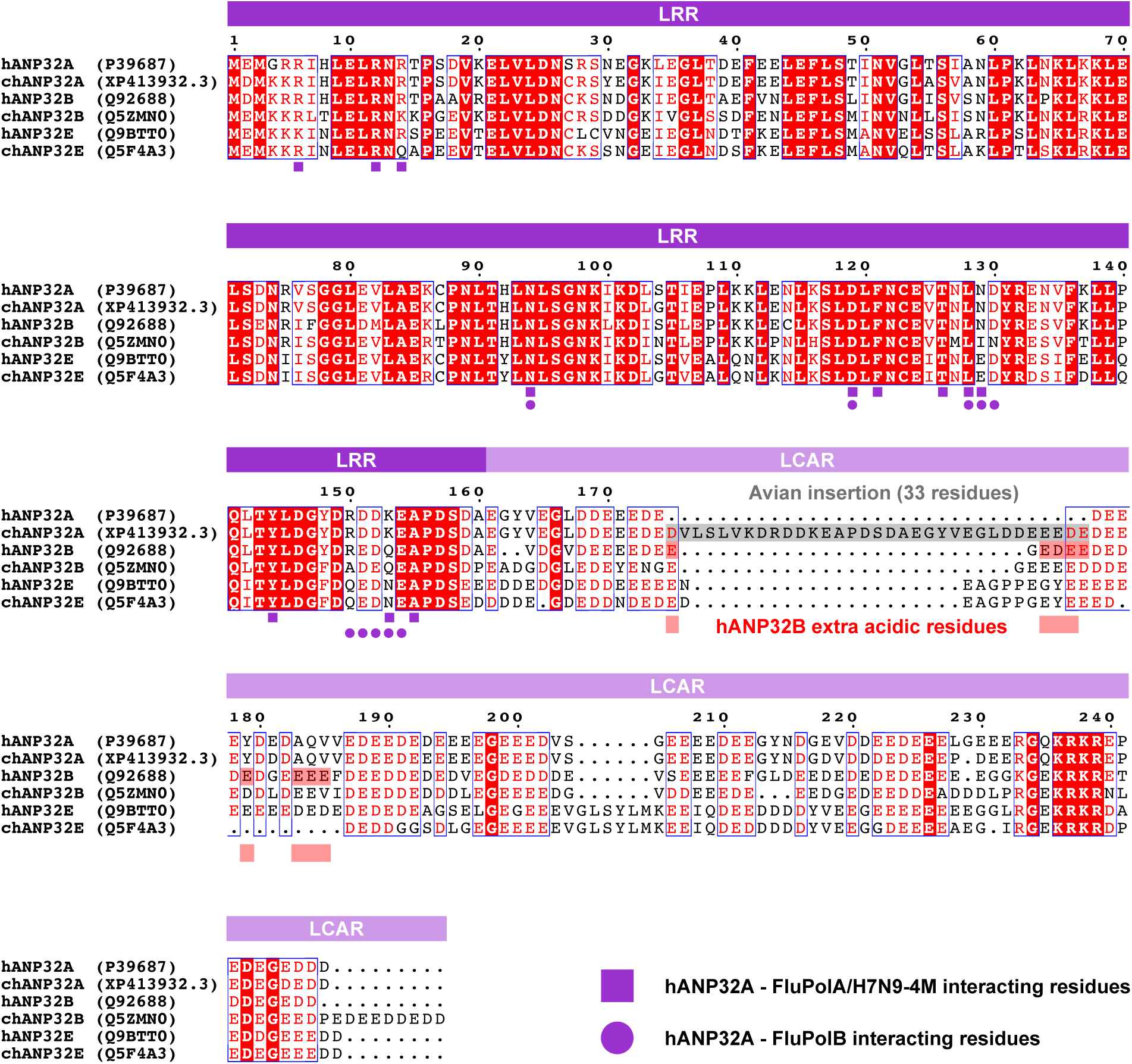
Multiple sequence alignment of human and chicken ANP32A, B and E. Human (h), chicken (ch), ANP32A/B/E UniProt numbers are indicated. Both Leucine Rich Repeat (LRR) and Low Complexity Acidic Region (LCAR) are indicated on top of the aligned sequences, respectively in dark and light purple. hANP32A interacting residues with FluPolA/H7N9-4M and FluPolB are indicated below aligned sequences, with respectively purple squares, or circles. The specific avian insertion of 33 residues is highlighted with a grey rectangle. hANP32B extra acidic residues are highlighted with red rectangles.

**Supplemental information 7.**
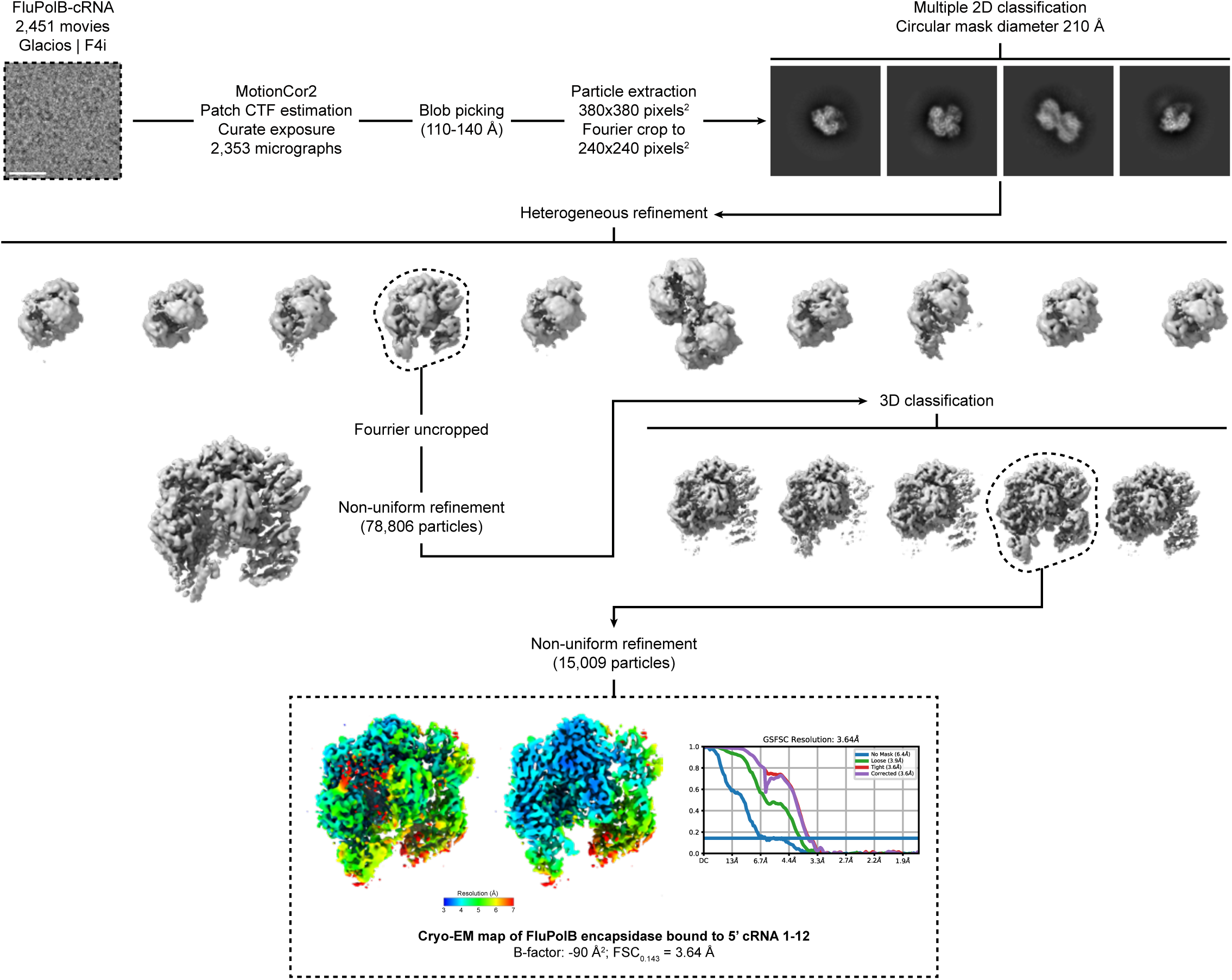
Cryo-EM image processing strategy applied to obtain FluPolB encapsidase bound to 5’ cRNA 1-12. Schematics of the image processing strategy used with the data collected on a TEM Glacios equipped with a Falcon4i direct electron detector mounted on a SelectrisX energy filter. Representative cropped micrograph, 2D class averages and 3D classes are displayed. Full and cutaway views of each local resolution filtered EM maps are shown. Fourier shell correlation curves (FSC) are displayed. Scale bar = 200 Å.

